# Age-specific survivorship and fecundity shape genetic diversity in marine fishes

**DOI:** 10.1101/2020.12.18.423459

**Authors:** Pierre Barry, Thomas Broquet, Pierre-Alexandre Gagnaire

**Affiliations:** ISEM, Univ Montpellier, CNRS, EPHE, IRD, Montpellier, France; CNRS & Sorbonne Université, UMR 7144, Station Biologique de Roscoff, 29680 Roscoff, France

**Keywords:** genetic diversity, life tables, adult lifespan, variance in reproductive success, marine fishes

## Abstract

Genetic diversity varies among species due to a range of eco-evolutionary processes that are not fully understood. The neutral theory predicts that the amount of variation in the genome sequence between different individuals of the same species should increase with its effective population size (*N*_*e*_). In real populations, multiple factors that modulate the variance in reproductive success among individuals cause *N*_*e*_ to differ from the total number of individuals (*N*). Among these, age-specific mortality and fecundity rates are known to have a direct impact on the 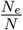 ratio. However, the extent to which vital rates account for differences in genetic diversity among species remains unknown. Here, we addressed this question by comparing genome-wide genetic diversity across 16 marine fish species with similar geographic distributions but contrasted lifespan and age-specific survivorship and fecundity curves. We sequenced the whole genome of 300 individuals to high coverage and assessed their genome-wide heterozygosity with a reference-free approach. Genetic diversity varied from 0.2 to 1.4% among species, and showed a negative correlation with adult lifespan, with a large negative effect (*slope* = − 0.089 per additional year of lifespan) that was further increased when brooding species providing intense parental care were removed from the dataset (*slope* = −0.129 per additional year of lifespan). Using published vital rates for each species, we showed that the 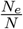 ratio resulting simply from life tables parameters can predict the observed differences in genetic diversity among species. Using simulations, we further found that the extent of reduction in 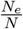 with increasing adult lifespan is particularly strong under Type III survivorship curves (high juvenile and low adult mortality) and increasing fecundity with age, a typical characteristic of marine fishes. Our study highlights the importance of vital rates as key determinants of species genetic diversity levels in nature.

**Author Summary:** Understanding how and why genetic diversity varies across species has important implications for evolutionary and conservation biology. Although genomics has vastly improved our ability to document intraspecific DNA sequence variation at the genome level, the range and determinants of genetic diversity remain partially understood. At a broad taxonomic scale in eukaryotes, the main determinants of diversity are reproductive strategies distributed along a trade-off between the quantity and the size of offspring, which likely affect the long-term effective population size. Long-lived species also tend to show lower genetic diversity, a result which has however not been reported by comparative studies of genetic diversity at lower taxonomic scales. Here, we compared genetic diversity across 16 European marine fish species showing marked differences in longevity. Adult lifespan was the best predictor of genetic diversity, with genome-wide average heterozygosity ranging from 0.2% in the black anglerfish (*L. budegassa*) to 1.4% in the European pilchard (*S. pilchardus*). Using life tables summarizing age-specific mortality and fecundity rates for each species, we showed that the variance in lifetime reproductive success resulting from age structure, iteroparity and overlapping generations can predict the range of observed differences in genetic diversity among marine fish species. We then used computer simulations to explore how combinations of vital rates characterizing different life histories affect the relationship between adult lifespan and genetic diversity. We found that marine fishes that display high juvenile but low adult mortality, and increasing fecundity with age, are typically expected to show reduced genetic diversity with increased adult lifespan. However, the impact of adult lifespan vanished using bird and mammal-like vital rates. Our study shows that variance in lifetime reproductive success can have a major impact on species genetic diversity and explains why this effect varies widely across taxonomic groups.

## Introduction

Genetic diversity, the substrate for evolutionary change, is a key parameter for species adaptability and vulnerability in conservation and management strategies (Frankham, 1995; Lande, 1995). Understanding the determinants of species’ genetic diversity has been, however, a long-standing puzzle in evolutionary biology (Lewontin, 1974). Advances in DNA sequencing technologies have allowed to describe the range of genetic diversity levels across eukaryote species and identify the main evolutionary processes governing that variation (Leffler et al., 2012; Romiguier et al., 2014). Yet, the extent and reasons for which life history traits, and in particular reproductive strategies, influence genetic diversity remain to be clarified (Ellegren and Galtier, 2016).

The neutral theory provides a quantitative prediction for the amount of genetic variation at neutral sites (Kimura, 1983). Assuming equilibrium between the introduction of new variants by mutations occurring at rate *µ*, and their removal by genetic drift at a rate inversely proportional to the effective population size *N*_*e*_, the amount of genetic diversity (*θ*) of a stable randomly mating population is equal to 4*N*_*e*_*µ* (Kimura and Crow, 1964). This quantity should basically determine the mean genome-wide heterozygosity expected at neutral sites for any given individual in that population. However, since the neutral mutation-drift balance can be slow to achieve, contemporary genetic diversity often keeps the signature of past demographic fluctuations rather than being entirely determined by the current population size. Therefore, genetic diversity should be well predicted by estimates of *N*_*e*_ that integrate the long-term effect of drift over the coalescent time. Unfortunately, such estimates are very difficult to produce using demographic data only.

Demographic variations set aside, the most proximate determinant of *N*_*e*_ is the actual number of individuals (*N*), also called the census population size. Comparative genomic studies in mammals and birds have showed that current species abundance correlates with the long-term coalescence *N*_*e*_, despite a potential deviation from long-term population stability in several of the species studied (Díez-Del-Molino et al., 2018; Leroy et al., 2020; Peart et al., 2020). General laws in ecology, such as the negative relationship between species abundance and body size (White et al., 2007) have also been used to predict the long-term *N*_*e*_. Higher genetic diversity in small body size species was found in butterflies and Darwin’s finches (Mackintosh et al., 2019; Brüniche-Olsen et al., 2019), while in the latter genetic diversity also positively correlated with island size, another potential proxy for the long-term *N*_*e*_ (Brüniche-Olsen et al., 2019). Surprisingly, however, genetic diversity variation across Metazoans is much better explained by fecundity and propagule size than classical predictors of species abundance such as body size and geographic range (Romiguier et al., 2014). This result has been attributed to differences in extinction risk for species that have contrasted reproductive strategies. Under this hypothesis, species with low fecundity and large propagule size (*K*-strategists) would be more resilient to low population size episodes compared to species with high fecundity and small propagule size (*r*-stategists) which would go extinct if they reach such population sizes (Romiguier et al., 2014). By contrast, Mackintosh et al. (2019) found no effect of propagule size on genetic diversity within Papilionoidea, a family showing little variation in reproductive strategy. Therefore, the major effect of the *r/K* gradient on genetic diversity variation across Metazoa probably hides other determinants that act within smaller branches of the tree of life. In particular, how demography and evolutionary processes influence genetic variation in different taxa remains unclear.

Other factors than fluctuations in population size are known to reduce the value of *N*_*e*_ relative to the census population size, impacting the 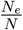 ratio to a different extent from one species to another. These factors include unbalanced sex-ratios, variance in lifetime reproductive success among individuals, age structure, kinship-correlated survival and some metapopulation configuration (Wright, 1969; Falconer, 1989; Lande and Barrowclough, 1987). A potentially strong effect comes from variance in the number of offspring per parent (*V*_*k*_), which reduces *N*_*e*_ compared to *N* following 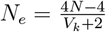 (Crow and Kimura, 1970). Variance in reproductive success can naturally emerge from particular age-specific demographic characteristics summarized in life tables that contain age- (or stage-) specific survival and fecundity rates (Ricklefs and Miller, 1999). The impact of life tables characteristics on expected 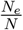 ratio has been the focus of a large body of theoretical and empirical works (Nunney, 1991, 1996; Waples, 2002, 2016b,a; Waples et al., 2018). Accounting for iteroparity and overlapping generations, a meta-analysis of vital rates in 63 species of plants and animals revealed that half of the variance in 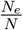 among species can be explained by just two life history traits: adult lifespan and age at maturity (Waples et al., 2013). Interestingly, longevity was the second most important factor explaining differences in genetic diversity across Metazoans (Romiguier et al., 2014). However, there is still no attempt to evaluate the extent to which lifetime variance in reproductive success explains differences in genetic diversity between species with different life table components.

Marine fishes are good candidates to address this issue. They are expected to show a particularly high variance in reproductive success as a result of high abundance, type III survivorship curves (i.e. high juvenile mortality and low adult mortality) and increasing fecundity with age. Consequently, it has been suggested that marine fish species show a marked discrepancy between adult census size and effective population size, resulting in 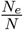 ratios potentially smaller than 10^−3^. The disproportionate contribution of a few lucky winners to the offspring of the next generation is sometimes referred as the “big old fat fecund female fish” (BOFFFF) effect, a variant of the “sweepstakes reproductive success” hypothesis (Hedgecock, 1994; Hedrick, 2005; Hedgecock and Pudovkin, 2011) that is often put forward to explain low empirical estimates of effective population sizes from genetic data (Hauser and Carvalho, 2008). However, subsequent theoretical work showed that low values of 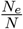 below 0.01 can only be generated with extreme age-structure characteristics (Waples, 2016b). The real impact of lifetime variance in reproductive success on genetic diversity thus remains unclear, even in species like fish in which its impact is supposed to be strong. Contrasting results have been obtained by comparative studies in marine fishes, including negative relationship between diversity and body size (Pinsky and Palumbi, 2014; Waples, 1991), fecundity (Martinez et al., 2018) and overfishing (Pinsky and Palumbi, 2014). However, these studies relied on few nuclear markers, that could provide inaccurate or biased estimates of genetic diversity (Väli et al., 2008). They also compared species sampled from different locations, thus, likely having different demographic histories, which could blur the relationship between species characteristics and genetic diversity (Ellegren and Galtier, 2016).

Here, we compared the genome-average heterozygosity to the life history traits and life table characteristics of 16 marine teleostean species sharing similar Atlantic and Mediterranean distributions. We estimated genetic diversity from unassembled whole-genome reads using `GenomeScope` (Vurture et al., 2017) and checked the validity of these estimates with those obtained using a high-standard reference-based variant calling approach. Using this data, we related species genetic diversity to eight simple quantitative and qualitative life history traits. Then, we built species life tables and determined if the lifetime variance in reproductive success induced by these tables could explain observed differences in genetic diversity using an analytical and a forward-in-time simulation approach. Finally, we generalized our findings by exploring the influence of age-specific survival and fecundity rates on the variance in reproductive success and ultimately genetic diversity via simulated lifetimes tables.

## Material and Methods

### Sampling, DNA extraction and whole-genome sequencing

We sampled 16 marine teleostean fish species presenting a wide diversity of life history strategies expected to affect genetic diversity (Table 1). All these species share broadly overlapping distributions across the North-eastern Atlantic and Mediterranean regions. Sampling was performed at the same four locations for all species: two in the Atlantic (the Bay of Biscay in South-western France or North-western Spain and the Algarve in Portugal), and two in the Western Mediterranean Sea (the Costa Calida region around Mar Menor in Spain and the Gulf of Lion in France see Fig 1A). Individual whole-genome sequencing libraries were prepared following the Illumina TruSeq DNA PCR-Free Protocol and sequenced to an average depth of 20X on an Illumina NovaSeq 6000 platform by Genewiz Inc (USA). Raw reads were preprocessed with `fastp` v.0.20.0 (Chen et al., 2018) using default parameters (see Supplementary Material).

**Table 1-.** Life history traits and observed genetic diversity of the 16 teleostean marine species. -For each species, number of individuals used for the estimation of genetic diversity; observed median genetic diversity among all individuals (± standard deviation); body size (in centimeters); trophic level; age at first maturity (in years), lifespan (in years), adult lifespan (in years, defined as the difference between lifespan and age at maturity), parental care behaviour (– = no egg protection; NG = nest-guarders; MP = male brood-pouch) and hermaphroditism (– = no hermaphroditism; PG = protogynous; PA = protandrous, RUD = rudimentary). Detailed bibliographic references are provided in supplementary material.

| Species | Vernacular name | N. | Genetic diversity (%) | Body size (cm) | Trophic level | Fecundity | Propagule Size (mm) | Maturity (years) | Lifespan (years) | Adult lifespan (years) | Parental care | Herma. |
| --- | --- | --- | --- | --- | --- | --- | --- | --- | --- | --- | --- | --- |
| *Coryphoblennius galerita* | Montagu's blenny | 16 | 0.607(±0.014) | 7 | 2.28 | NA | 3.3 | 1.5 | 6 | 4.5 | NG | – |
| *Coris julis* | Rainbow wrasse | 20 | 1.172(±0.056) | 27.2 | 3.24 | 169.81 | 0.63 | 1 | 7 | 6 | – | PG |
| *Dicentrarchus labrax* | European sea bass | 20 | 0.375(±0.031) | 102.15 | 3.47 | 12436.52 | 1.15 | 3 | 15 | 12 | – | – |
| *Diplodus puntazzo* | Sharp-snout seabream | 19 | 0.533(±0.074) | 49.69 | 3.07 | 277.87 | 0.87 | 2 | 10 | 8 | – | RUD |
| *Hippocampus guttulatus* | Long-snouted seahorse | 12 | 0.313(±0.090) | 19.8 | 3.5 | 1.21 | 12 | 0.5 | 5 | 4.5 | MP | – |
| *Lophius budegassa* | Blackbellied angler | 20 | 0.225(±0.015) | 103 | 4.23 | 2304.03 | 1.88 | 7.5 | 21 | 13.5 | – | – |
| *Lithognathus mormyrus* | Striped seabream | 20 | 0.553(±0.027) | 37.85 | 3.42 | 214.09 | 0.75 | 2 | 12 | 10 | – | PA |
| *Merluccius merluccius* | European hake | 20 | 0.844(±0.025) | 88.9 | 4.43 | 2294.54 | 1.07 | 3 | 11 | 8 | – | – |
| *Mullus surmuletus* | Striped red mullet | 19 | 1.135(±0.048) | 30.18 | 3.46 | 2569.32 | 0.86 | 1.5 | 6 | 4.5 | – | – |
| *Pagellus erythrinus* | Common pandora | 19 | 1.100(±0.020) | 36 | 3.46 | 2280.46 | 0.77 | 2 | 8 | 6 | – | PG |
| *Serranus cabrilla* | Comber | 19 | 1.205(±0.055) | 30.8 | 3.68 | 37.97 | 0.91 | 2 | 6 | 4 | – | – |
| *Spondyliosoma cantharus* | Black seabream | 19 | 0.478(±0.034) | 35.7 | 3.27 | 425.62 | 2.1 | 3 | 10 | 7 | NG | PG |
| *Symphodus cinereus* | Grey wrasse | 10 | 0.660(±0.125) | 14.1 | 3.3 | 13.20 | 2.87 | 1.5 | 6 | 4.5 | NG | – |
| *Sardina pilchardus* | European pilchard | 20 | 1.415(±0.182) | 20.35 | 2.94 | 22.89 | 1.64 | 1 | 5 | 4 | – | – |
| *Syngnathus typhle* | Broadnosed pipefish | 20 | 0.859(±0.047) | 26.2 | 3.75 | 0.38 | 20 | 1 | 3 | 2 | MP | – |
| *Sarda sarda* | Atlantic bonito | 20 | 0.896(±0.208) | 68.9 | 4.34 | 15647.73 | 1.3 | 1 | 4 | 3 | – | – |

**Figure 1:**
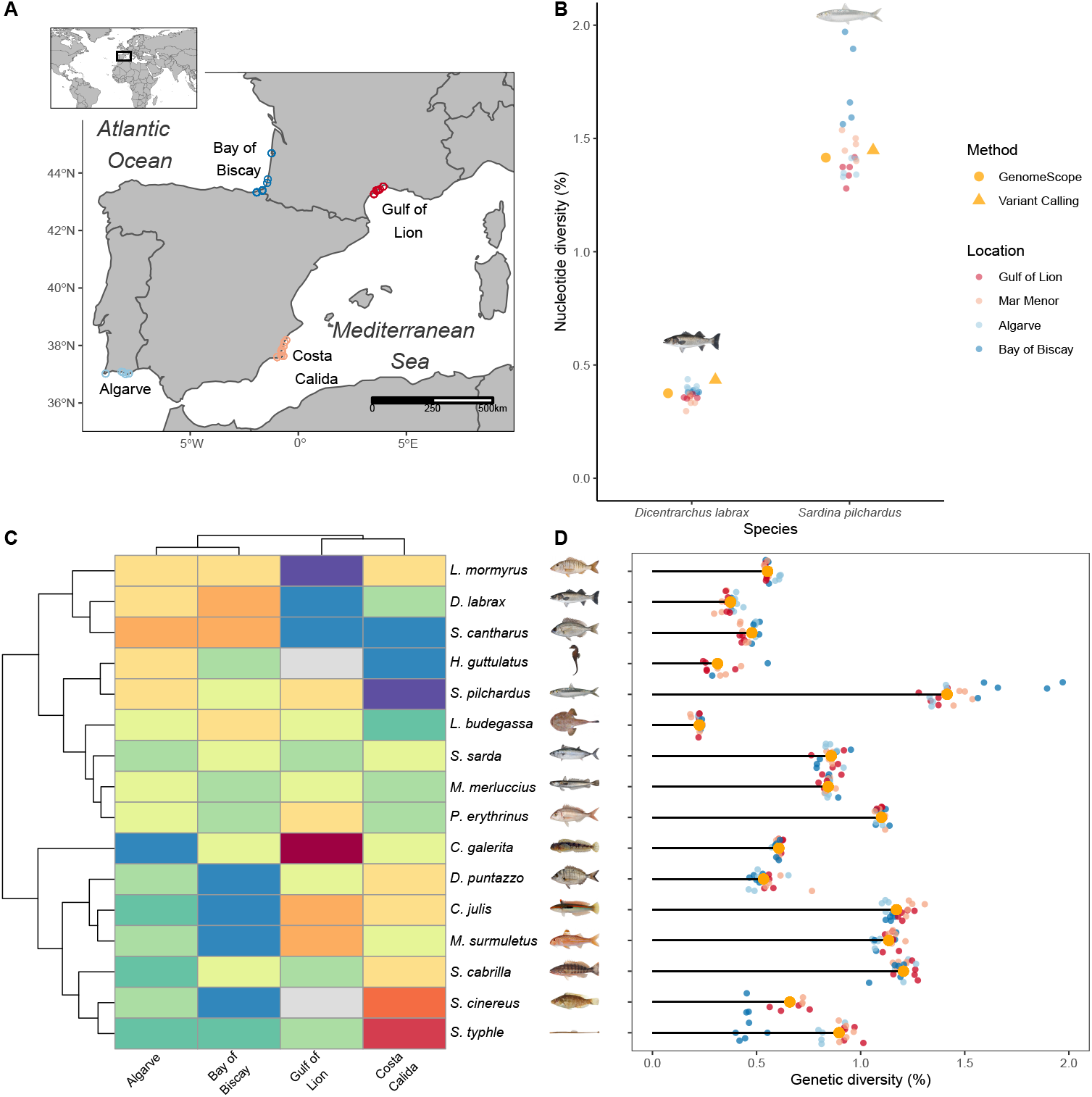
Sampling and estimation of genetic diversity in 16 marine fish species-. In panels A, B and D, the geographical origin of samples is represented by colors. Atlantic: Bay of Biscay (dark blue), Faro region in Algarve (light blue). Mediterranean: Murcia region in Costa Calida (pink), Gulf of Lion (red). **(A)** Sampling map of all individuals included in this study. Each point represents the coordinates of a sample taken from one of four locations: two in the Atlantic Ocean and two in the Mediterranean Sea. **(B)** Genome-wide diversity in the European pilchard (*S. pilchardus*) and European sea bass (*D*.*labrax*) estimated after variant calling (orange triangle) or from GenomeScope (orange dot: median; smaller dots: individual estimates) **(C)** Heatmap clustering showing the variance in genetic diversity within species among locations. Each line represents one species, with the corresponding species name written on the right side; every column represents one location. Blue and red colors respectively indicate higher and lower genetic diversity within a location for a given species compared to the average species genetic diversity. **(D)** Individual and median genetic diversity within each species estimated with GenomeScope. Species illustrations were retrieved from Iglésias (2013) with permissions.

### Estimation of genetic diversity

We used `GenomeScope` v.1.0 to estimate individual genome-wide heterozygosity (Vurture et al., 2017). Briefly, this method uses a *k* -mers based statistical approach to infer overall genome characteristics, including total haploid genome size, percentage of repeat content and genetic diversity from unassembled short-read sequencing data. We used `jellyfish` v.2.2.10 to compute the *k* -mer profile of each individual (Marçais and Kingsford, 2011). The genetic diversity of each species was determined as the median of the individual genome-wide heterozygosity values. We chose the median instead of the mean diversity since it is less sensitive to the possible presence of individuals with non-representative genetic diversity values (e.g. inbred or hybrid individuals) in our samples.

In order to assess the reliability of `GenomeScope` and detect potential systematic bias, we compared our results with high-standard estimates of genetic diversity obtained after read alignment against available reference genomes (see details in Supplementary Material). To perform this test, we used the sea bass (*D. labrax*) and the European pilchard (*S. pilchardus*), two species that represent the lower and upper limits of the range of genetic diversity in our dataset (Table 1, Fig 1D).

### Life history traits database

We collected seven simple quantitative variables describing various aspects of the biology and ecology of the 16 species: body size, trophic level, fecundity, propagule size, age at maturity, lifespan and adult lifespan (Table 1, Table S4 for detailed informations on bibliographic references). We used the most representative values for each species and each trait when reported traits varied among studies due to plasticity, selection or methodology. In addition, we collected two qualitative variables describing the presence/absence of hermaphroditism and brooding behaviour, as revealed by males carrying the eggs in a brood pouch (*H. guttulatus* and *S. typhle*) or nest-guarding (*C. galerita, S. cinereus* and *S. cantharus*). Detailed information on data collection is available in Supplementary Material.

### Construction of life tables

Life tables summarize survival rates and fecundities at each age during lifetime (Ricklefs and Miller, 1999). Thus, they provide detailed information on vital rates that influence the variance in lifetime reproductive success among individuals. This tool is well designed to describe population structure from the probability of survival to a specific age at which a specific number of offspring are produced. Ideally, age-specific survival is estimated by direct demographic measures, such as mark-recapture. Unfortunately, direct estimates of survival were not available for the 16 studied species. We thus followed Benvenuto et al. (2017) to construct species life tables. Age-specific mortality of species *sp, m*_*sp,a*_, is a function of species body length at age *a, L*_*sp,a*_, species asymptotic Von Bertalanffy length *L*_*sp,inf*_, and species Von Bertalanffy growth coefficient, *K*_*sp*_ :

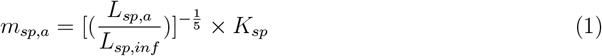

Age-specific survival rates, *s*_*sp,a*_ were then estimated as:

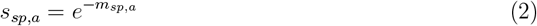

We collected age-specific length from empirical data and estimated *L*_*inf*_ and *K* values from age-length data as explained in the appendix, setting survival probability to zero at the maximum age (Appendix 1). When differences in age-specific lengths between sexes were apparent in the literature, we estimated a different age-specific survival curve for each sex. The relationship between absolute fecundity and individual length is usually well fitted with the power-law function (*F* = *αL*^*β*^), although some studies also used an exponential function (*F* = *αe*^*βL*^) or a linear function (*F* = *α* + *Lβ*). We collected empirical estimates of *α* and *β* and determined age-specific fecundity from the age-specific length and the fecundity-length function reported in the literature for each species. Fecundity was set to zero before the age at first maturity.

### Effect of the variance in reproductive success on the Ne/N ratio

To understand how differences in life tables drive differences in genetic diversity between species, we estimated the variance in lifetime reproductive success, *V*_*k*_ and the ensuing ratio 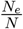 using the analytic framework developed in `AgeNe` (Waples et al., 2011). `AgeNe` infers *V*_*k*_ using informations from life tables only. Hence, the estimated variance in reproductive success estimated is only generated by inter-individual differences in fecundity and survival. `AgeNe` assumes constant population size, stable age structure, and no heritability of survival and fecundity. We used the life tables constructed as described above and set the number of new offspring to 1000 per year. This setting is an arbitrary value which has no influence on the estimation of either *V*_*k*_ nor 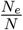 by `AgeNe`. For all species, we set an initial sex ratio of 0.5 and equal contribution of individuals of the same age (i.e. no sweepstake reproductive success among same-age individuals). We ran `AgeNe` and estimated 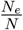 for each species.

Four life tables components can generate differences in 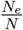 between species: age at maturity, age-specific survival rates, age-fecundity relationships and sex-related differences in these components. To determine the role that each parameter plays in shaping levels of genetic diversity among species, we built 16 alternative life tables where the effect of each component was added one after the other, while the others were kept constant across species. Thus, in our null model, age at maturity was set at 1 year old for all species, fecundity and survival did not vary with age (constant survival chosen to have 0.01% of individuals remaining at maximum age, following Waples (2016b)), and there were no differences between sexes. Next, the effect of each component was tested by replacing these constant values with their biological values in species’ life tables. For each of the 16 life tables thus constructed, we tested whether variation in 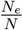 explained the variation in observed genetic diversity after scaling these two variables by their maximum value. With this scaling, the correlation between 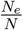 and genetic diversity should overlap with the *y* = *x* function in cases where a decrease in 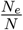 predicts an equal decrease in genetic diversity, indicating a strong predictive power of the components induced in life tables.

### Forward simulations

A complementary analysis of the contribution of life table properties on genetic diversity was performed using forward simulations in `SLiM` v.3.3.1 (Haller and Messer, 2017). Stochastic forward simulations allow a different formalization compared to the deterministic model implemented in `AgeNe`. Thus, they provide another approach to the problem and can lead to a more intuitive understanding of why vital rates affect *N*_*e*_ over the long-term, and ultimately genetic diversity. We simulated populations with overlapping generations, sex-specific lifespan, and age- and sex-specific fecundity and survival. We used life tables estimated as previously, and sex-specific lifespan estimates were collected in the literature as described above. Age and species-specific fecundity were determined as previously and scaled between 0 (age 0) and 100 (maximum age) within each species. In the simulations, each individual first reproduces and then either survives to the next year or dies following a probability determined by its age and the corresponding life table. We kept population size constant and estimated the mean genetic diversity (i.e., the proportion of heterozygous sites along the locus) over the last 10000 years of the simulation after the mutation-drift equilibrium was reached and using 50 replicates (see Supplementary Material for further informations).

As previously, we evaluated the contribution of each component among 8 alternative life tables by comparing scaled observed and simulated genetic diversity.

### Evaluating the impact of life tables beyond marine fish

To generalize our understanding of the influence of life tables on genetic diversity beyond the species analyzed in this study, we simulated a wide range of age-specific survival and fecundity curves and explored their effect on the relationship between adult lifespan and variance in reproductive success. To this end, we defined 16 theoretical species with age at first maturity and lifespan equal to that of our real species and then introduced variation in survival and fecundity curves. First, age-specific mortality was simulated following Pinder et al. (1978):

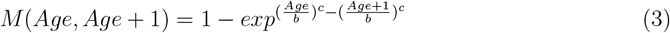

where *c* defines the form of the survivorship curve, with *c* > 1, *c* = 1 and *c* < 1 defining respectively a *Type I* (e.g. mammals), *Type II* (e.g. birds) and *Type III* (e.g. fish) survival curves. We took values of *c* from 0.01 to 30 (Fig 4A). Parameter *b* was equal to 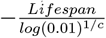 to scale survivorship curves in such a way that 1% of the initial population remains at maximum age.

Second, age-specific fecundity was simulated with two models: constant and exponential. In the first model, fecundity is constant for all ages since maturity. In the second model, fecundity increases or decreases exponentially with age following *F*_*Age*_ = *exp*^*f*×*Age*^, as it is often observed in marine fishes (Curtis and Vincent, 2006). We first set *f* = 0.142 as the median of the *f* values for the 16 species. Secondly, we took values of *f* ranging from −1 to 1 (Fig 4A). We scaled maximum fecundity to 1 for all simulations.

For each combination of *c* and *f*, and for each fecundity model, we simulated all species life tables given age at maturity and lifespan. Then, we ran `AgeNe` and estimated 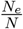 for each simulated species and estimated the slope of the regression between adult lifespan and 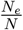 across all 16 species. We explored the impact of alternative fecundity-age models on the relationship between adult lifespan and 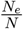 (see details in Supplementary Material).

### Intraspecific variation in genetic diversity

We addressed the potential effects of population structure, demography and historical contingencies on genetic diversity by examining the extent of spatial variation in genetic diversity between the four populations within each species. First, we evaluated the relative amount of intraspecific compared to interspecific variation in genetic diversity. Then, we applied a *z*-transformation of individual genetic diversity within each species to put spatial differences in within-species diversity on the same scale. In order to detect similar spatial patterns of genetic diversity among species, we finally performed a hierarchical clustering analysis of the matrix of *z*-transformed genetic diversity values with `pheatmap` function available in `pheatmap v1.0.12` R package.

### Statistical analyses

All statistical analyses were carried out using `R-3.6.1` (R Core Team, 2018). We fitted beta regression models between genetic diversity and any covariate with the R-package `betareg` v.3.1-3 (Cribari-Neto and Zeileis, 2010). We tested statistical interactions between any quantitative and qualitative covariates using likelihood tests with the `lmtest` v.0.9-37 package (Zeileis and Hothorn, 2002).

## Results

### Whole-genome resequencing data set

We resequenced 300 individual genomes from 16 marine teleostean species, with high read quality scores (mean Q30 rate = 92.4%) and moderate duplication rates (10.8%) (Fig S2). GC content was moderately variable among species and highly consistent among individuals of the same species, except for three individuals that showed a marked discrepancy with the overall GC content of their species (Fig S2). These three individuals were thus removed from downstream analyses to avoid potential issues due to contamination or poor sequencing quality.

### Estimation of genetic diversity with GenomeScope

The `GenomeScope` model successfully converged for all of the 297 individual genomes retained (Fig S6E). The average depth of sequencing coverage per diploid genome exceeded 20X in most individuals. Estimated genome sizes were very consistent within species (Fig S6A-C). Estimated levels of genetic diversity were also homogeneous among individuals of the same species with some few exceptions (e.g. *S. cinereus* and *S. typhle*) and most of the variability in genetic diversity was observed between species (Fig 1D). Two individuals (one *D. puntazzo* and one *P. erythrinus*) showed a surprisingly high genetic diversity (more than twice the average level of their species), indicating possible issues in the estimation of genome-wide heterozygosity. Therefore we removed these individuals from subsequent analysis, although their estimated genome size and GC content matched their average species values (therefore excluding contamination as a cause of genetic diversity estimation failures).

Observed values of genetic diversity ranged from 0.225% for *L. budegassa* to 1.415% for *S. pilchardus*. We found no correlation between species genetic diversity and genome size (*p* − *value* = 0.983). The estimation of genetic diversity was robust to the choice for *k*-mer lengths ranging from 21 to 25, suggesting a low sensiblity of `GenomeScope` regarding this parameter (Fig S4). The fraction of reads mapped against reference genomes ranged between 96.72 and 98.50% for *D. labrax* and between 87.45 and 96.42 % for *S. pilchardus* (Table S2; Fig S3). We found similar species genetic diversity estimates between `GenomeScope` and the `GATK` reference-based variant calling approach for the two control species, representing the two limits of the range of genetic diversity in our dataset (Fig 1B).

### Adult lifespan is the best predictor of genetic diversity

We evaluated the effect of several key life history traits that potentially affect species genetic diversity (Table S1).

Two widely used predictors of population size, body size and trophic level, were not significantly correlated to genetic diversity (*p-value* = 0.119 and 0.676 respectively, Fig S8A-B). Although we detected a significant negative relationship between the logarithm of fecundity and propagule size (*p-value* = 0.00131, slope = − 0.4385 ± 0.1076) as in Romiguier et al. (2014), we found no significant correlation between either propagule size (*p-value* = 0.561), or the logarithm of fecundity (*p-value* = 0.785) and genetic diversity (Fig S8C-D).

By contrast, both lifespan (*p-value* = 0.011) and adult lifespan (*p-value* = 0.007) were significatively negatively correlated with genetic diversity (Table S1, Fig 2). The percentage of variance explained by each variable reached 43.8 and 42.9 %, respectively. Repeating the same statistical analyses with genetic diversity estimates either only from mediterranean or atlantic individuals led to the same results, revealing no effect of within-species population structure on the relationship between genetic diversity and life history traits (Fig 1C, Fig S9, Table S3).

**Figure 2:**
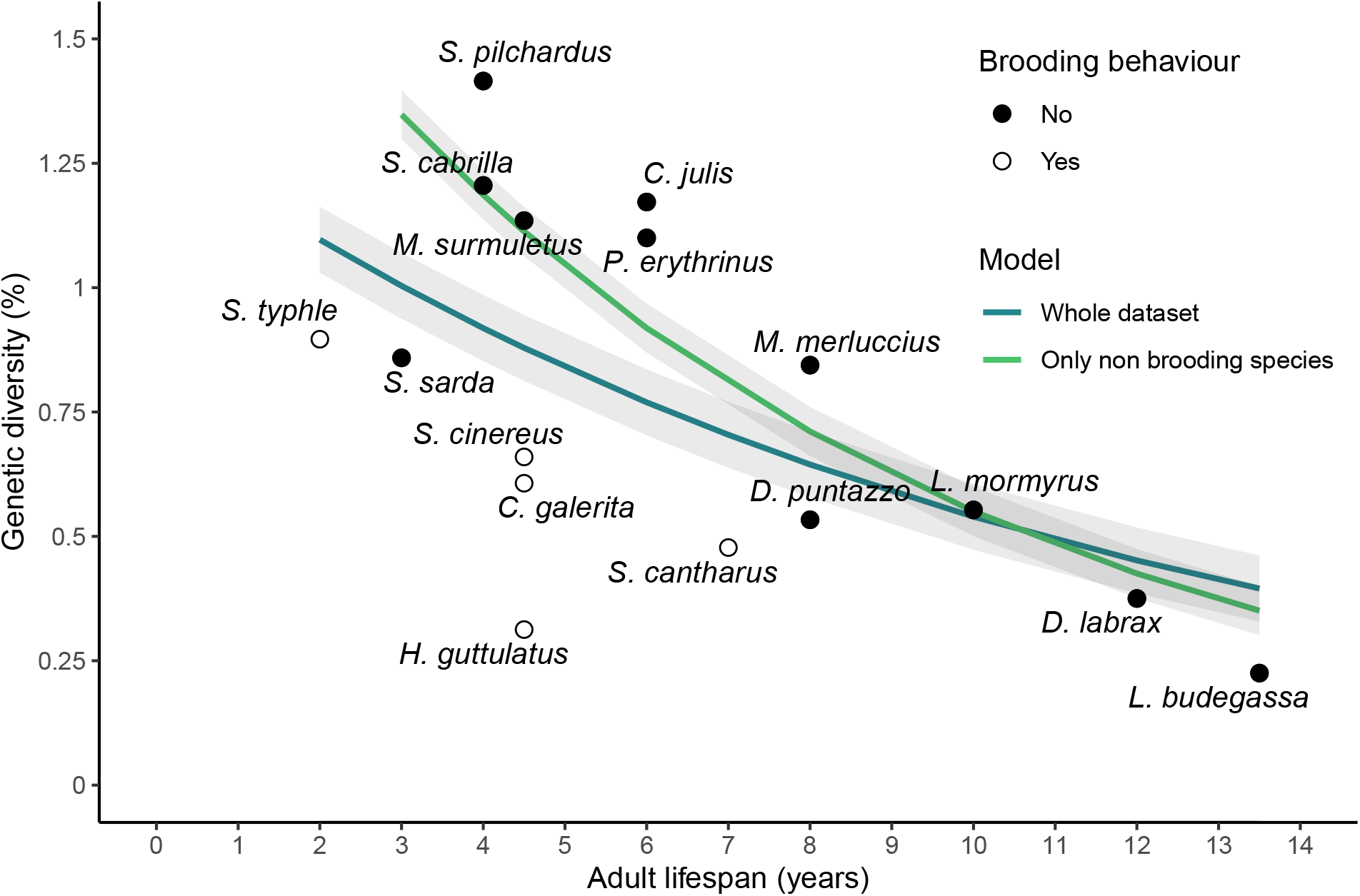
Relationship between species median genetic diversity (%) and adult lifespan-. Each point represents the median of the individual genetic diversities for a given species. Adult lifespan is defined as the difference between lifespan and age at first maturity in years. Dot points and empty circles represent non-brooding species and brooding species, respectively. Blue and green lines represent the beta regression between adult lifespan and genetic diversity considering either the whole dataset (16 species), or the 11 non-brooding species only, respectively.

We found no significant interaction between hermaphroditism and any of the previous variables on genetic diversity. By contrast, parental care showed a significant interaction with lifespan (*p-value* = 0.0011), adult lifespan (*p-value* = 0.0008) and body size (*p-value* = 0.0035) on genetic diversity. Brooding species (nest protection by males for *C. galerita, S. cinereus* and *S. cantharus* and male abdominal brood-pouch for *H. guttulatus* and *S. typhle*) had systematically lower genetic diversity than non-brooding species with similar lifespan.

When considering only non-brooding species, we found steeper negative correlations and higher percentages of between-species variance in genetic diversity explained by lifespan (*p-value* = 1.017*e*^−7^, pseudo-*R*^2^ = 0.851) and adult lifespan (*p-value* = 1.645*e*^−7^, pseudo-*R*^2^ = 0.829, Fig 2, Table S1). To test the relevance of considering this sub-dataset, we estimated the slope of the regression and the pseudo-*R*^2^ for all combinations of 11 out of 16 species and compared the distribution of these values to the estimated slope and pseudo-*R*^2^ obtained for the 11 non-brooding species (Fig S13). The estimated slope for non-brooders lied outside of the 95% confidence interval of the distribution of estimated slopes (*slope* = − 0.129, 95% CI = [− 0.122, − 0.049]) and the same was found for pseudo-*R*^2^ (pseudo-*R*^2^ = 0.829, 95% CI = [0.073, 0.727]). Furthermore, considering non-brooding species only, there was still no significant correlation between genetic diversity and trophic level (*p-value* = 0.259), propagule size (*p-value* = 0.170), and fecundity (*p-value* = 0.390), but genetic diversity appeared significantly negatively correlated to body size (*p-value* = 6.602*e*^−5^, pseudo-*R*^2^ = 0.616). We did not detect any significant correlation between any trait variable and genetic diversity within the sub-dataset of brooding species. However, this should be taken with caution given the very low number of brooding species (*n* = 5) in our dataset.

Body size and lifespan were highly positively correlated traits in our dataset (*p-value* = 0.0013, *R*^2^ = 0.536, Fig S7). Thus, using empirical observations only, it was not possible to fully disentangle the impact of each of these traits among the possible determinants of genetic diversity in marine fishes. However, we found important differences in effect sizes for body size (*slope* = − 0.014), lifespan (−0.095) and adult lifespan (− 0.129), which rule out body size as a major determinant of diversity in our dataset.

### Variance in reproductive success explains levels of observed genetic diversity

To understand the mechanisms by which adult lifespan affects genetic diversity and test if it can alone explain our results, we built life tables for each of the 16 species by gradually incorporating age-specific fecundity and survival, age at first maturity, lifespan and sex-specific differences in these parameters.

Non-genetic estimates of 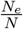 ratio obtained with `AgeNe` ranged from 0.104 in *L. budegassa* to 0.671 for *S. cinereus*. When considering the 16 species together, the 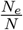 ratio was not significatively correlated with genetic diversity (*p* − *value* = 0.0935). However, four out of five brooding species had low genetic diversity despite high 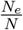 ratios (Fig 3A). As previously observed, removing the 5 brooders increased the slope and the percentage of variance of genetic diversity explained by the 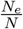 ratio above null expectations obtained by removing groups of 5 species at random (*slope* = 1.849, 95% CI = [0.048, 1.582], pseudo-*R*^2^ = 0.55, 95% CI = [0.004, 0.533], Fig S14). Thus, the 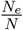 ratio predicted by life tables was positively correlated to genetic diversity when considering non-brooding species only (Fig 3A).

**Figure 3:**
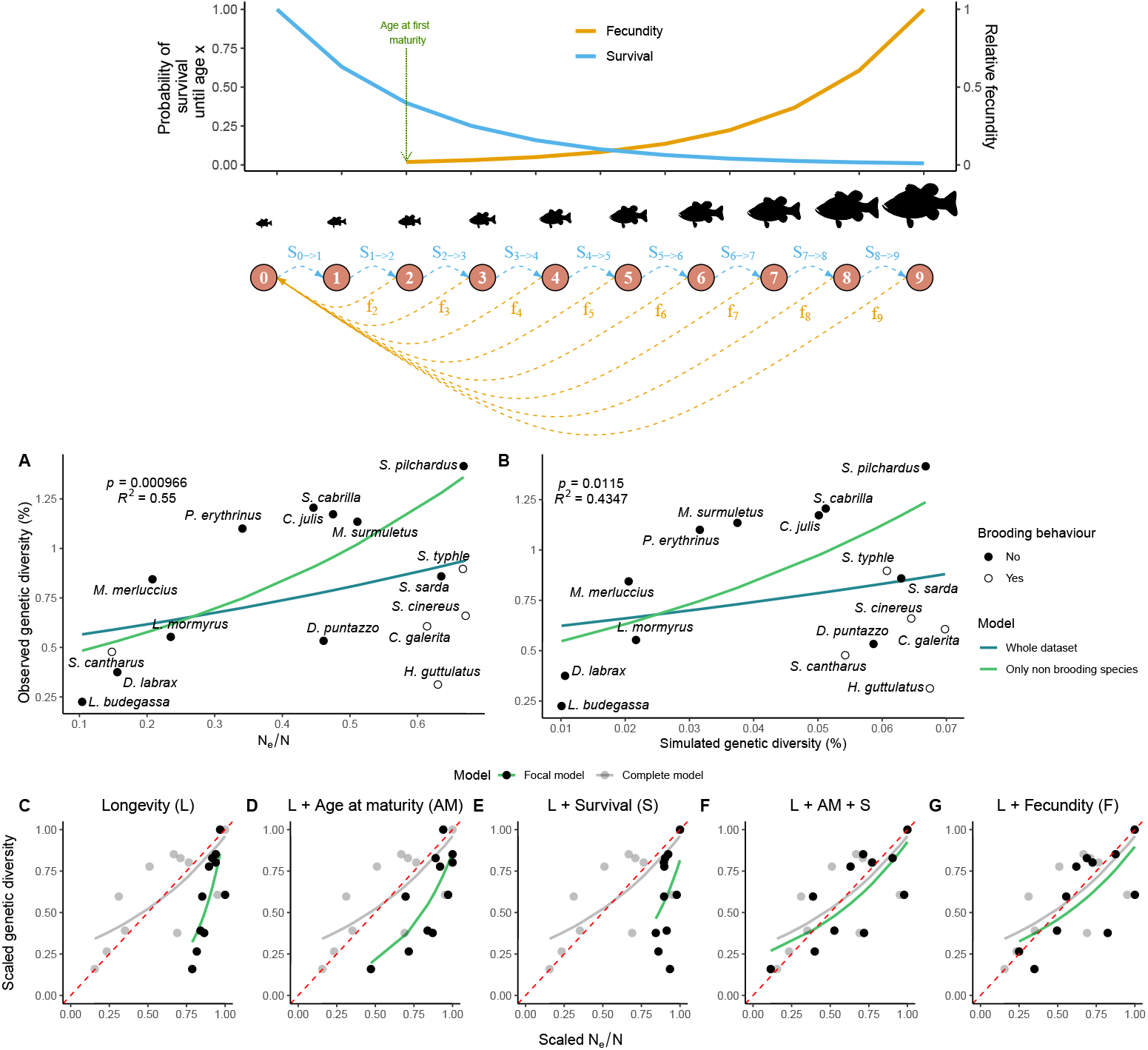
Variance in reproductive success induced by age-specific vital rates and adult lifespan correlate with observed genetic diversity-. On top, schematic illustration of age-specific fecundity (*f*_*age*_, in orange) and survival (*S*_*age*−>*age*+1_, blue) for a simulated species. (A) and (B) represents the relationship between observed genetic diversity on the *y*-axis and, respectively, 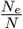 estimated by `AgeNe`, and simulated genetic diversity with forward-in-time simulations in `SLiM` v.3.31 (Haller and Messer, 2017), on *x*-axis. Life tables containing information on age-specific survival, fecundity and lifespan were used for the 16 species. Age at maturity was used only with `AgeNe`. Dot points represent non-brooding species and empty circles, brooding species. Blue and green lines represent the beta regression between adult lifespan and genetic diversity considering the whole dataset (16 species), and the 11 non-brooding species only, respectively. The *p* −*value* and the pseudo-*R*^2^ are represented on the top left for each of the two top panels for the non-brooders model. Panels (C)-(G) represent the relationship between scaled genetic diversity and scaled 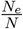 (i.e., divided by the maximum corresponding value) for the 11 non-brooding species. In each panel, the grey points represent scaled 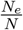 estimated from life tables with age at maturity, age-specific fecundity and survival and sex-specific differences (as in panel A). Black points are scaled estimates of 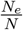 from life tables with only: C) longevity (L), D) longevity (L) and age at maturity (AM), E) longevity (L) and age-specific survival (S), F) longevity (L), age at maturity (AM) and age-specific survival (S) and G) longevity (L) and age-specific fecundity (F). Beta regression models (grey and green lines) that closely overlap the red dotted line indicate that a decrease in 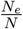 leads to a similar decrease in genetic diversity.

**Figure 4:**
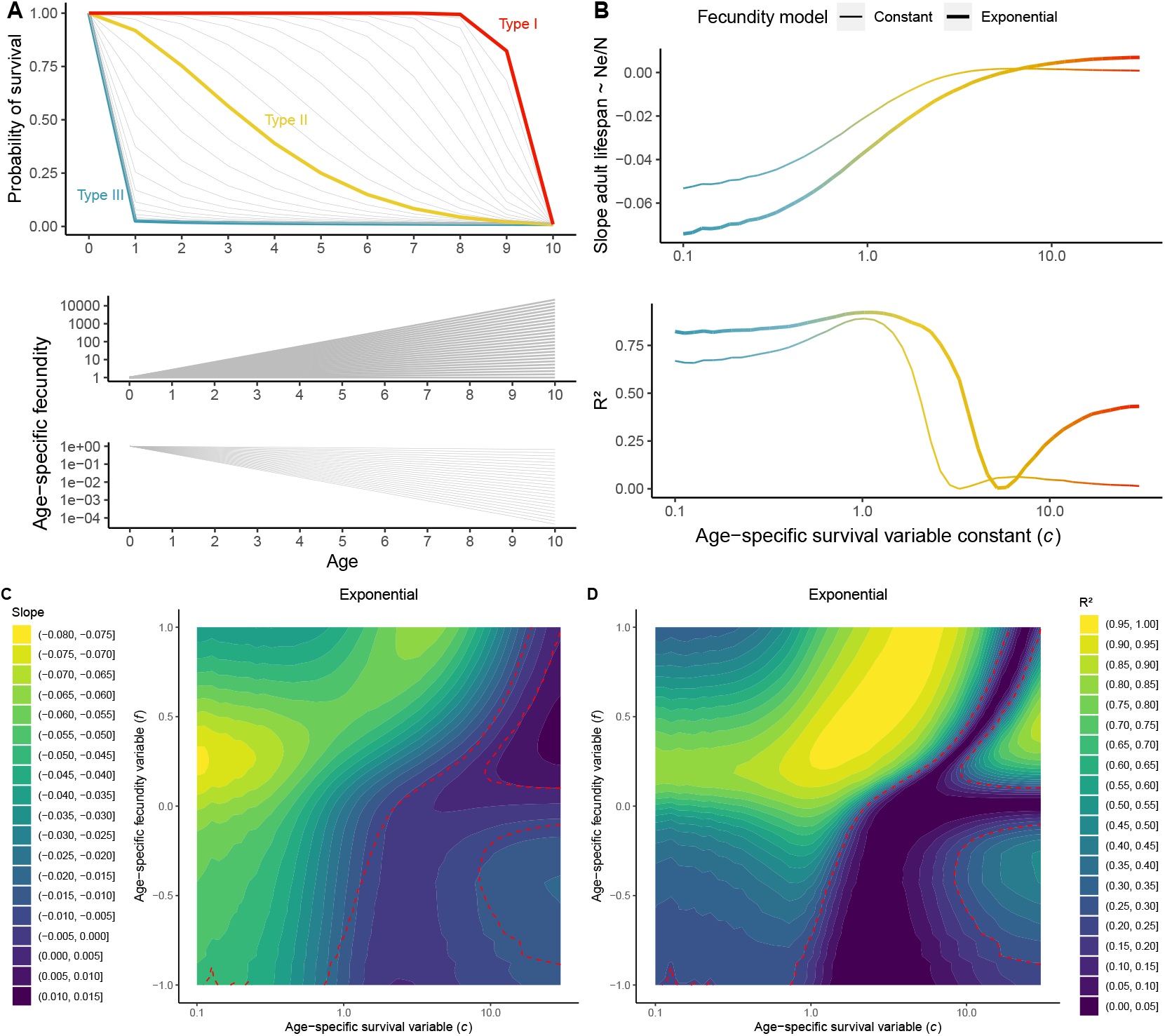
Slope of the linear model between adult lifespan and 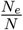 ratio estimated with AgeNe for different combinations of age-specific survival and fecundity-. A) On top, gradient of survivorship curves simulated, ranging from type III (blue, *c* < 1), high juvenile mortality and low adult mortality; to type II (orange, *c* around 1), constant mortality and type I (red), low juvenile mortality high adult mortality. At the bottom, simulated fecundity either increases or decreases exponentially with age as *F*_*Age*_ = *exp*^*f*×*Age*^, with *f* ranging from -1 to 1. 16 simulated life tables were constructed with the same values of age at maturity and lifespan as the 16 studied species, and all possible survivorship curve and fecundity-age relationship shown in Panel A. B) Slope and *R*^2^ of the regression between adult lifespan and 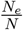 ratio for the 16 simulated species as a function of *c*, for constant fecundity with age (thin line) and exponential increase of fecundity with age with *f* = 0.142 (thick line). C) Slope and D) *R*^2^ of the regression between adult lifespan and 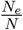 ratio for the 16 simulated species for a gradient of values of *c* and *f*. In C), warmer colors indicate steeper slopes; in D) higher *R*^2^.

Our next step was to determine the impact of each component of life tables as well as their combinations on genetic diversity (Fig 3C-G). Starting from a null model (Fig 3C), in which species life tables differed only in lifespan, we found that the 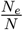 ratio ranged from 0.558 to 0.733, a variance much lower than that of observed genetic diversities. Then, adding separately age at maturity (Fig 3D) or age-specific survival (Fig 3E) did not better predict the range of observed genetic diversities. However, combining age at maturity and age-specific survival (Fig 3F) or adding only age-specific fecundity (Fig 3G) enable us to explain the range of observed diversity values. Finally, combining these three parameters together (age at maturity, age-specific survival, and fecundity, model 8, Fig S10H) resulted in the best fit for both the slope and the intercept and for both non-brooding species and the whole data set. Adding sex-specific differences in life tables did not improve the fit, however (models 9 to 16, Fig S10I-P).

Our final step was to further explore the role of the variance in reproductive success on genetic diversity by simulating genetic diversity at mutation-drift equilibrium with the age-specific vital rates of the 16 species.

We simulated a population of 2000 individuals with age-specific survival and fecundity. As expected, including age-specific vital rates decreased the equilibrium level of genetic diversity compared to expectations under the classical Wright-Fisher model (*θ* = 4*N*_*e*_*µ* = 0.08%). It was reduced to 0.070% in the species with the least effect of age-specific vital rates (*C. galerita*), and down to 0.010% in the species with the greatest effect (*L. budegassa*). Again, simulated genetic diversity was not correlated to genetic diversity considering all 16 species (*p-value* = 0.297, Fig 3B), but significantly positively correlated within the sub-sample of the 11 non-brooding species (*p-value* = 0.0115).

### Life tables drive correlation between lifespan and the Ne/N ratio

In order to determine the general effect of life table properties on the relation between adult lifespan and 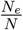 beyond the case of marine fish, we modeled 16 life tables with age at maturity and lifespan similar to those observed in our species but with simulated age-specific survival and fecundity (Fig 4A).

Considering models including constant fecundity with age, we found a significant relationship between adult lifesan and 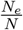 for species with type III survivorship curves (*c* < 1) but not for species having an age-specific survivorhip curve constant, *c*, superior to 2, including type I species (Fig 4B). The slope between adult lifespan and 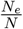 was steepest for type III species, reaching -0.053 for *c* = 0.1. For *c* < 2, the percentage of variation in 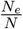 explained by adult lifespan was higher than 60%. Interestingly, it reached a maximum for *c* = 1.03 at 89% and abruptly dropped down around *c* = 2 (Fig 4B).

Then, we added an exponential increase in fecundity with age, first taking *f* = 0.142, which is close to the empirical estimations for our 16 species (Fig 4B). The slope between adult lifespan and 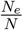 became steeper for type I and type II species and reached -0.074 for extreme type III species (*c* = 0.01). When we included this exponential increase of fecundity with age, the percentage of variation explained was superior for approximately all values of *c*, and the abrupt drop of the percentage of variation explained shifted toward higher *c* values, around *c* = 3. Interestingly, we found significant positive relationships associated with low slope values when *c* became superior to 10 (type I species).

Then, we compared values of slope and *R*^2^ for all *c* values and for *f* ranging from -1 to 1 (Fig 4C-D). The steepest slope between adult lifespan and 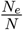 that we obtained reached -0.076 for extreme type III species (*c* around 0.1), and exponential constant, *f*, between 0.18 and 0.31. For type III and type II species (*c* < 1), both the slope and the percentage of variation explained first increased with increasing exponential constant and then decreased. Significant negative relationships were found for *c* < 1 for any values of *f*, except some extreme values near -1, whereas no significant relationship was found for *c* > 1 when *f* is negative except for values of *c* near 1 and values of *f* near 0. The steepest slope and the highest percentage of variation explained were obtained for type III species with intermediate values of *f* (0.1 < *f* < 0.5) and for type II species (1 < *c* < 5) for positive values of *f*. For type I species, as *c* values increased, higher values of *f* are needed to obtain a significant negative relationship between adult lifespan and the 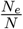 ratio. Above *c* > 20, no significant negative relationship was found for any values of *f*. Again, we found significant positive relationships and low slopes for *c* > 15 and intermediate positive values of *f*.

We found similar results considering a power-law relationship between age and fecundity, with slightly flatter slopes between 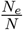 and adult lifespan, and no significant correlations for extreme positive values of *f* and extreme low values of *c*. In contrast, we found limited or no impact of *f* on the relationship between 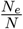 and adult lifespan, respectively, for the linear and the polynomial age-fecundity model.

## Discussion

In this study, we used whole-genome high-coverage sequencing data to estimate the genetic diversity of 16 marine teleost fish with similar geographic distribution ranges. We found that adult lifespan was the best predictor of genetic diversity, species with long reproductive lifespans generally having lower genetic diversities (Fig 2). Longevity was already identified as one of the most important determinants of genetic diversity across Metazoans and plants, in which it also correlates with the efficacy of purifying selection (Romiguier et al., 2014; Chen et al., 2017). A positive correlation between longevity and the ratio of nonsynonymous to synonymous substitutions (*dN/dS*) was also found in teleost fishes (Rolland et al., 2020), thus suggesting lower *N*_*e*_ in long-lived species. However, the mechanisms by which lifespan impacts genetic diversity remain poorly understood and may differ among taxonomic groups. Here we showed that age-specific fecundity and survival (i.e. vital rates), summarized in life tables, naturally predict the empirical correlation between adult lifespan and genetic diversity in marine fishes.

### Impact of life tables on genetic diversity

On a broad taxonomic scale including plants and animals, Waples et al. (2013) showed that almost half of the variance in 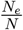 estimated from life tables can be explained with only two life history traits: age at maturity and adult lifespan. Therefore, the effect of adult lifespan on genetic diversity should reflect variations in age-specific fecundity and survival across species. If the species vital rates used to derive 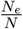 ratios are relatively stable over time, the reduction in *N*_*e*_ due to lifetime variance in reproductive success should not only apply to contemporary time scales but more generally throughout the coalescent time. Thus, a direct impact of life tables on genetic diversity can be expected for iteroparous species with overlapping generations.

Using both an analytical (with `AgeNe`) and a simulation-based (with `SLiM`) approach, we showed that age-specific survival and fecundity rates alone can explain a significant fraction of the variance in genetic diversity among species (Fig 3A-B). This may appear surprising at first sight, considering that we did not account for among species variation in population census sizes, which vary by several orders of magnitude in marine fishes (Hauser and Carvalho, 2008). Our results thus support that intrinsic vital rates are crucial demographic components of the neutral model to understand differences in levels of genetic diversity in marine fishes. But how generalizable is this finding to other taxa?

Age-specific survivorship curves are one of the main biological components of life tables. Three main types of survivorship curves are classically distinguished: type I curves are characterized by low juvenile and adult mortality combined with an abrupt decrease of survival when approaching the maximum age (e.g. mammals); in type II curves, survival is relatively constant during lifetime (e.g. birds) while type III curves are characterized by high juvenile mortality followed by low adult mortality (e.g. fishes and marine invertebrates). Type III survivorship curves favor the disproportionate contribution of a few lucky winners that survive to old age, compared to type I survivorship curves, where individuals have more equal contributions to reproduction, generating a lower variance in reproductive success. Thus, in type III species, higher lifetime variance in reproductive success is expected as the lifespan of a species increases. By simulating extreme type III survivorship curves (*c* = 0.1) for our 16 species while keeping their true adult lifespans, we found that 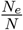 can decrease by at most 0.05 per year of lifespan (Fig 4B, extreme left). This can theoretically induce up to 60% difference in genetic diversity between the species with the shortest and the longest lifespans of our dataset. In contrast, we found no correlation between adult lifespan and 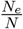 when simulating type I survivorship curves with the true lifespan values of the 16 species studied here (Fig 4B, *c* > 2), meaning than lifespan and variance in reproductive success may have limited influence in other taxonomic groups, such as birds or mammals.

Another important component of life tables is age-specific fecundity. In marine fishes, fecundity is positively correlated to female ovary size, and the relationship between fecundity and age is usually well approximated with an exponential (*F* = *aexp*^*Ab*^) or power-law (*F* = *aA*^*b*^) function. By adding an exponential increase in fecundity with age to our simulations, we found that 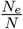 decreases even more strongly with increasing adult lifespan (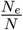 decreases by up to 0.07 per extra year of reproductive life). Using both type III survivorship and exponentially increasing fecundity with age, we could thus predict up to 84% of the variance in genetic diversity between species with the shortest and longest lifespans.

We found that 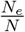 predicted from fecundity alone or age at maturity combined with age-specific survival, explained as much variation in genetic diversity as life tables with both of these components (Fig 3). This is because both of these two scenarios create sharp differences in fitness between young and old age classes. By contrast, variation in age at maturity alone (all other parameters being held constant across species) introduces some variation in 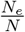 because the onset of reproduction varies from 1 to 7 years old depending on the species, but this effect is buffered by the long subsequent period during which adults will reproduce equally. Similarly, the effect of survival alone is insufficient because individuals of all species start reproducing early enough (1 year old).

Although these predicted relationships were pretty close to our empirical findings, genomewide heterozygosity decreased by about 0.09 per additional year of lifespan in our real dataset (Fig 2), which seems to be a stronger effect compared to theoretical predictions based on vital rates alone. It is thus likely that other correlates of adult lifespan and unaccounted factors also contribute to observed differences in genetic diversity among species.

### Correlated effects

When relating measures of diversity with the estimates of 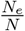 derived from life tables, we did not take into account differences in census size (*N*) between species. Population census sizes can be huge and are notoriously difficult to estimate in marine fishes. For that reason, abundance data remain largely unavailable for the 16 species of this study. We nevertheless expect long-lived species to have lower abundance compared to short-lived species because in marine fishes *N* is generally negatively correlated to body size (White et al., 2007), which is itself positively correlated to adult lifespan in our dataset (Fig S7). Hence, while we have demonstrated here that variation in vital rates has a direct effect on long-term genetic diversity, the slope between adult lifespan and genetic diversity may be inflated by uncontrolled variation in *N*. Recent genome-wide comparative studies found negative correlations between 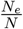 and *N* in Pinnipeds (Peart et al., 2020) as well as between genetic diversity and body size in butterflies and birds (Mackintosh et al., 2019; Brüniche-Olsen et al., 2019). Here, a highly significant negative correlation was found between genetic diversity and body size and the strength of that correlation was comparable to that found in a meta-analysis of microsatellite diversity using catch data and body size as proxies for fish abundance (Mccusker and Bentzen, 2010). We note, however, that body size was not as good a predictor of genetic diversity as lifespan and adult lifespan for the 11 non-brooding species and it was even not significant in the whole dataset of the 16 species (Table S1).

Another potentially confounding effect is the impact of r/K strategies which are the main determinant of genetic diversity across Metazoans (Romiguier et al., 2014). In our dataset, fecundity and propagule size (proxies for the r/K gradient) showed only little variance compared to their range of variation across Metazoans, and none of them were correlated to adult lifespan. However, we found that the 5 brooding species of our dataset, which are typical K-strategists, displayed lower genetic diversities with respect to their adult lifespan (Fig 2). Most interestingly, when these species were removed from the analysis, the effect of adult lifespan on genetic diversity was amplified, indicating a potentially confounding effect of parental care in marine fishes. Alternatively, low levels of genetic diversity in brooding species can also be explained by underestimated lifetime variance in reproductive success by `AgeNe` due to unaccounted variance in reproductive success within age-class. This may be particularly important in males as the age-fecundity relationship is empirically estimated for females only. This effect could be high for species with strong sexual selection and mate choice (Hastings, 1988; Naud et al., 2009). Moreover, most of these species inhabit lagoons and coastal habitats, corresponding to smaller ecological niches compared to species with no parental care, thus potentially resulting in lower long-term abundances. The discrepancy introduced by brooders in the relationship that we observed here between adult lifespan and genetic diversity may thus involve a variety of effects that remain to be elucidated.

Temporal fluctuations of effective population size may also have impacted observed levels of genetic diversity (Nei et al., 1975). All studied species possibly went through a bottleneck during the Last Glacial Maximum (Jenkins et al., 2018), which may have simultaneously decreased their genetic diversities. As the time of return to mutation-drift equilibrium is positively correlated to generation time, which is itself directly linked to adult lifespan, we may expect long-lived species to have recovered less genetic variation than short-lived species following their latest bottleneck. Moreover, long-lived species may not have recovered their pre-bottleneck population sizes as rapidly as short-lived species. If true, the negative relationship between adult lifespan and genetic diversity may be inflated compared to the sole effect of life tables.

Variation in mutation rates between species could not be accounted for due to a lack of estimates. However, if species-specific mutation rates were correlated with adult lifespan, we would expect mutation rate variation to have a direct effect on genetic diversity. Mutation rate could be linked with species life history traits through three possible mechanisms. First, the drift-barrier hypothesis predicts a negative correlation between species effective population size and the per-generation mutation rate (Sung et al., 2012). However, this hypothesis can not explain our results since species with the highest effective population sizes have the highest genetic diversity. Second, species with larger genome size tend to have more germline cell divisions, hence possibly higher mutation rates. But we did not find any correlation between genome size and genetic diversity or any other qualitative and quantitative life history traits. Third, species with longer generation time, which is positively correlated to lifespan and age at maturity, may have higher per-generation mutation rate as older individuals accumulate more germinal mutations throughout their lives. Again, under this assumption, we would expect species with longer lifespan to have higher mutation rate and genetic diversity, which goes against our observations. In summary, variation in mutation rates among species due to differences in lifespan is unlikely to explain the negative lifespan-diversity relationship we observed. If anything, variation in mutation rates should theoretically oppose this relationship.

Using one of the few direct estimates of the per-generation mutation rate in fish, Feng et al. (2017) explained the surprisingly low nucleotide diversity found in the Atlantic herring *Clupea harengus* (*π* = 0.3%) by a very low mutation rate of 2 ×10^9^ estimated from pedigree analysis. Although the herring is one of the most abundant and fecund pelagic species in the North Atlantic Ocean, its genetic diversity appears approximately 80% lower than that of the European pilchard *S. pilchardus*, another member of the *Clupeidae* family that shows the highest diversity in our study. Even if *C. harengus* has a larger body size (approximately 30 cm, compared to 20 cm for *S. pilchardus*, Froese et al. (2000)), it has above all a much longer lifespan (between 12 and 25 years) and a later age at maturity (between 2 and 6.5 years) (Jennings and Beverton, 1991). Considering even the lowest estimate of adult lifespan reported for the herring (10 years), the corresponding genetic diversity predicted by our model linking adult lifespan to genetic diversity would be around 0.5 %, which is pretty close to the empirical estimate.

Finally, we did not take into account the erosion of neutral diversity through linked selection. Addressing that issue would need to generate local estimates of nucleotide diversity and population recombination rate along the genome of each species using resequencing data aligned to a reference assembly, which was out of the scope of this study. The predicted effect of linked selection could be, however, to remove more diversity in species with large compared to small *N*_*e*_. It is therefore likely that linked selection would rather attenuate the negative relationship between adult lifespan and genetic diversity compared to neutral predictions.

## Conclusion

Here we used a simple approach to generate reference-free genome-wide estimates of diversity with *k*-mer analyses. Tested on two species with genetic diversities ranging from 0.22 to 1.42% the *k*-mer approach performed close to the level of a high-standard reference-based method in capturing fine-scale variation in diversity between evolutionary lineages and even populations of the same species. This opens the possibility to address the determinants of genetic diversity in other groups of taxa at limited costs without relying on existing genomics resources. Across Metazoans, the level of genetic diversity showed no significant relationship with the species’ conservation status (Romiguier et al., 2014). Studies performed at lower phylogenetical scales such as in Darwin’s finches and Pinnipeds, however, found reduced contemporary genetic diversity in threatened compared to non-threatened species (Brüniche-Olsen et al., 2019; Peart et al., 2020). Our results complement and extend this literature by showing the importance of taking into account life tables in comparisons of genetic diversity between species.

## Acknowledgments

The data used in this work were partly produced with the support of the GenSeq genotyping and sequencing platform, and bioinformatics data analysis benefited from the Montpellier Bioinformatics Biodiversity MBB platform, both platforms being supported by ANR program “Investissements d’avenir” (ANR-10-LABX-04-01). We would like to thank Rémy Dernat and Khalid Belkhir for their invaluable assistance in data storage, management and processing. We are grateful to the colleagues who provided us with samples as well as to those who facilitated or participated in sampling: F. Schlichta, T. Pastor, R. Castilho, R. Cunha, R. Lechuga, D. Pilo, C. Mena, J. Charton, T. Robinet, A. Darnaude, S. Vaz, M. Duranton, N. Bierne, S. Villéger, S. Blouet, as well as the fishermen and employees of fish markets and fish auctions. This work was supported by the ANR grant CoGeDiv ANR-17-CE02-0006-01. The authors declare no conflicts of interest.

## Author contributions

P.B., T.B. and P.-A.G. wrote the manuscript. P.B. and P.-A.G performed fieldwork. P.B. performed molecular experiments, and all bioinformatics and evolutionary genomics analyses with inputs from T.B. and P.-A.G. P.-A.G. conceived the project and managed financial support and genome sequencing.

## Data archiving

Data and scripts used in this study are freely available in the GitHub repository https://github.com/pierrebarry/life_tables_genetic_diversity_marine_fishes. All sampling metadata are accessible under GEOME at the CoGeDiv Project Homepage: https://geome-db.org/workbench/project-overview?projectId=357. Sequence reads have been deposited in the GenBank Sequence Read Archive under the accession code BioProject PRJNAXXXX.

## References

Benvenuto, C., Coscia, I., Chopelet, J., Sala-Bozano, M., and Mariani, S. (2017). Ecological and evolutionary consequences of alternative sex-change pathways in fish. Scientific Reports, 7(1):9084.

Brüniche-Olsen, A., Kellner, K. F., and DeWoody, J. A. (2019). Island area, body size and demographic history shape genomic diversity in Darwin’s finches and related tanagers. Molecular Ecology, 28(22):4914–4925.

Chen, J., Glémin, S., and Lascoux, M. (2017). Genetic Diversity and the Efficacy of Purifying Selection across Plant and Animal Species. Molecular Biology and Evolution, 34(6):1417–1428.

Chen, S., Zhou, Y., Chen, Y., and Gu, J. (2018). Fastp: An ultra-fast all-in-one FASTQ preprocessor. Bioinformatics, 34(17):i884–i890.

Cribari-Neto, F. and Zeileis, A. (2010). Beta Regression in R. Journal of Statistical Software, 34(1):1–24.

Crow, J. F. and Kimura, M. (1970). An Introduction to Population Genetics Theory. Harper & Row.

Curtis, J. M. R. and Vincent, A. C. J. (2006). Life history of an unusual marine fish: Survival, growth and movement patterns of Hippocampus guttulatus Cuvier 1829. Journal of Fish Biology, 68(3):707–733.

Díez-Del-Molino, D., Sánchez-Barreiro, F., Barnes, I., Gilbert, M. T. P., and Dalén, L. (2018). Quantifying Temporal Genomic Erosion in Endangered Species. Trends in Ecology & Evolution, 33(3):176–185.

Domínguez-Seoane, R., Pajuelo, J. G., Lorenzo, J. M., and Ramos, A. G. (2006). Age and growth of the sharpsnout seabream Diplodus puntazzo (Cetti, 1777) inhabiting the Canarian archipelago, estimated by reading otoliths and by backcalculation. Fisheries Research, 81(2):142–148.

Ellegren, H. and Galtier, N. (2016). Determinants of genetic diversity. Nature Reviews Genetics, 17(7):422–433.

Falconer, D. S. (1989). Introduction to Quantitative Genetics. Longman, Scientific & Technical ; Wiley, Burnt Mill, Harlow, Essex, England : New York, 3rd ed edition.

Feng, C., Pettersson, M., Lamichhaney, S., Rubin, C.-J., Rafati, N., Casini, M., Folkvord, A., and Andersson, L. (2017). Moderate nucleotide diversity in the Atlantic herring is associated with a low mutation rate. eLife, 6:e23907.

Frankham, R. (1995). Conservation Genetics. Annual Review of Genetics, 29(1):305–327.

Froese, R., Pauly, D., and Editors (2000). FishBase 2000: Concepts, design and data sources. page 344.

Gage, T. B. (2001). Age-specific fecundity of mammalian populations: A test of three mathematical models. Zoo Biology, 20(6):487–499.

Ganias, K., Somarakis, S., Machias, A., and Theodorou, A. J. (2003). Evaluation of spawning frequency in a Mediterranean sardine population (Sardina pilchardus sardina). Marine Biology, 142(6):1169–1179.

Haller, B. C. and Messer, P. W. (2017). SLiM 2: Flexible, Interactive Forward Genetic Simulations. Molecular Biology and Evolution, 34(1):230–240.

Hastings, P. A. (1988). Female choice and male reproductive success in the angel blenny, Coralliozetus angelica (Teleostei: Chaenopsidae). Animal Behaviour, 36(1):115–124.

Hauser, L. and Carvalho, G. R. (2008). Paradigm shifts in marine fisheries genetics: Ugly hypotheses slain by beautiful facts. Fish and Fisheries, 9(4):333–362.

Hedgecock, D. (1994). Does variance in reproductive success limit effective population sizes of marine organisms? In A. Genetics and Evolution of Aquatic Organisms.

Hedgecock, D. and Pudovkin, A. I. (2011). Sweepstakes Reproductive Success in Highly Fecund Marine Fish and Shellfish: A Review and Commentary. Bulletin of Marine Science, 87(4):971–1002.

Hedrick, P. (2005). Large variance in reproductive success and the Ne/N ratio. Evolution, 59(7):1596–1599.

Iglésias, S. (2013). Actinopterygians from the North-Eastern Atlantic and the Mediterranean (A Natural Classification Based on Collection Specimens, with DNA Barcodes and Standardized Photographs), Volume I (Plates), Provisional Version 09.

Jenkins, T. L., Castilho, R., and Stevens, J. R. (2018). Meta-analysis of northeast Atlantic marine taxa shows contrasting phylogeographic patterns following post-LGM expansions. PeerJ, 6:e5684.

Jennings, S. and Beverton, R. J. H. (1991). Intraspecific variation in the life history tactics of Atlantic herring (Clupea harengus L.) stocks. ICES Journal of Marine Science, 48(1):117–125.

Kimura, M. (1983). The Neutral Theory of Molecular Evolution. Cambridge University Press, Cambridge.

Kimura, M. and Crow, J. F. (1964). The Number of Alleles That Can Be Maintained in a Finite Population. Genetics, 49(4):725–738.

Kraljević, M., Matić-Skoko, S., Dul\vcić, J., Pallaoro, A., Jardas, I., and Glamuzina, B. (2007). Age and growth of sharpsnout seabream Diplodus puntazzo (Cetti, 1777) in the eastern Adriatic Sea.

Lande, R. (1995). Mutation and Conservation. Conservation Biology, 9(4):782–791.

Lande, R. and Barrowclough, G. F. (1987). Effective population size, genetic variation, and their use in population management. In Soulé, M. E., editor, Viable Populations for Conservation, pages 87–124. Cambridge University Press, Cambridge.

Leffler, E. M., Bullaughey, K., Matute, D. R., Meyer, W. K., Ségurel, L., Venkat, A., Andolfatto, P., and Przeworski, M. (2012). Revisiting an Old Riddle: What Determines Genetic Diversity Levels within Species? PLOS Biology, 10(9):e1001388.

Leroy, T., Rousselle, M., Tilak, M.-K., Caizergues, A., Scornavacca, C., Carrasco, M. R., Fuchs, J., Illera, J. C., Swardt, D. H. D., Thébaud, C., Milà, B., and Nabholz, B. (2020). Endemic island songbirds as windows into evolution in small effective population sizes. bioRxiv, page 2020.04.07.030155.

Lewontin, R. C. (1974). The Genetic Basis of Evolutionary Change. Number 25 in Columbia Biological Series. Columbia Univ. Pr, New York.

Li, H. and Durbin, R. (2009). Fast and accurate short read alignment with Burrows-Wheeler transform. Bioinformatics (Oxford, England), 25(14):1754–1760.

Louro, B., De Moro, G., Garcia, C., Cox, C. J., Veríssimo, A., Sabatino, S. J., Santos, A. M., and Canário, A. V. M. (2019). A haplotype-resolved draft genome of the European sardine (Sardina pilchardus). GigaScience, 8(5).

Mackintosh, A., Laetsch, D. R., Hayward, A., Charlesworth, B., Waterfall, M., Vila, R., and Lohse, K. (2019). The determinants of genetic diversity in butterflies. Nature Communications, 10(1):1–9.

Marçais, G. and Kingsford, C. (2011). A fast, lock-free approach for efficient parallel counting of occurrences of k-mers. Bioinformatics, 27(6):764–770.

Martinez, A. S., Willoughby, J. R., and Christie, M. R. (2018). Genetic diversity in fishes is influenced by habitat type and life-history variation. Ecology and Evolution, 8(23):12022–12031.

Mccusker, M. R. and Bentzen, P. (2010). Positive relationships between genetic diversity and abundance in fishes. Molecular Ecology, 19(22):4852–4862.

Milton, P. (1983). Biology of littoral blenniid fishes on the coast of south-west England. Journal of the Marine Biological Association of the United Kingdom, 63(1):223–237.

Murua, H. and Motos, L. (2006). Reproductive strategy and spawning activity of the European hake Merluccius merluccius (L.) in the Bay of Biscay. Journal of Fish Biology, 69(5):1288–1303.

Naud, M.-J., Curtis, J. M. R., Woodall, L. C., and Gaspar, M. B. (2009). Mate choice, operational sex ratio, and social promiscuity in a wild population of the long-snouted seahorse Hippocampus guttulatus. Behavioral Ecology, 20(1):160–164.

Nei, M., Maruyama, T., and Chakraborty, R. (1975). The Bottleneck Effect and Genetic Variability in Populations. Evolution, 29(1):1.

Nunney, L. (1991). The Influence of Age Structure and Fecundity on Effective Population Size. Proceedings: Biological Sciences, 246(1315):71–76.

Nunney, L. (1996). The influence of variation in female fecundity on effective population size. Biological Journal of the Linnean Society, 59(4):411–425.

Pauly, D., Morgan, G. R., International Center for Living Aquatic Resources Management, and Ma’had al-Kuwayt lil-Abh. āth al-’Ilmīyah, editors (1987). Length-Based Methods in Fisheries Research. Number no. 325 in ICLARM Contribution. International Center for Living Aquatic Resources Management ; Kuwait Institute for Scientific Research, Makati, Metro Manila, Philippines : Safat, Kuwait.

Peart, C. R., Tusso, S., Pophaly, S. D., Botero-Castro, F., Wu, C.-C., Aurioles-Gamboa, D., Baird, A. B., Bickham, J. W., Forcada, J., Galimberti, F., Gemmell, N. J., Hoffman, J. I., Kovacs, K. M., Kunnasranta, M., Lydersen, C., Nyman, T., de Oliveira, L. R., Orr, A. J., Sanvito, S., Valtonen, M., Shafer, A. B. A., and Wolf, J. B. W. (2020). Determinants of genetic variation across eco-evolutionary scales in pinnipeds. Nature Ecology & Evolution, pages 1–10.

Pinder, J. E., Wiener, J. G., and Smith, M. H. (1978). The Weibull Distribution: A New Method of Summarizing Survivorship Data. Ecology, 59(1):175–179.

Pinsky, M. L. and Palumbi, S. R. (2014). Meta-analysis reveals lower genetic diversity in overfished populations. Molecular Ecology, 23(1):29–39.

Poplin, R., Ruano-Rubio, V., DePristo, M. A., Fennell, T. J., Carneiro, M. O., der Auwera, G. A. V., Kling, D. E., Gauthier, L. D., Levy-Moonshine, A., Roazen, D., Shakir, K., Thibault, J., Chandran, S., Whelan, C., Lek, M., Gabriel, S., Daly, M. J., Neale, B., MacArthur, D. G., and Banks, E. (2018). Scaling accurate genetic variant discovery to tens of thousands of samples. bioRxiv, page 201178.

Ricklefs, R. E. and Miller, G. L. (1999). Ecology. W.H.Freeman & Co Ltd, New York, 4th edition edition.

Rolland, J., Schluter, D., and Romiguier, J. (2020). Vulnerability to Fishing and Life History Traits Correlate with the Load of Deleterious Mutations in Teleosts. Molecular Biology and Evolution.

Romiguier, J., Gayral, P., Ballenghien, M., Bernard, A., Cahais, V., Chenuil, A., Chiari, Y., Dernat, R., Duret, L., Faivre, N., Loire, E., Lourenco, J. M., Nabholz, B., Roux, C., Tsagkogeorga, G.a. T., Weber, A., Weinert, L. A., Belkhir, K., Bierne, N., Glémin, S., and Galtier, N. (2014). Comparative population genomics in animals uncovers the determinants of genetic diversity. Nature, 515(7526):261–263.

Sung, W., Ackerman, M. S., Miller, S. F., Doak, T. G., and Lynch, M. (2012). Drift-barrier hypothesis and mutation-rate evolution. Proceedings of the National Academy of Sciences, 109(45):18488–18492.

Tine, M., Kuhl, H., Gagnaire, P.-A., Louro, B., Desmarais, E., Martins, R. S. T., Hecht, J., Knaust, F., Belkhir, K., Klages, S., Dieterich, R., Stueber, K., Piferrer, F., Guinand, B., Bierne, N., Volckaert, F. A. M., Bargelloni, L., Power, D. M., Bonhomme, F., Canario, A. V. M., and Reinhardt, R. (2014). European sea bass genome and its variation provide insights into adaptation to euryhalinity and speciation. Nature Communications, 5:5770.

Tsikliras, A. C. and Stergiou, K. I. (2015). Age at maturity of Mediterranean marine fishes. Mediterranean Marine Science, 16(1):5–20.

Väli, Ü., Einarsson, A., Waits, L., and Ellegren, H. (2008). To what extent do microsatellite markers reflect genome-wide genetic diversity in natural populations? Molecular Ecology, 17(17):3808–3817.

Vurture, G. W., Sedlazeck, F. J., Nattestad, M., Underwood, C. J., Fang, H., Gurtowski, J., and Schatz, M. C. (2017). GenomeScope: Fast reference-free genome profiling from short reads. Bioinformatics, 33(14):2202–2204.

Waples, R. S. (1991). Heterozygosity and Life-History Variation in Bony Fishes: An Alternative View. Evolution, 45(5):1275–1280.

Waples, R. S. (2002). Evaluating the effect of stage-specific survivorship on the Ne/N ratio. Molecular Ecology, 11(6):1029–1037.

Waples, R. S. (2016a). Life-history traits and effective population size in species with overlapping generations revisited. Heredity, 117(4):241–250.

Waples, R. S. (2016b). Tiny estimates of the Ne/N ratio in marine fishes: Are they real? Journal of Fish Biology, 89(6):2479–2504.

Waples, R. S., Do, C., and Chopelet, J. (2011). Calculating Ne and Ne/N in age-structured populations: A hybrid Felsenstein-Hill approach. Ecology, 92(7):1513–1522.

Waples, R. S., Luikart, G., Faulkner, J. R., and Tallmon, D. A. (2013). Simple life-history traits explain key effective population size ratios across diverse taxa. Proceedings of the Royal Society B: Biological Sciences, 280(1768).

Waples, R. S., Mariani, S., and Benvenuto, C. (2018). Consequences of sex change for effective population size. Proceedings of the Royal Society B: Biological Sciences, 285(1893):20181702.

White, E. P., Ernest, S. K. M., Kerkhoff, A. J., and Enquist, B. J. (2007). Relationships between body size and abundance in ecology. Trends in Ecology & Evolution, 22(6):323–330.

Wright, S. (1969). The Theory of Gene Frequencies. Number Sewall Wright ; Vol. 2 in Evolution and the Genetics of Populations. Univ. of Chicago Press, Chicago, Ill., paperback ed edition.

Zeileis, A. and Hothorn, T. (2002). Diagnostic checking in regression relationships. R News, 2(3):7–10.

