## Supplementary Material for "Age-specific survivorship and fecundity shape genetic diversity in marine fishes"

**Additional tables** **6**

**Additional figures** **7**

### Additional methods

#### Sampling, DNA extraction, whole-genome sequencing and reads quality control

We sampled 16 marine teleostean fish species presenting a wide diversity of life history strategies expected to affect genetic diversity. For 12 of these species, 20 individuals were sampled (5 per location). For the 4 other species, the total number of samples ranged from 10 to 19 (Table 1). Individuals were either sampled from landings in local fish markets, captured in the field (using hand nets, lure fishing, spearfishing or beach seines) or provided by collaborators. The majority of the sampling was done in 2018 and 2019. Whole-genomic DNA was extracted from fin or tissue clips stored in 95% ethanol using the NucleoSpin Tissue Kit (Macherey-Nagel) and treated with RNAse A to remove residual RNA. Double-stranded nucleic acid concentration was quantified using Qubit2.0 and standardized to 20ng per $\mu$l. Individual whole-genome sequencing libraries were prepared following the Illumina TruSeq DNA PCR-Free Protocol and sequenced by Genewiz Inc (USA). Libraries were quantified and multiplexed by groups of 40 individuals and sequenced on two S4 flow cells on a NovaSeq6000 instrument (Illumina) to generate 150 bp paired-end reads, targetting an average read depth of 20X per individual. Raw reads were preprocessed with `fastp` v.0.20.0 (Chen et al., 2018) using default parameters, allowing quality control, filtering by quality, length and complexity, and adapter trimming to be performed in a single step. Base correction was performed using a quality comparison between overlapping bases of paired-end reads, and polyG tail trimming was enabled to correct for artefactual G repetitions occurring in Novaseq read tails.

#### Collection of life history traits database

As growth is indeterminate in fish, we defined adult body size as the infinite length, $L_{inf}$ determined by the Von Bertalanffy equation ($L_t = L_{inf}[1 - exp^{-K(t-t0)}]$), that links individual body size $L_t$ to age $t$, with $K$ a parameter defining the shape of this relationship (Pauly et al., 1987). We estimated adult body size as the median of all $L_{inf}$ values reported for each species in the online database Fishbase (Froese et al., 2000). As $L_{inf}$ was not documented in Fishbase for *D. puntazzo* and *C. galerita*, we took the median of the values reported in (Kraljević et al., 2007) and (Domínguez-Seoane et al., 2006) for *D. puntazzo*, and the maximum length observed in (Milton, 1983) for *C. galerita*. Trophic level was retrieved from Fishbase. Fecundity was defined as the absolute fecundity, i.e. the mean number of eggs in an ovary of a female in a single spawning event. Females may spawn several times during one reproductive season (Ganias et al., 2003; Murua and Motos, 2006), so absolute fecundity is not the value most directly relevant to global genetic diversity. However, it is the most commonly reported in the literature as the number of spawnings events per reproductive season is difficult to measure. Because fecundity is proportional to individual body size, we computed fecundity at infinite length, $L_{inf}$. Propagule size was determined following Romiguier et al. (2014), as the size of the dispersal stage that becomes independent of the parents. For all species of this study, this corresponded to egg diameter, except for brooders, for which we used hatching size. All propagule size data were retrieved from species-specific references. Age at maturity was defined as the age at which 50% of the population is mature. Values for age at maturity were taken from Tsikliras and Stergiou (2015) for seven species while other values were retrieved from species-specific references. Likewise, lifespan values were taken from (Tsikliras and Stergiou, 2015) for six species and completed with specific references. Finally, adult lifespan was defined as $Lifespan - Age\ at\ Maturity$ (Waples et al., 2013).

#### Estimation of genetic diversity with GenomeScope

Provided a sufficient average coverage depth (e.g. 20X), `GenomeScope` evaluates the fraction of heterozygous sites from the ratio of the height of the heterozygous to the homozygous $k$-mer peak, occurring at 50% (i.e. 10X) and 100% (i.e. 20X) of the average coverage depth, respectively. The number of different possible $k$-mers (and thus the precision of the method) increases with $k$, but so does the runtime and the probability of "wrong" $k$-mers due to sequencing errors. We set $k = 21$ as recommended by `GenomeScope` and performed a sensitivity analysis by estimating genetic diversity and genome size for one individual of *D. labrax* using $k$ from 17 to 25 (Fig S4).

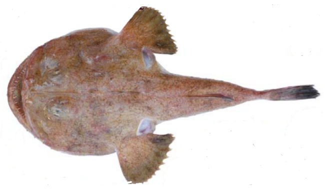

*Lophius budegassa*:
high genome size and
low diversity (0.225 %)

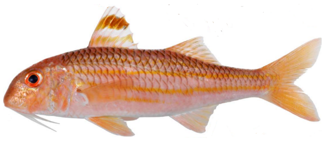

*Mullus surmuletus*:
low genome size and
high diversity (1.135 %)

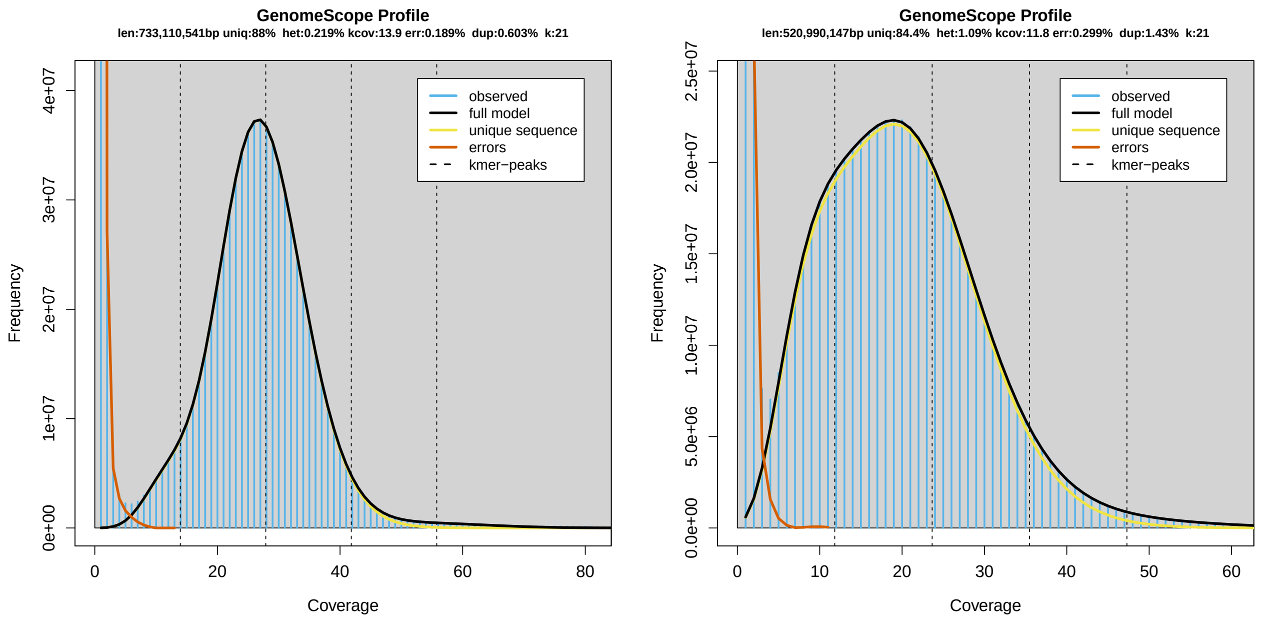

Figure S1: $k-mer$ **frequency-coverage relationship and estimation by** `GenomeScope` **v.1.0 (Vurture et al., 2017) for two species, *L. budegassa* and *M.surmuletus***

In order to assess the reliability of `GenomeScope`, the 20 resequenced genomes for two species (*S. pilchardus* and *D. labrax*) were aligned with `bwa-mem` v.0.7.17 (Li and Durbin, 2009) to the reference genomes retrieved from Louro et al. (2019) and Tine et al. (2014) for *S. pilchardus* and *D. labrax*, respectively. We then removed PCR duplicates with the Picard tools `MarkDuplicates` v.2.23.2. We followed the best-practice pipeline in `GATK` v.4.1.6.0 for variant calling (Poplin et al., 2018): we ran `HaplotypeCaller` with default options to generate individual GVCFs files, stored them in a database with `GenomicsDBImport` and finally computed VCF files with `GenotypeGVCFs`. We didn't apply post variant calling filtering steps, such as hard filters on genotype quality scores or Hardy-Weinberg Equilibrium criterion, in order to avoid potential bias in the comparison of genetic diversity between species with very different rates of heterozygosity. However, we assume that possible bias due to the absence of variant filtering should not impact differences among individuals within each species. However, we generated VCF files with all sites, including non-variant ones, to avoid underestimation of genetic diversity due to the assumption that missing sites are homozygous for the reference allele. We estimated species genetic diversity as:

$$\pi = \frac{\sum^{L} \sum_{i<j} k_{ij}}{\sum^{L} \binom{n}{2}} \tag{S1}$$

where $\sum_{i<j} k_{ij}$ is the number of pairwise nucleotide differences between all haplotypes at a given site, $\binom{n}{2}$ is the total number of pairwise nucleotide comparison between all haplotypes at a given site, and $L$, the number of sites. We thus exclude missing data from the analysis. However, we include multiallelic sites as removing these sites may underestimate genetic diversity, especially for species with high genetic diversity.

#### Forward simulations

We used `SLiM` v.3.3.1 (Haller and Messer, 2017) to perform forward simulations to estimate genetic diversity at mutation-drift equilibrium incorporating age-specific survival and fecundity. For each individual, the number of offspring produced per year was determined by a Poisson distribution with mean $\lambda_{S,a}$ specific to each species and each age. Age at first maturity was set to 1 for all simulations. To keep population size constant in these non Wright-Fisher forwards simulations, we introduced a carrying capacity parameter, allowing population size to fluctuate around this capacity (Fig S17 - S24). We arbitrarily set this parameter to $N = 2000$, and simulated non-recombining 1Mb loci with a mutation rate of $\mu = 1e^{-7}$. Each simulation was run for 25000 years, which was long enough for genetic diversity to reach mutation-drift equilibrium (Fig S25 - S32). For each simulation, we estimated the mean genetic diversity (i.e., the proportion of heterozygous sites along the 1Mb locus) over the last 10000 years after checking that an equilibrium has been reached. For each species, we ran 50 replicates and defined the genetic diversity predicted by a given simulation scenario as the mean genetic diversity at equilibrium averaged over the 50 replicates.

#### Evaluating the impact of life tables beyond marine fish

We explored the impact of alternative fecundity-age models on the relationship between adult lifespan and $\frac{N_e}{N}$ using three additional biologically realistic models: linear ($F_{Age} = a \times Age + b$), polynomial ($F_{Age} = [Age - AgeMat][(AgeMat + Lifespan - Age)^2]$, common in mammal) (Gage, 2001) and power-law ($F_{Age} = Age^f$). For the linear and the polynomial model, $f$ describes the maximum fecundity at lifespan and age with the highest fecundity, respectively (i.e. higher absolute values of $f$ correspond to higher differences in fecundity between low and high fecund ages for both models). $f$ was between -1 and 1 for the linear and the polynomial model. For the power-law model, we took values of $f$ from $-5$ to 5.

### Additional results

#### Whole-genome resequencing data set

We resequenced 300 individual genomes from 16 marine teleostean species, generating from $59.86 \times 10^6$ to $200.92 \times 10^6$ reads per individual (mean = $129 \times 10^6$, sd = $20 \times 10^6$, Fig S2). The read quality score (Q30 rate) ranged between 88% and 94% (mean = 92.4%, sd = 1.1) and the duplication rate lied between 5 and 15% (mean = 10.8%, sd = 2.6) (Fig S2). GC content was moderately variable among species and highly consistent among individuals of the same species, except for one individual of *S. cabrilla*, *D. puntazzo* and *M. surmuletus* that showed a marked discrepancy with the overall GC content of their species (Fig S2). These three individuals were thus removed from downstream analyses to avoid potential issues due to contamination or poor sequencing quality (see discussion).

#### Validation of GenomeScope estimations

Although, we detected slight differences in within-species genetic diversity between individuals of different basins that can be divided in two clusters: the first one included 9 species with generally lower genetic diversity in the Mediterranean than Atlantic localities, while the opposite was observed in the second cluster (7 species) (Fig 1C). The species of the second cluster are often found in coastal habitats, lagoons, estuaries whereas species of the first cluster are rather pelagic, epi-pelagic or benthic species. The only exception was the presence of *H. guttulatus* in the Atlantic cluster.

### Additional tables

| Dataset | Predictor | *p-value* | Pseudo $R^2$ | Slope estimate ($\pm$ 95% interval) |
|---|---|---|---|---|
| Whole data set | Body size | 0.119 | 0.192 | $-0.006(-0.014; 0.002)$ |
| | Trophic level | 0.676 | 0.012 | $-0.091(-0.524; 0.343)$ |
| | Propagule size | 0.562 | 0.015 | $-0.014(-0.062; 0.034)$ |
| | Fecundity | 0.653 | 0.013 | $-1.22e^{-5}(-6.63e^{-5}; 4.20e^{-5})$ |
| | Lifespan | **0.0107** | 0.438 | $-0.062(-0.111; -0.013)$ |
| | Adult lifespan | **0.0070** | 0.429 | $-0.089(-0.156; -0.023)$ |
| | Hermaphroditism | 0.434 | 0.034 | $0.1779(-0.278; 0.633)$ |
| | Parental Care | 0.274 | 0.075 | $-0.273(-0.772; 0.226)$ |
| No parental care | Body size | $\mathbf{6.60e^{-5}}$ | 0.616 | $-0.014(-0.021; -0.007)$ |
| | Trophic level | 0.256 | 0.093 | $-0.326(-0.902; 0.251)$ |
| | Propagule size | 0.170 | 0.175 | $-0.518(-1.273; 0.237)$ |
| | Fecundity | 0.390 | 0.056 | $-2.51e^{-5}(-8.35e^{-5}; 3.33e^{-5})$ |
| | Lifespan | $\mathbf{1.017e^{-7}}$ | 0.851 | $-0.095(-0.131; -0.060)$ |
| | Adult lifespan | $\mathbf{1.65e^{-7}}$ | 0.829 | $-0.129(-0.179; -0.080)$ |
| | Hermaphroditism | 0.454 | 0.044 | $0.206(-0.345; 0.757)$ |

Table S1: **Statistical relationships between species genetic diversity and life history traits** - Genetic diversity was fitted to 6 quantitative (body size, trophic level, propagule size, fecundity, lifespan and adult lifespan) and two qualitative predictors (hermaphroditism and parental care) with a beta regression model using the `betareg` R package (Zeileis and Hothorn, 2002). In the upper part of the table, regressions were performed with the whole dataset, while in the lower part only the 11 non-brooding species were considered.

Table S2: **Mapping and variant calling statistics for *D. labrax* and *S. pilchardus* individuals** - For each, individual, number and percentage of reads mapped with `bwa-mem` v.0.7.17 (Li and Durbin, 2009). Individual GCVF files were created from bam files with `HaplotypeCaller` from `GATK` v.4.1.6.0 (Poplin et al., 2018), then stored in a GCVF database with `GenomicsDBImport`, and VCF files were finally generated with `GenotypeGVCFs`. Individual heterozygosity was estimated with a custom script. $O_{het}$ is the number of observed heterozygous positions at $N_{sites}$ number of variable sites. $Het_{bwa+GATK}$ and $Het_{GenomeScope}$ correspond to genome-wide average heterozygosity estimated with the variant calling and `GenomeScope` approaches respectively. Sample names indicate geographical origin (Li = Gulf of Lion, Mu = Costa Calida, Fa = Algarve, Ga = Bay of Biscay).

| Species | Sample | Reads mapped | % reads mapped | $O_{het}$ | $N_{sites}$ | $Het_{bwa+GATK}$ | $Het_{GenomeScope}$ |
|---|---|---|---|---|---|---|---|
| *D. labrax* | DlabrFa1 | 113 750 879 | 98.50 | 2 147 840 | 503 641 870 | 0.4264618 | 0.4023185 |
| | DlabrFa3 | 113 675 336 | 98.47 | 2 208 200 | 503 847 656 | 0.4382674 | 0.4056315 |
| | DlabrFa4 | 119 760 663 | 98.26 | 2 168 714 | 503 981 403 | 0.4303163 | 0.3963415 |
| | DlabrFa5 | 111 632 052 | 98.40 | 2 280 744 | 503 764 199 | 0.4527404 | 0.4370325 |
| | DlabrFa6 | 118 251 538 | 98.39 | 2 153 188 | 503 902 634 | 0.4273024 | 0.391308 |
| | DlabrMu1 | 111 356 113 | 98.20 | 1 719 388 | 502 790 222 | 0.3419693 | 0.2962895 |
| | DlabrMu2 | 78 923 334 | 96.72 | 1 930 312 | 498 084 060 | 0.3875474 | 0.3733955 |
| | DlabrMu3 | 100 325 589 | 98.00 | 1 874 416 | 502 625 989 | 0.3729246 | 0.332756 |
| | DlabrMu4 | 113 701 410 | 98.29 | 1 955 294 | 503 912 767 | 0.3880223 | 0.355843 |
| | DlabrMu6 | 117 131 984 | 98.11 | 1 923 386 | 503 422 444 | 0.3820620 | 0.3329545 |
| | DlabrLi1 | 100 519 118 | 98.16 | 1 919 555 | 503 396 155 | 0.3813209 | 0.3627435 |
| | DlabrLi2 | 103 717 839 | 98.18 | 1 926 045 | 503 259 727 | 0.3827139 | 0.3566875 |
| | DlabrLi3 | 101 815 078 | 98.18 | 1 917 988 | 503 399 019 | 0.3810075 | 0.3713805 |
| | DlabrLi4 | 94 399 356 | 98.21 | 1 920 186 | 502 848 166 | 0.3818620 | 0.3520245 |
| | DlabrLi5 | 115 515 514 | 98.06 | 1 918 720 | 503 724 011 | 0.3809070 | 0.354731 |
| | DlabrGa2 | 129 379 987 | 98.45 | 2 114 798 | 503 898 866 | 0.4196870 | 0.3770105 |
| | DlabrGa3 | 118 863 303 | 98.02 | 2 082 274 | 503 524 710 | 0.4135396 | 0.388246 |
| | DlabrGa4 | 125 269 167 | 98.02 | 2 085 977 | 503 675 656 | 0.4141508 | 0.380342 |
| | DlabrGa5 | 121 308 016 | 98.05 | 2 093 356 | 503 693 457 | 0.4156012 | 0.3807775 |
| | DlabrGa6 | 113 020 888 | 98.04 | 2 150 834 | 503 563 171 | 0.4271230 | 0.379656 |
| *S. pilchardus* | SpilcFa1 | 98 789 082 | 96.42 | 7 128 735 | 604 856 771 | 1.178582 | 1.331 |
| | SpilcFa3 | 109 782 816 | 96.24 | 7 452 162 | 613 645 048 | 1.214409 | 1.340575 |
| | SpilcFa4 | 97 444 013 | 95.07 | 7 195 048 | 609 516 704 | 1.180451 | 1.347235 |
| | SpilcFa5 | 110 357 654 | 95.67 | 7 545 357 | 615 252 143 | 1.226385 | 1.40959 |
| | SpilcFa6 | 104 223 183 | 95.90 | 7 500 848 | 615 157 704 | 1.219337 | 1.41341 |
| | SpilcMu1 | 124 885 493 | 96.20 | 7 471 544 | 617 558 276 | 1.209852 | 1.500415 |
| | SpilcMu2 | 109 948 539 | 95.70 | 7 272 037 | 615 199 861 | 1.182061 | 1.474785 |
| | SpilcMu3 | 108 564 997 | 92.46 | 7 272 929 | 614 357 522 | 1.183827 | 1.537 |
| | SpilcMu4 | 122 541 302 | 95.47 | 7 283 499 | 616 999 758 | 1.180470 | 1.446285 |
| | SpilcMu6 | 105 613 487 | 96.25 | 7 172 809 | 614 547 029 | 1.167170 | 1.40057 |
| | SpilcLi2 | 121 934 021 | 95.06 | 7 123 340 | 614 907 641 | 1.158441 | 1.373815 |
| | SpilcLi3 | 126 015 301 | 95.03 | 7 581 636 | 617 017 545 | 1.228755 | 1.417255 |
| | SpilcLi4 | 116 445 615 | 95.68 | 7 350 852 | 615 226 870 | 1.194820 | 1.37415 |
| | SpilcLi5 | 119 470 404 | 95.48 | 7 097 138 | 612 517 142 | 1.158684 | 1.336275 |
| | SpilcLi6 | 112 805 605 | 96.10 | 6 835 691 | 611 663 555 | 1.117557 | 1.279095 |
| | SpilcGa1 | 99 664 874 | 93.16 | 7 195 528 | 612 305 296 | 1.175154 | 1.5632 |
| | SpilcGa3 | 106 538 562 | 95.80 | 7 353 467 | 615 590 542 | 1.194539 | 1.65899 |
| | SpilcGa4 | 97 977 975 | 90.70 | 6 894 734 | 609 669 475 | 1.130897 | 1.895515 |
| | SpilcGa5 | 102 032 394 | 95.10 | 7 137 835 | 612 596 676 | 1.165177 | 1.592515 |
| | SpilcGa6 | 97 318 612 | 87.45 | 7 129 996 | 611 349 731 | 1.166271 | 1.97076 |

| Dataset | Predictor | *p-value* | | | Pseudo $R^2$ | | | Slope estimate (± 95% interval) | | |
|---|---|---|---|---|---|---|---|---|---|---|
| | | All | Med. | Atl. | All | Med. | Atl. | All. | Med. | Atl. |
| Whole data set | Body size | 0.119 | 0.105 | 0.159 | 0.192 | 0.206 | 0.169 | −0.006 (−0.014; 0.002) | −0.006 (−0.014; 0.001) | −0.005 (−0.013; 0.002) |
| | Trophic level | 0.676 | 0.671 | 0.672 | 0.012 | 0.012 | 0.012 | −0.091 (−0.524; 0.343) | −0.094 (−0.536; 0.349) | −0.090 (−0.516; 0.336) |
| | Propagule size | 0.562 | 0.608 | 0.450 | 0.015 | 0.012 | 0.032 | −0.014 (−0.062; 0.034) | −0.013 (−0.061; 0.036) | −0.018 (−0.066; 0.030) |
| | Fecundity | 0.653 | 0.605 | 0.723 | 0.013 | 0.017 | 0.008 | $-1.22e^{-5}$ ($-6.63e^{-5}; 4.20e^{-5}$) | $-1.44e^{-5}$ ($-7.00e^{-5}; 4.13e^{-5}$) | $-9.2e^{-6}$ ($-6.18e^{-5}; 4.35e^{-5}$) |
| | Lifespan | **0.0107** | **0.0085** | **0.0219** | 0.438 | 0.453 | 0.404 | −0.062 (−0.111; −0.013) | −0.0652 (−0.115; −0.016) | −0.056 (−0.104; −0.007) |
| | Adult lifespan | **0.0070** | **0.0055** | **0.0216** | 0.429 | 0.444 | 0.374 | −0.089 (−0.156; −0.023) | −0.093 (−0.161; −0.026) | −0.077 (−0.144; −0.0010) |
| | Hermaphroditism | 0.434 | 0.465 | 0.350 | 0.034 | 0.03 | 0.050 | 0.1779 (−0.278; 0.633) | 0.1701 (−0.296; 0.636) | 0.2069 (−0.2359; 0.6497) |
| | Parental Care | 0.274 | 0.287 | 0.168 | 0.075 | 0.071 | 0.121 | −0.273 (−0.772; 0.226) | −0.271 (−0.781; 0.239) | −0.335 (−0.822; 0.152) |
| No parental care | Body size | $\mathbf{6.60e^{-5}}$ | $\mathbf{3.85e^{-5}}$ | $\mathbf{8.17e^{-5}}$ | 0.616 | 0.636 | 0.607 | −0.014 (−0.021; −0.007) | −0.014 (−0.021; −0.007) | −0.014 (−0.021; −0.007) |
| | Trophic level | 0.256 | 0.245 | 0.254 | 0.093 | 0.097 | 0.095 | −0.326 (−0.902; 0.251) | −0.340 (−0.925; 0.244) | −0.328 (−0.903; 0.247) |
| | Propagule size | 0.170 | 0.155 | 0.200 | 0.175 | 0.184 | 0.157 | −0.518 (−1.273; 0.237) | −0.544 (−1.310; 0.221) | −0.484 (−1.240; 0.272) |
| | Fecundity | 0.390 | 0.353 | 0.401 | 0.056 | 0.065 | 0.054 | $-2.51e^{-5}$ ($-8.35e^{-5}; 3.33e^{-5}$) | $-2.76e^{-5}$ ($-8.71e^{-5}; 3.19e^{-5}$) | $-2.44e^{-5}$ ($-8.26e^{-5}; 3.37e^{-5}$) |
| | Lifespan | $\mathbf{1.017e^{-7}}$ | $\mathbf{7.28e^{-8}}$ | $\mathbf{5.451e^{-7}}$ | 0.851 | 0.856 | 0.832 | −0.095 (−0.131; −0.060) | −0.097 (−0.134; −0.061) | −0.094 (−0.131; −0.056) |
| | Adult lifespan | $\mathbf{1.65e^{-7}}$ | $\mathbf{1.36e^{-7}}$ | $\mathbf{8.778e^{-7}}$ | 0.829 | 0.837 | 0.805 | −0.129 (−0.179; −0.080) | −0.131 (−0.181; −0.081) | −0.127 (−0.178; −0.075) |
| | Hermaphroditism | 0.454 | 0.407 | 0.489 | 0.044 | 0.053 | 0.038 | 0.206 (−0.345; 0.757) | 0.231 (−0.326; 0.787) | 0.191 (−0.360; 0.741) |

Table S3: **Statistical relationships between different estimations of species genetic diversity and life history traits** - Genetic diversity, either estimated from all individuals (all), individuals from Mediterranean Sea (Med.), Atlantic Ocean (Atl.) was fitted to 6 quantitative (body size, trophic level, propagule size, fecundity, lifespan and adult lifespan) and two qualitative predictors (hermaphroditism and parental care) with a beta regression model using the `betareg` R package (Zeileis and Hothorn, 2002). In the upper part of the table, regressions were performed with the whole dataset, while in the lower part only the 11 non-brooding species were considered.

**Table S4 - Life-history traits and observed genetic diversity of the 16 teleostean marine species.** - For each species, number of individuals used for the estimation of genetic diversity ; observed median genetic diversity among all individuals (± standard deviation) ; body size (in centimeters); trophic level; age at first maturity (in years), lifespan (in years), adult lifespan (in years, defined as the difference between lifespan and age at maturity), parental care behaviour (– = no eggs protection ; NG = nest-guarders ; MP = male-pooch) and hermaphroditism (– = no hermaphroditism ; PG = protogynous ; PA = protandrous, RUD = rudimentary).

| Species | Vernacular name | N. | Genetic diversity (%) | Body size (cm) | Trophic level | Fecundity | Propagule Size (mm) | Maturity (years) | Lifespan (years) | Adult lifespan (years) | Parental care | Herma. |
|---|---|---|---|---|---|---|---|---|---|---|---|---|
| *Coryphoblennius galerita* | Montagu's blenny | 16 | 0.607(±0.014) | 7[17] | 2.28[18] | NA | 3.3[41] | 1.5[32] | 6[17] | 4.5[17,32] | NG[1] | – |
| *Coris julis* | Rainbow wrasse | 20 | 1.172(±0.056) | 27.2[18] | 3.24[18] | 169.81[44] | 0.63[11] | 1[16] | 7[21] | 6[16,21] | –[16] | PG[29] |
| *Dicentrarchus labrax* | European sea bass | 20 | 0.375(±0.031) | 102.15[18] | 3.47[18] | 12436.52[22] | 1.15[7] | 3[47] | 15[47] | 12[47] | –[18] | – |
| *Diplodus puntazzo* | Sharp-snout seabream | 19 | 0.533(±0.074) | 49.69[27] | 3.07[18] | 277.87[46] | 0.87[25] | 2[25] | 10[27] | 8[25,27] | –[49] | RUD[24] |
| *Hippocampus guttulatus* | Long-snouted seahorse | 12 | 0.313(±0.090) | 19.8[18] | 3.5[18] | 1.21[12] | 12[30] | 0.5[12] | 5[12] | 4.5[12] | MP[23] | – |
| *Lophius budegassa* | Blackbellied angler | 20 | 0.225(±0.015) | 103[18] | 4.23[18] | 2304.03[9] | 1.88[8] | 7.5[14] | 21[28] | 13.5[28,14] | –[18] | – |
| *Lithognathus mormyrus* | Striped seabream | 20 | 0.553(±0.027) | 37.85[18] | 3.42[18] | 214.09[15] | 0.75[13] | 2[47] | 12[47] | 10[47] | –[3] | PA[16] |
| *Merluccius merluccius* | European hake | 20 | 0.844(±0.025) | 88.9[18] | 4.43[18] | 2294.54[36] | 1.07[5] | 3[47] | 11[35] | 8[35,47] | –[35] | – |
| *Mullus surmuletus* | Striped red mullet | 19 | 1.135(±0.048) | 30.18[18] | 3.46[18] | 2569.32[2] | 0.86[41] | 1.5[47] | 6[47] | 4.5[47] | –[16] | – |
| *Pagellus erythrinus* | Common pandora | 19 | 1.100(±0.020) | 36[18] | 3.46[18] | 2280.46[40] | 0.77[26] | 2[47] | 8[47] | 6[47] | –[3] | PG[16] |
| *Serranus cabrilla* | Comber | 19 | 1.205(±0.055) | 30.8[18] | 3.68[18] | 37.97[39] | 0.91[41] | 2[19] | 6[48] | 4[48,19] | –[24] | – |
| *Spondyliosoma cantharus* | Black seabream | 19 | 0.478(±0.034) | 35.7[18] | 3.27[18] | 425.62[20] | 2.1[41,43] | 3[33] | 10[34] | 7[33,34] | NG[16] | PG[16] |
| *Symphodus cinereus* | Grey wrasse | 10 | 0.660(±0.125) | 14.1[18] | 3.3[18] | 13.20[23] | 2.87[23] | 1.5[16] | 6[23] | 4.5[23,16] | NG[16,23] | – |
| *Sardina pilchardus* | European pilchard | 20 | 1.415(±0.182) | 20.35[18] | 2.94[18] | 22.89[6] | 1.64[10] | 1[47] | 5[47] | 4[47] | –[16] | – |
| *Syngnathus typhle* | Broadnosed pipefish | 20 | 0.859(±0.047) | 26.2[18] | 3.75[18] | 0.38[42] | 20[23] | 1[4] | 3[45] | 2[45] | MP[23] | – |
| *Sarda sarda* | Atlantic bonito | 20 | 0.896(±0.208) | 68.9[18] | 4.34[18] | 15647.73[37] | 1.3[38] | 1[47] | 4[47] | 3[47] | –[31] | – |

### References

[1] Almada, V. C., Carreiro, H., Faria, C., and Gonçalves, E. J. (1996). The breeding season of Coryphoblennius galerita in Portuguese waters. *Journal of Fish Biology*, 48(2):295–297.

[2] Amin, A., Madkour, F., Abu El-Regal, M., and Moustafa, A. (2016). Reproductive biology of Mullus surmuletus (Linnaeus, 1758) from the Egyptian Mediterranean Sea (Port Said. *iNTERNATIONAL jOURNAL OF eNVIRONMENTAL sCIENCE and engineering*, 7:1–10.

[3] Benvenuto, C., Coscia, I., Chopelet, J., Sala-Bozano, M., and Mariani, S. (2017). Ecological and evolutionary consequences of alternative sex-change pathways in fish. *Scientific Reports*, 7(1):9084.

[4] Bernet, P., Rosenqvist, G., and Berglund, A. (1998). Female-Female Competition Affects Female Ornamentation in the Sex-Role Reversed Pipefish Syngnathus typhle. *Behaviour*, 135(5):535–550.

[5] Bjelland, R. M. and Skiftesvik, A. B. (2006). Larval development in European hake (Merluccius merluccius L.) reared in a semi-intensive culture system. *Aquaculture Research*, 37(11):1117–1129.

[6] Bouhali, F., Lechekhab, S., Ladaimia, S., Assia, B., Amara, R., and Borhane, D. (2015). Reproduction et maturation des gonades de Sardina pilchardus dans le golfe d'Annaba (Nord-Est algérien). *Cybium: international journal of ichthyology*.

[7] Cerdá, J., Carrillo, M., Zanuy, S., Ramos, J., and de la Higuera, M. (1994). Influence of nutritional composition of diet on sea bass, Dicentrarchus labrax L., reproductive performance and egg and larval quality. *Aquaculture*, 128(3):345–361.

[8] Colmenero, A. (2017). *Towards Biological and Ecological Knowledge of Lophius Spp. in the NW Mediterranean Sea for a Sustainable Fishery*. PhD thesis, Universidad de Barcelona.

[9] Colmenero, A. I., Tuset, V. M., Recasens, L., and Sanchez, P. (2013). Reproductive biology of Black Anglerfish (Lophius budegassa) in the northwestern Mediterranean Sea. *Fishery Bulletin*, 111(4):390–401.

[10] Coombs, S., Boyra, G., Rueda, L., Uriarte, A., Santos, M., Conway, D., and Halliday, N. (2004). Buoyancy measurements and vertical distribution of eggs of sardine ( Sardina pilchardus) and anchovy ( Engraulis encrasicolus). *Marine Biology*, 145:959–970.

[11] Crec'hriou, R., Marinaro, J.-Y., and Planes, S. (2015). Advance in identification of pelagic eggs of mediterranean fish: Development of a new identification key. *Vie et Milieu*, 65:47–61.

[12] Curtis, J. M. R. and Vincent, A. C. J. (2006). Life history of an unusual marine fish: Survival, growth and movement patterns of Hippocampus guttulatus Cuvier 1829. *Journal of Fish Biology*, 68(3):707–733.

[13] DIVANACH, P. and KENTOURI, M. (1983). Données préliminaires sur les caractéristiques du développement embryonnaire et larvaire du marbre Lithognathus mormyrus en élevage extensif. *Données préliminaires sur les caractéristiques du développement embryonnaire et larvaire du marbre Lithognathus mormyrus en élevage extensif*, 7(4):89–103.

[14] Duarte, R., Azevedo, M., Landa, J., and Pereda, P. (2001). Reproduction of anglerfish (Lophius budegassa Spinola and Lophius piscatorius Linnaeus) from the Atlantic Iberian coast. *Fisheries Research*, 51(2):349–361.

[15] Faraj, E., Alssalam, A., Ali, S., Sayed, M., Sayed, E., Mor, E., Ali, R., Ali, S., Salem, E., and Alfergani, E. (2016). Reproductive Biology of the Striped Seabream Lithognathus mormyrus (Linnaeus, 1758) from Al Haneah Fishing Site, Mediterranean Sea, Eastern Libya. *Journal of Life Sciences*, 10.

[16] Fischer, W., Bauchot, M.-L., and Schneider, M. (1987). *Fiches FAO d'identification Desespèces Pour Les Besoins de La Pêche.(Révision 1). Méditerranée et Mer Noire.Zone de Pêche 37. Volume II. Vertébrés.Publication Préparée Par La FAO,Résultat d'un Accord Entre La FAO et laCommission Des Communautés Euro-Péennes (Projet GCP/INT/422/EEC)Financée Conjointement Par Ces Deuxorganisations.Rome, FAO.*, volume 2.

[17] Fives, J. M. (1980). Littoral and Benthic Investigations on the West Coast of Ireland: XI. The Biology of Montagu's Blenny, Coryphoblennius galerita L. (Pisces), on the Connemara Coast. *Proceedings of the Royal Irish Academy. Section B: Biological, Geological, and Chemical Science*, 80B:61–77.

[18] Froese, R., Pauly, D., and Editors (2000). FishBase 2000: Concepts, design and data sources. page 344.

[19] García-Díaz, M. M., Tuset, V. M., González, J. A., and Socorro, J. (1997). Sex and reproductive aspects in Serranus cabrilla (Osteichthyes: Serranidae): Macroscopic and histological approaches. *Marine Biology*, 127(3):379–386.

[20] Gonçalves, J. and Erzini, K. (2000). The reproductive biology of Spondyliosoma cantharus (L.) from the SW Coast of Portugal. *Scientia Marina*, 64:403–411.

[21] Gordoa, A., Molí, B., and Raventós, N. (2000). Growth performance of four wrasse species on the north-western Mediterranean coast. *Fisheries Research*, 45(1):43–50.

[22] Kara, M. H. (1997). Cycle sexuel et fécondité du Loup Dicentrarchus labrax (Poisson Moronidé) du golfe d'Annaba. *Cahiers de Biologie Marine*, (3).

[23] Kara, M. H. and Quignard, J.-P. (2018a). *Les poissons des lagunes et des estuaires de Méditerranée. 2, 2,.*

[24] Kara, M. H. and Quignard, J.-P. (2018b). *Les poissons des lagunes et des estuaires de Méditerranée 3B: Les poissons migrateurs.*

[25] Klimogiann, A., Kalanji, M., Pyrenis, G., Zoulioti, A., and Trakos, G. (2011). Ontogeny of Embryonic and Yolk-Sac Larval Stage of the Sparid Sharpsnout Sea Bream (Diplodus puntazzo Cetti, 1777). *Journal of Fisheries and Aquatic Science*, 6:62–73.

[26] Klimogianni, A., Koumoundouros, G., Kaspiris, P., and Kentouri, M. (2004). Effect of temperature on the egg and yolk-sac larval development of common pandora, Pagellus erythrinus. *Marine Biology*, 145(5):1015–1022.

[27] Kraljević, M., Matić-Skoko, S., Dul\vcić, J., Pallaoro, A., Jardas, I., and Glamuzina, B. (2007). Age and growth of sharpsnout seabream Diplodus puntazzo (Cetti, 1777) in the eastern Adriatic Sea.

[28] Landa, J., Pereda, P., Duarte, R., and Azevedo, M. (2001). Growth of anglerfish (Lophius piscatorius and L. budegassa) in Atlantic Iberian waters. *Fisheries Research*, 51(2):363–376.

[29] Lejeune, P. (1987). The effect of local stock density on social behavior and sex change in the Mediterranean labrid Coris julis. *Environmental Biology of Fishes*, 18(2):135–141.

[30] Lourie, S. A. (2004). A guide to the identification of seahorses.

[31] Macías, D., Gomez Vives, M. J., García, S., and Urbina, J. (2005). Reproductive characteristics of Atlantic bonito (Sarda sarda) from the south-western Spanish Mediterranean. 58.

[32] Milton, P. (1983). Biology of littoral blenniid fishes on the coast of south-west England. *Journal of the Marine Biological Association of the United Kingdom*, 63(1):223–237.

[33] Mouine, N., Ktari, M.-H., and Chakroun-Marzouk, N. (2011). Reproductive characteristics of Spondyliosoma cantharus (Linnaeus, 1758) in the Gulf of Tunis. *Journal of Applied Ichthyology*, 27(3):827–831.

[34] Mouine-Oueslati, N., Romdhani, A., Chater, I., Ktari, M.-H., and Chakroun-Marzouk, N. (2015). Age and growth of Spondyliosoma cantharus (Sparidae) in the Gulf of Tunis. *Scientia Marina*, 79:319–324.

[35] Murua, H. (2010). The Biology and Fisheries of European Hake, Merluccius merluccius, in the North-East Atlantic. In *Advances in Marine Biology*, volume 58, pages 97–154. Elsevier.

[36] Murua, H., Motos, L., and LUCIO, P. (1998). Reproductive modality and batch fecundity of the European hake (Merluccius merluccius L.) in the Bay of Biscay. *California Cooperative Oceanic Fisheries Investigations Reports*, 39.

[37] Orsi Relini, L., Garibaldi, F., Cima, C., Palandri, G., Lanteri, L., and Relini, M. (2005). Biology of atlantic bonito, Sarda sarda (Bloch, 1793), in the western and central mediterranean. A summary concerning a possible stock unit.

[38] Ortega Garcia, A. (2015). *Cultivo Integral de Dos Especies de Escómbridos : Atún Rojo Del Atlántico (Thunnus Thynnus, L. 1758) y Bonito Atlántico (Sarda Sarda, Bloch 1793).* PhD thesis, Universidad de Murcia.

[39] Palacios Sartagal, N. (2017). *Estudi de La Fecunditat i Estratègia Reproductiva de Serranus Cabrilla (Pisces, Serranidae).* PhD thesis, Universitat de Girona.

[40] Papaconstantinou, C., G.Petrakis, and Vassilopoulou, V. (1986). The fecundity of hake (Merluccius merluccius L.) and red pandora (Pagellus erythrinus L.) in the Greek seas. *Acta Adriatica*, 27:85–95.

[41] Ré, P. and Meneses, I. (2008). *Early stages of marine fishes occurring in the Iberian peninsula.* IPIMAR, Lisboa.

[42] Rispoli, V. F. and Wilson, A. B. (2008). Sexual size dimorphism predicts the frequency of multiple mating in the sex-role reversed pipefish Syngnathus typhle. *Journal of Evolutionary Biology*, 21(1):30–38.

[43] Rodriguez, J. M. (2017). *A Guide to the Eggs and Larvae of 100 Common Western Mediterranean Sea Bony Fish Species.* Number Non Serial (FishFinder Guide eggs&larvae). FAO, Rome, Italy.

[44] Škeljo, F. (2012). *Dinamika Populacije Kneza, Coris Julis (Linnaeus, 1758) u Istočnom Jadranu.* PhD thesis, University of Split.

[45] Svensson, I. (1988). Reproductive Costs in Two Sex-Role Reversed Pipefish Species (Syngnathidae). *Journal of Animal Ecology*, 57(3):929–942.

[46] Taieb, A. H., Ghorbel, M., and Jarboui, O. (2013). Study of fecundity for Diplodus vulgaris (Teleost, Sparidae) in Gulf of Gabes. page 5.

[47] Tsikliras, A. C. and Stergiou, K. I. (2015). Age at maturity of Mediterranean marine fishes. *Mediterranean Marine Science*, 16(1):5–20.

[48] Uçkun İlhan, D., Akalın, S., tosunoğlu, Z., and Ozaydin, O. (2010). Growth Characteristics and Reproduction of Comber, Serranus Cabrilla (Actinopterygii, Perciformes, Serranidae), in the Aegean Sea. *Acta Ichthyologica Et Piscatoria*, 40:55–60.

[49] Vigliola, L., Harmelin-Vivien, M. L., Biagi, F., Galzin, R., Garcia-Rubies, A., Harmelin, J.-G., Jouvenel, J.-Y., Direach-Boursier, L. L., Macpherson, E., and Tunesi, L. (1998). Spatial and temporal patterns of settlement among sparid fishes of the genus Diplodus in the northwestern Mediterranean. *Marine Ecology Progress Series*, 168:45–56.

### Additional figures

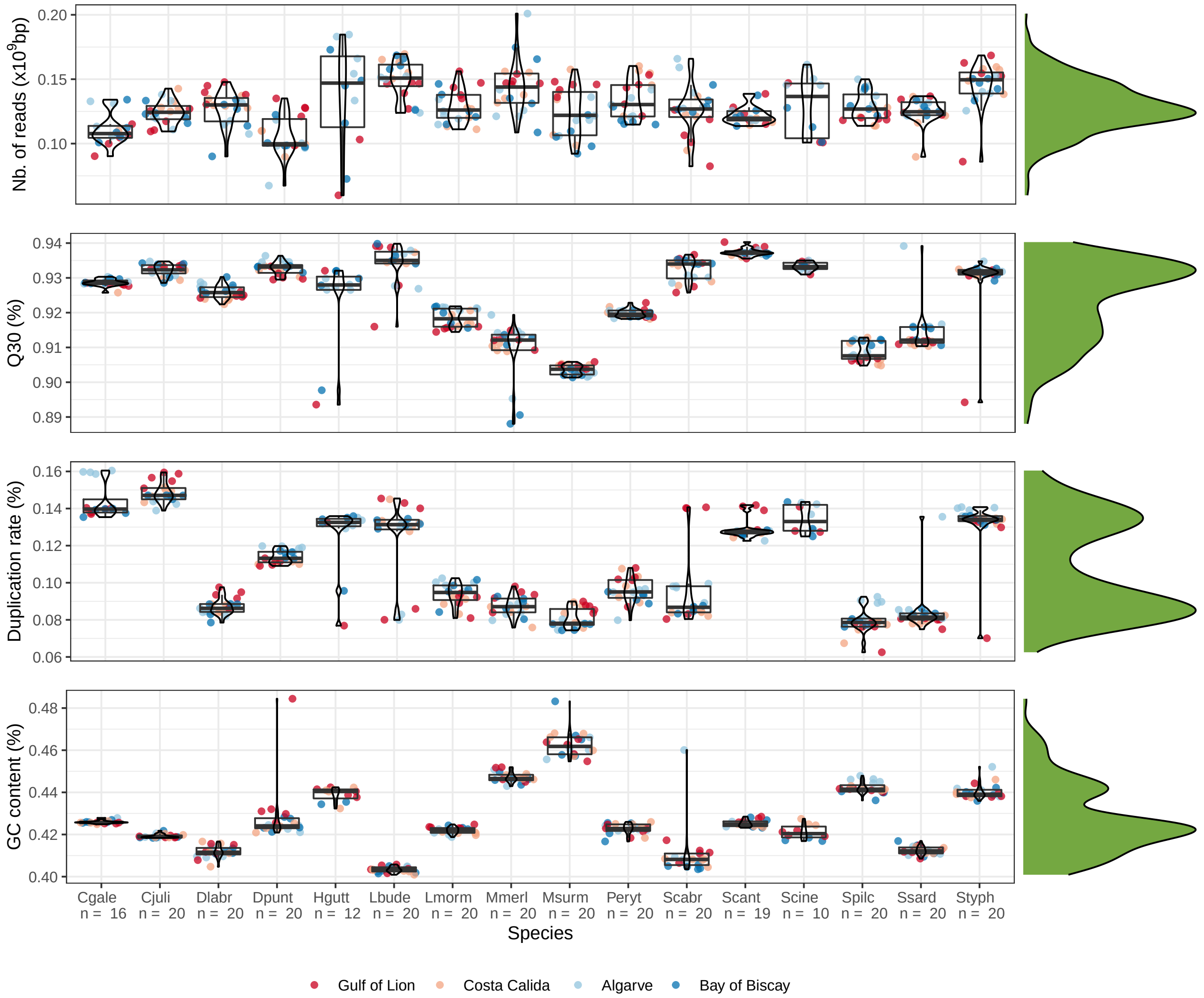

Figure S2: **Number of reads ($10^9$ bp), percentage of reads with quality superior to Q30, duplication rate and GC content after filtering, correcting and trimming steps carried out with `fastp` v.0.20.0 (Chen et al., 2018)**. Each point represents an individual unassembled genome: species are represented on the $y$-axis. Colors represent sampling locations: mediterranean locations in warm colors (Gulf of Lion in dark red, Murcia in pale red), atlantic locations in cold colors (Faro in pale blue, Bay of Biscay in dark blue). Overall distributions of each parameters are represented on the right side of each panel. Cgale = *Coryphoblennius galerita*, Cjuli = *Coris julis*, Dlabr = *Dicentrarchus labrax*, Dpunt = *Diplodus puntazzo*, Hgutt = *Hippocampus guttulatus*, Lbude = *Lophius budegassa*, Lmorm = *Lithognathus mormyrus*, Mmerl = *Merluccius merluccius*, Msurm = *Mullus surmuletus*, Peryt = *Pagellus erythrinus*, Scabr = *Serranus cabrilla*, Scant = *Spondyliosoma cantharus*, Scine = *Symphodus cinereus*, Spilc = *Sardina pilchardus*, Ssard = *Sarda sarda*, Styph = *Syngnathus typhle*.

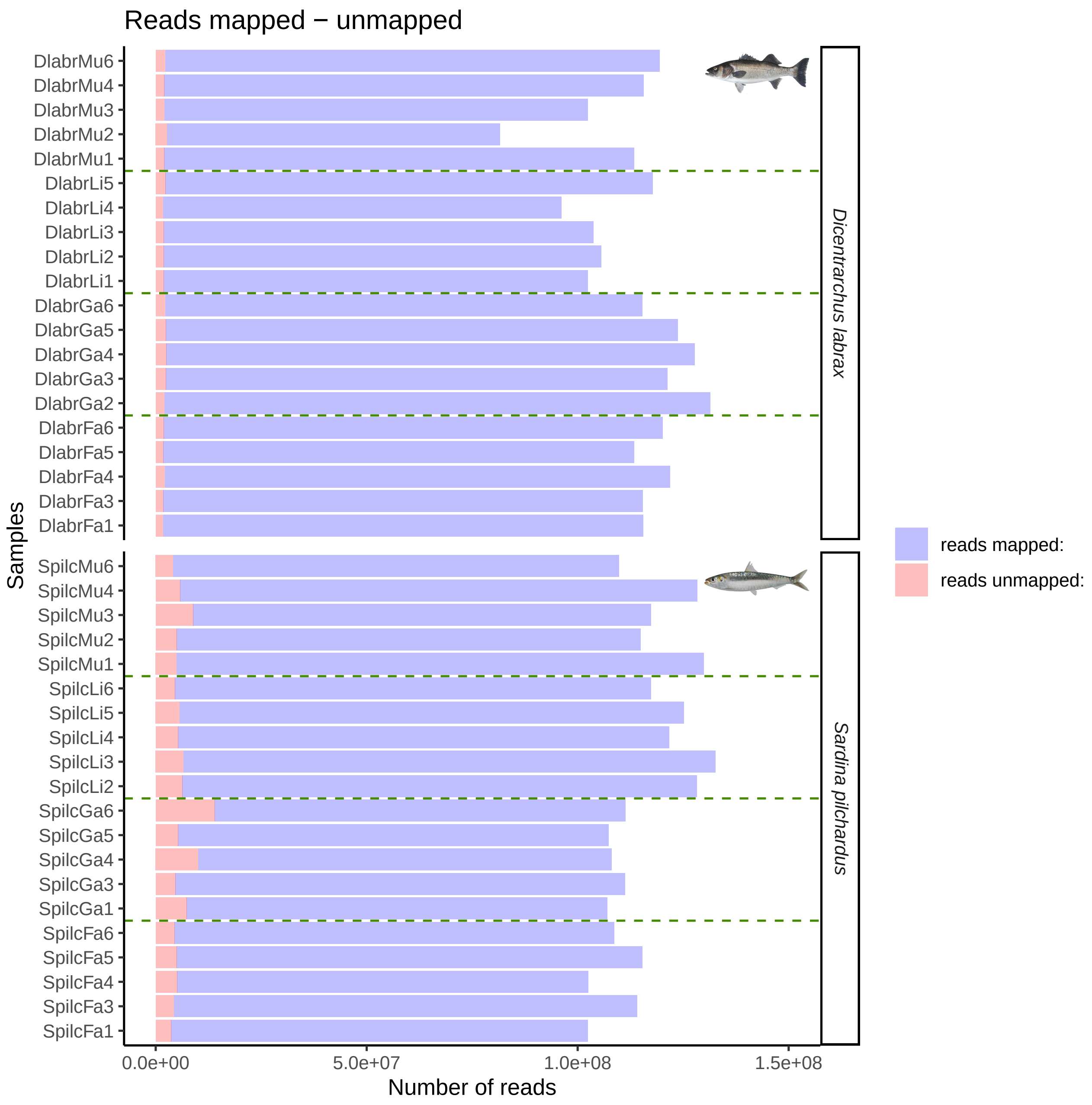

Figure S3: **Mapping statistics** - Number of reads mapped in red and unmapped blue with `bwa` v.0.7.17 for the 20 indiviuals of *D. labrax* and *S. pilchardus*.

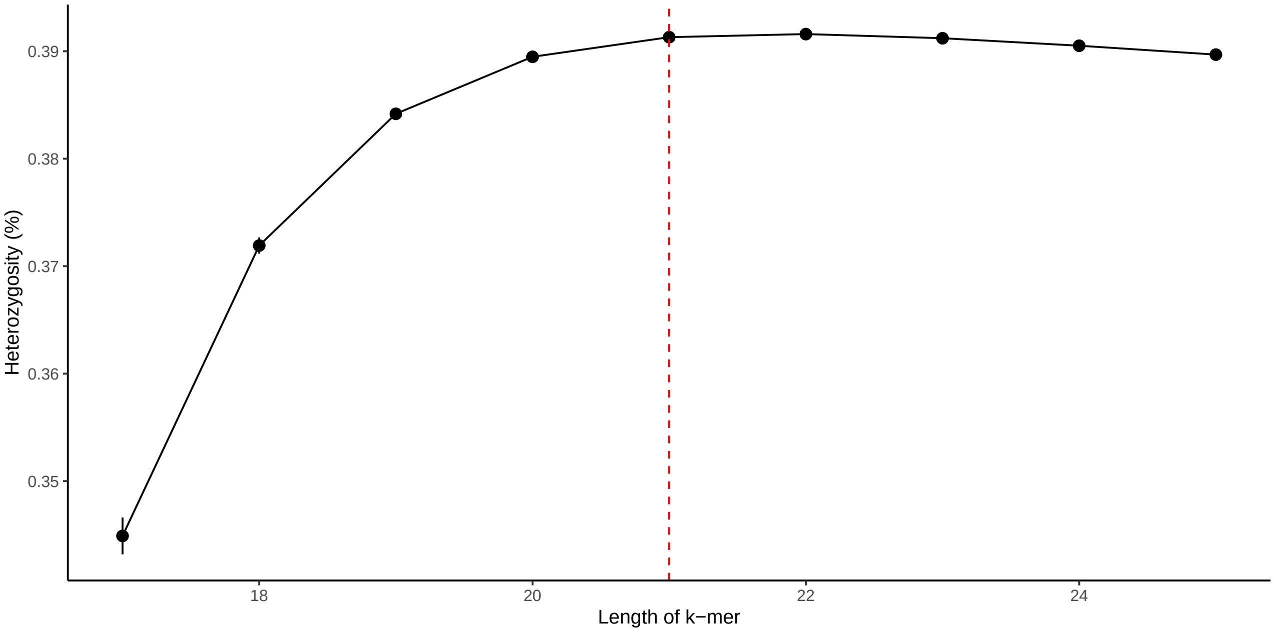

Figure S4: **Effect of $k - mer$ length on genetic diversity estimation with `GenomeScope` v.1.0 (Vurture et al., 2017)** - Genetic diversity was estimated with `GenomeScope` with different values of $k - mer$ length (17,19,21,23 and 25) on one European sea bass individual (*D. labrax*), all other parameters being equal (as detailed in the main text). The red vertical dashed line represents the value used in this study (21).

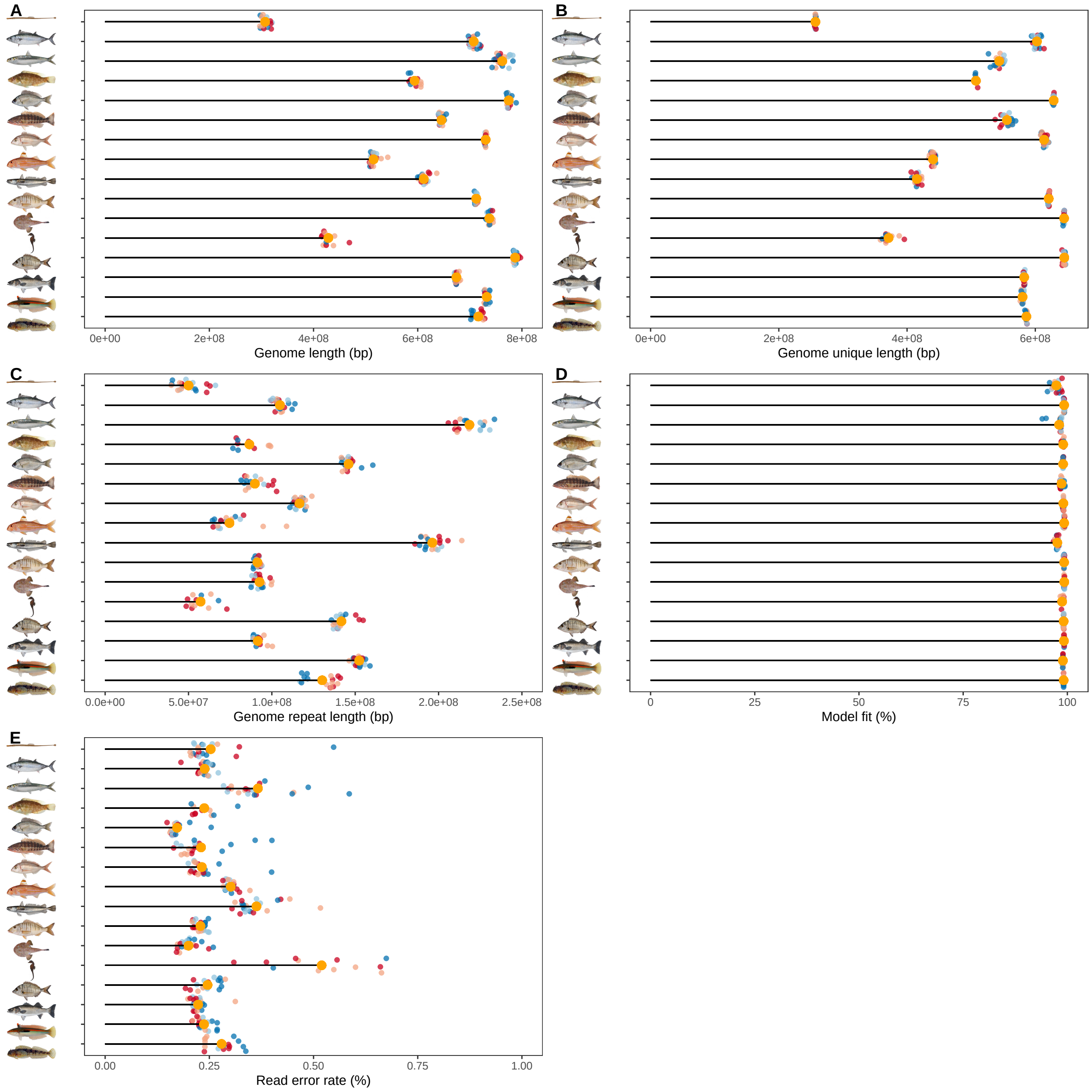

Figure S5: **Individual whole-genome sequences features estimated with `GenomeScope` v.1.0 (Vurture et al., 2017)**. A) Genome length (number of base pairs), B) genome unique length (number of base pairs), C) genome repeat length (number of base pairs), D) read error rate (%) and E) model fit (%). Individual feature and species median feature is represented in yellow in each panel. Species illustrations were retrieved from Iglésias (2013) with permissions.

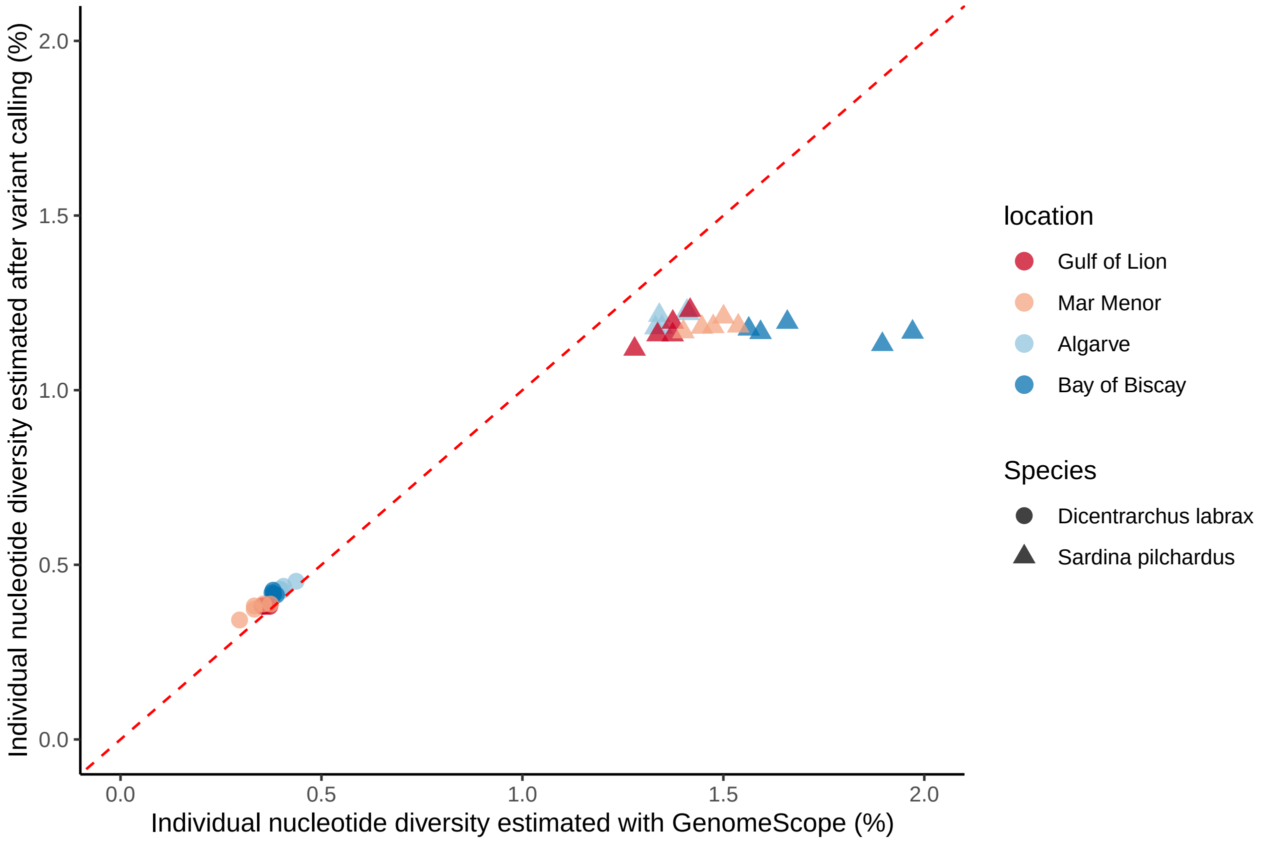

Figure S6: Relationship between individual mean genome-wide heterozygosity estimated with the k-mer based reference-free approach in GenomeScope ($x$-axis), and the high standard reference-based approach in GATK ($y$-axis), for european sea bass (*D. labrax*, dots) and european pilchard (*S. pilchardus*, triangle).

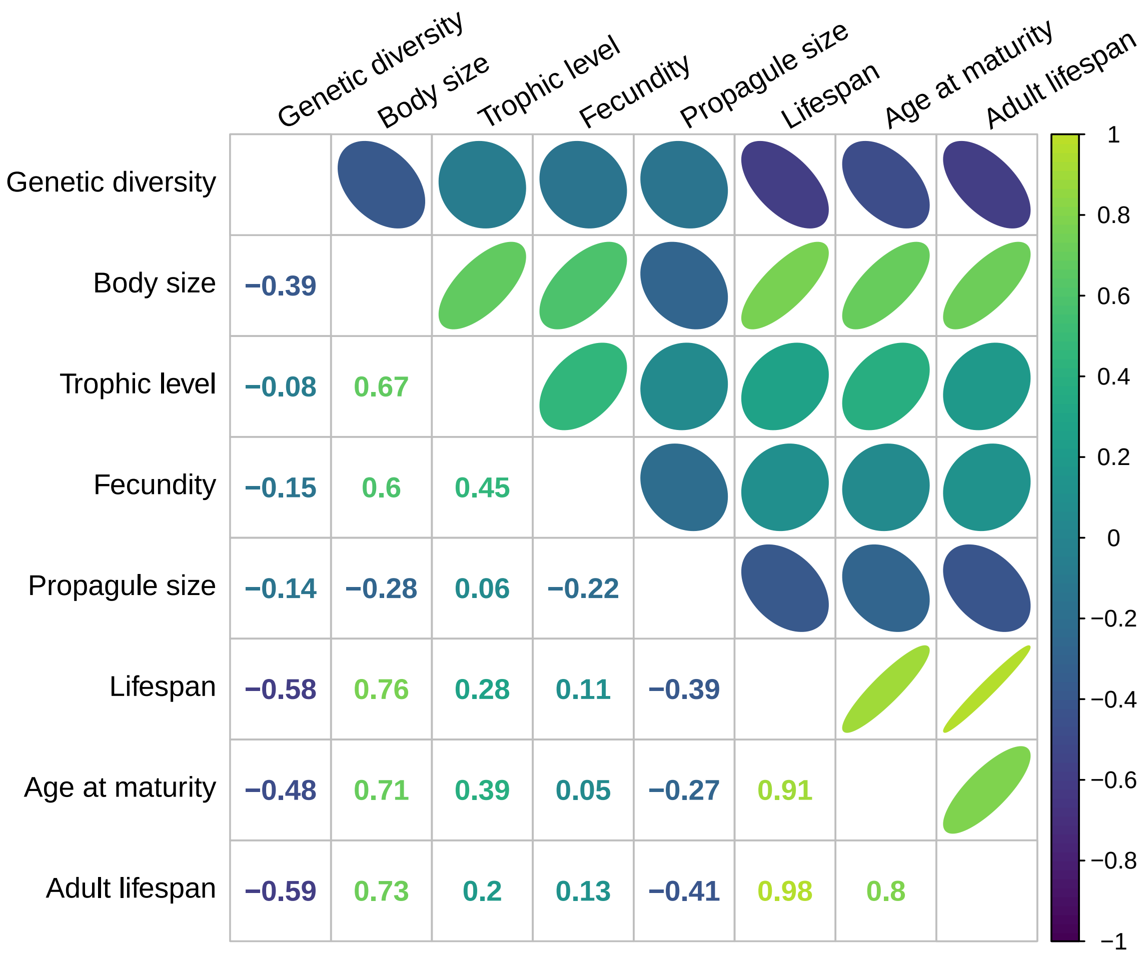

Figure S7: **Correlation matrix between genetic diversity and all quantitative life history traits** - Upper triangle represents the strength and the direction of the correlation, the lower triangle the coefficient of correlation.

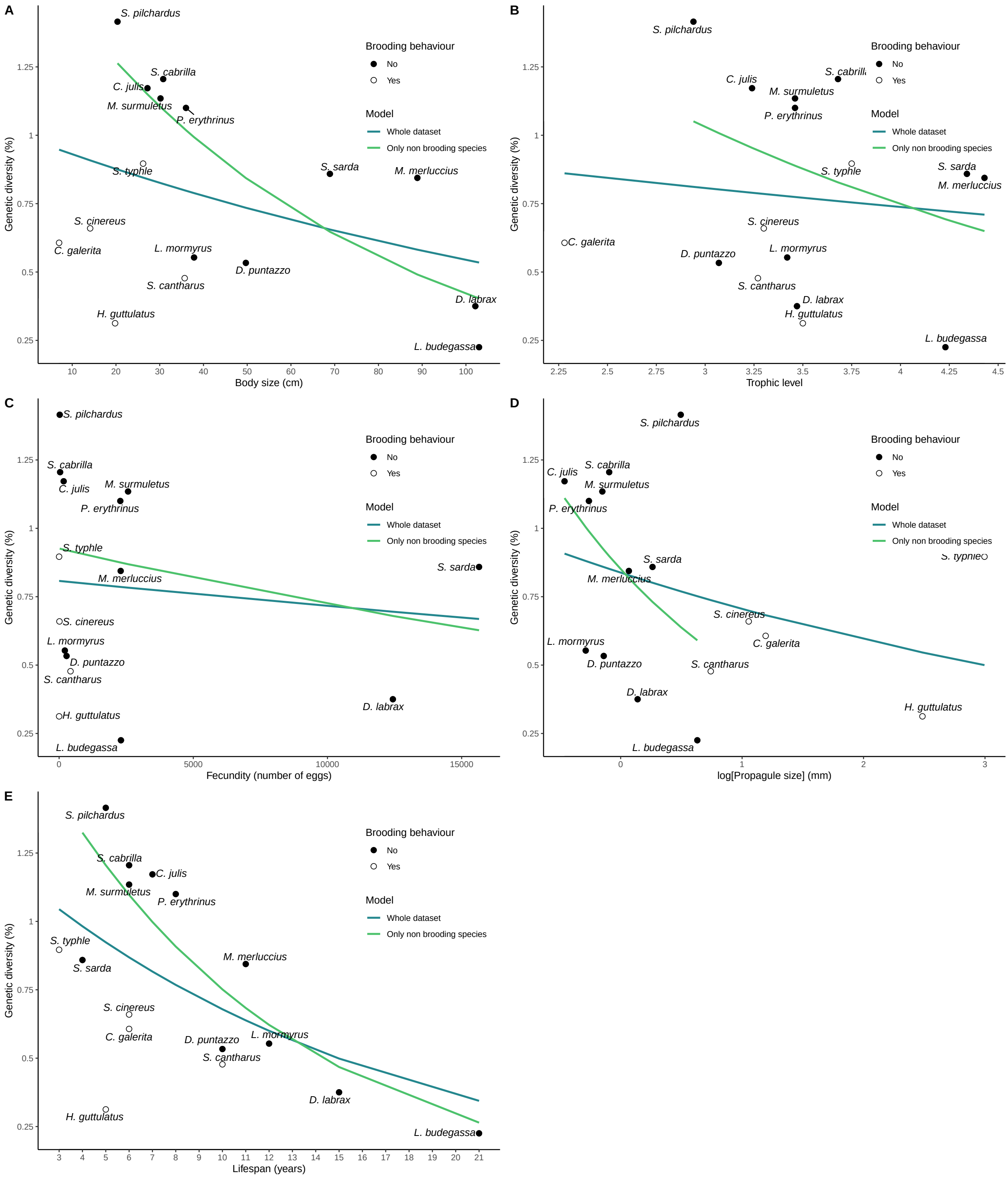

Figure S8: **Relationship between species median genetic diversity (%) and 5 covariables** - Each point represents the median of observed genetic diversity among individuals within each species. Full points represents non-brooding species, empty circles, brooding species. Blue and green line represent the beta regression between each predictive variable and genetic diversity considering either the whole dataset (16 species), or the 11 non-brooding species only, respectively. **A)** adult body size, **B)** trophic level, **C)** fecundity, **D)**, propagule size and **E)**, lifespan.

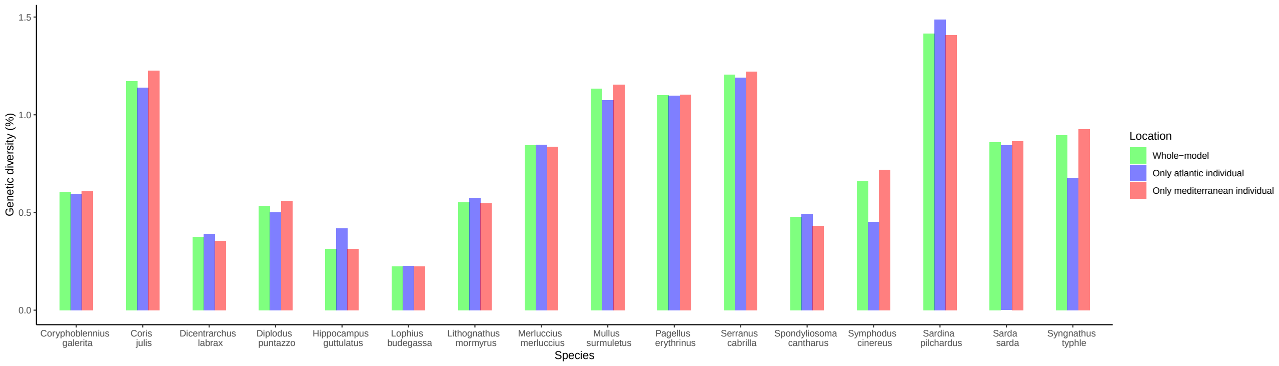

Figure S9: **Effect of population structure on genetic diversity estimates**. For each species on the $x$-axis, genetic diversity is estimate from the median of individual genome-wide heterozygosities for all the individuals (in green), from the individuals from the Mediterranean Sea (in red) or from the individuals from the Atlantic Ocean (in blue).

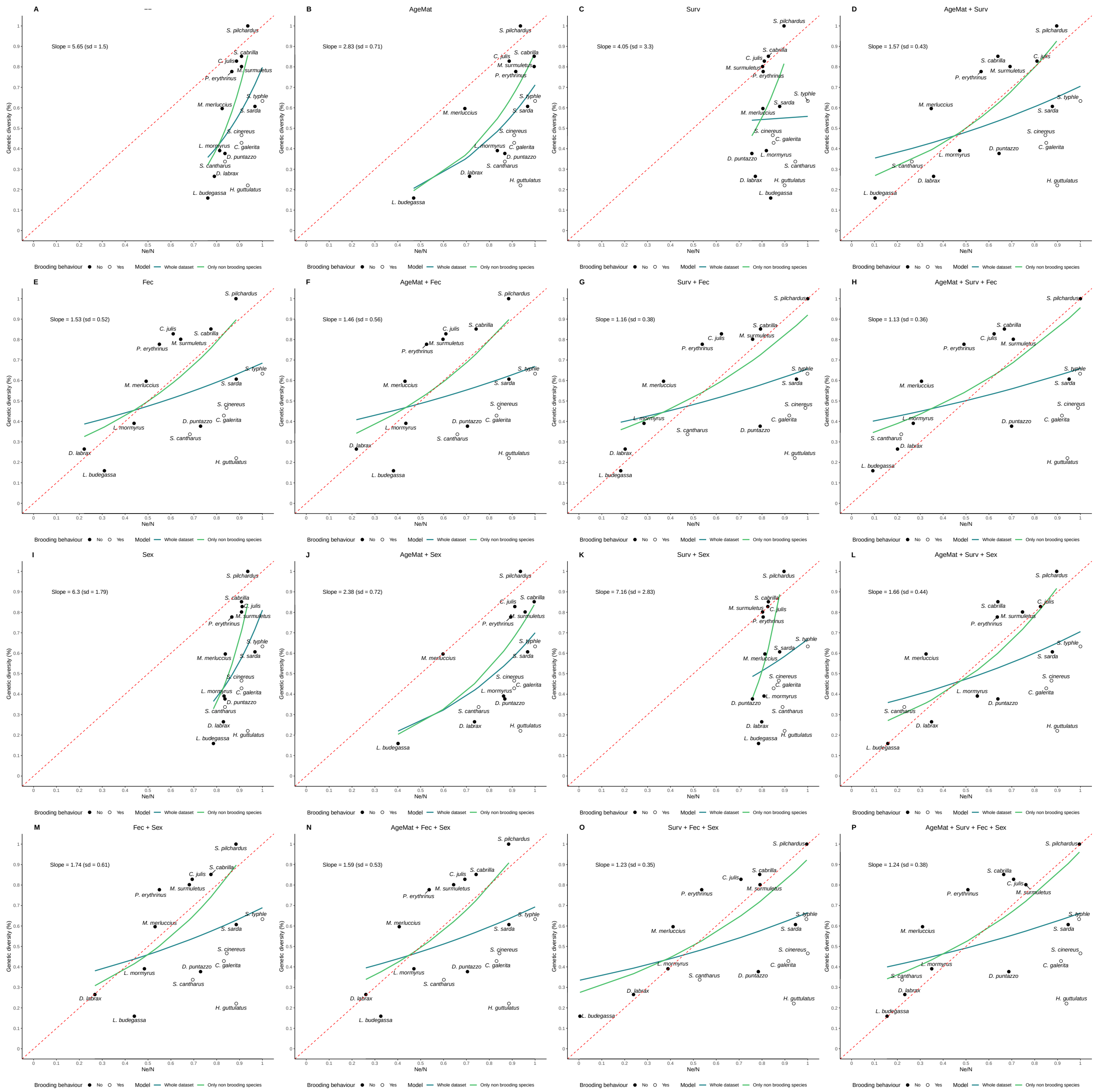

Figure S10: **Relationship between relative species genetic diversity and relative $\frac{N_e}{N}$ estimated by AgeNe for 16 sets of life tables.** Estimated species genetic diversity divided by the highest empirical estimates (i.e. *Sardina pilchardus*) on $y$-axis, and $\frac{N_e}{N}$ estimated by AgeNe divided by the highest estimates in each of the 16 sets of life tables. Each of the 16 panels represents the impact of species life tables considering some characteristics: AgeMat (delayed first age at maturity), Surv (age-specific survival estimated from empirical species age-length distributions), Fec (empirical estimates of increasing fecundity with age), Sex (sex-specific differences in either, age at first maturity, age-specific survival or/and fecundity). A) Age at first maturity at 1 year old; constant age-specific survival; constant age-specific fecundity and no differences between sex life tables, B) Species-specific age at first maturity; constant age-specific survival; constant age-specific fecundity and no differences between sex life tables, C) Age at first maturity at 1 year old; increasing age-specific survival rate; D) Species-specific age at first maturity; increasing age-specific survival rate; constant age-specific fecundity and no differences between sex life tables, constant age-specific fecundity and no differences between sex life tables, E) Age at first maturity at 1 year old; constant age-specific survival; increasing fecundity with age and no differences between sex life tables, F) Species-specific age at first maturity; constant age-specific survival; increasing fecundity with age and no differences between sex life tables, G) Age at first maturity at 1 year old; increasing age-specific survival rate; increasing fecundity with age and no differences between sex life tables, H) Species-specific age at first maturity; increasing age-specific survival rate; increasing fecundity with age and no differences between sex life tables, I) Age at first maturity at 1 year old; constant age-specific survival; constant age-specific fecundity and sex-specific differences between life tables, J) Species-specific age at first maturity; constant age-specific survival; constant age-specific fecundity and sex-specific differences between life tables, K) Age at first maturity at 1 year old; increasing age-specific survival rate; constant age-specific fecundity and sex-specific differences between life tables, L) Species-specific age at first maturity; increasing age-specific survival rate; constant age-specific fecundity and sex-specific differences between life tables, M) Age at first maturity at 1 year old; constant age-specific survival; increasing fecundity with age and sex-specific differences between life tables, N) Species-specific age at first maturity; constant age-specific survival; increasing fecundity with age and sex-specific differences between life tables, O) Age at first maturity at 1 year old; increasing age-specific survival rate; increasing fecundity with age and sex-specific differences between life tables, P) Species-specific age at first maturity; increasing age-specific survival rate; increasing fecundity with age and sex-specific differences between life tables. Full points represents non-brooding species, empty points, brooding species. Dashed blue line and solid green represent the beta regression between relative genetic diversity and relative estimated $\frac{N_e}{N}$. Dotted red line represents the identity curve, with slope equals 1 and intercept 0

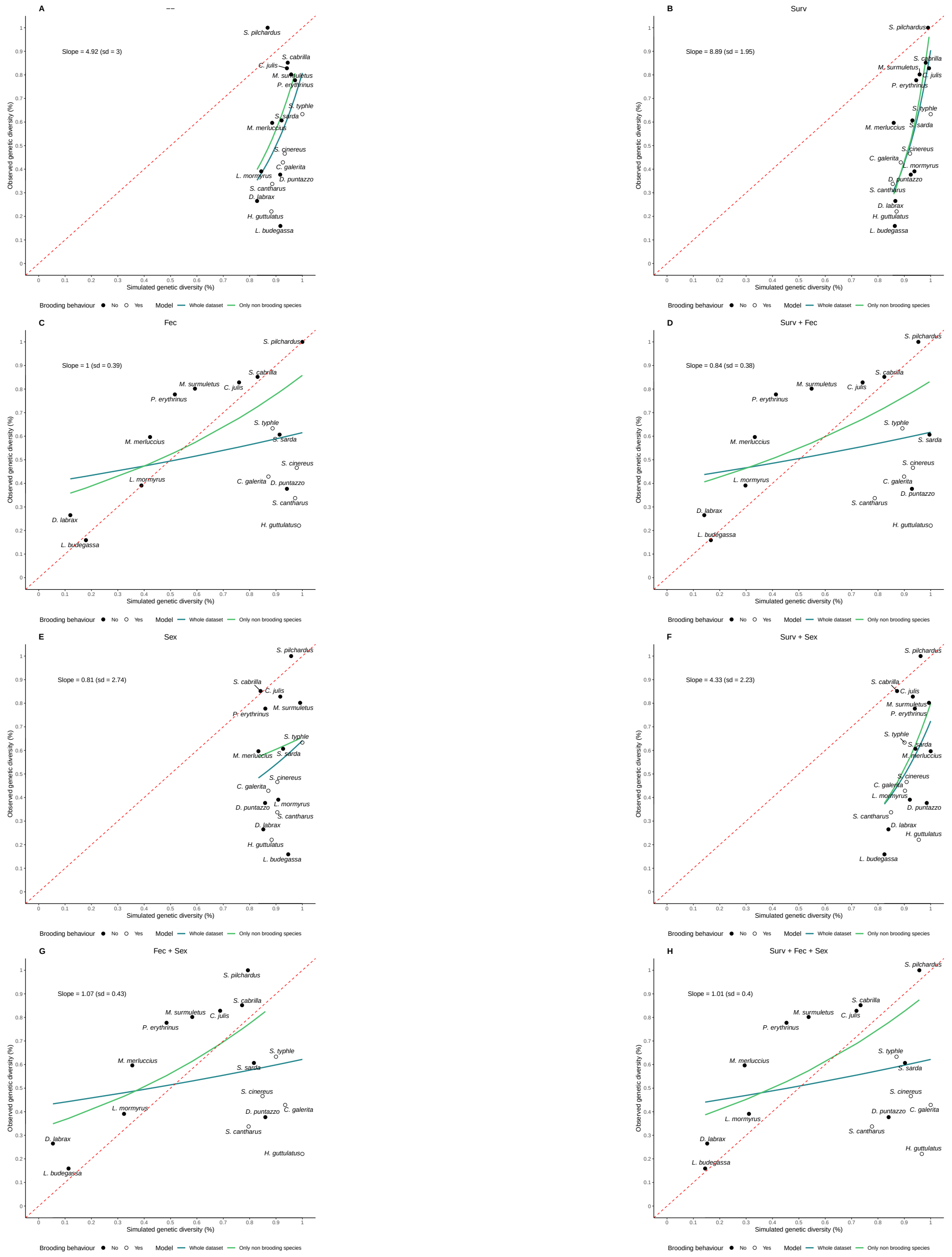

Figure S11: **Relationship between relative species genetic diversity and simulated genetic diversity with forward-in-time simulations for 16 sets of life tables**. Estimated species genetic diversity divided by the highest empirical estimates (i.e. *Sardina pilchardus*) on *y*-axis, and simulated genetic diversity estimated by `SLiM` v.3.3.1 (Haller and Messer, 2017) divided by the highest estimates in each of the 16 set of life tables. Each of the 16 panels represents species life tables considering some characteristics: AgeMat (delayed first age of maturity), Surv (age-specific survival estimated from empirical species age-length distributions), Fec (empirical estimates of increasing fecundity with age), Sex (sex-specific differences in either, age at first maturity, age-specific survival or/and fecundity). A) Age at first maturity at 1 year old; constant age-specific survival; constant age-specific fecundity and no differences between sex life tables, B) Age at first maturity at 1 year old; increasing age-specific survival rate; C) Age at first maturity at 1 year old; constant age-specific survival; increasing fecundity with age and no differences between sex life tables, D) Age at first maturity at 1 year old; increasing age-specific survival ratel; increasing fecundity with age and no differences between sex life tables, E) Age at first maturity at 1 year old; constant age-specific survival; constant age-specific fecundity and sex-specific differences between life tables, F) Age at first maturity at 1 year old; increasing age-specific survival rate; constant age-specific fecundity and sex-specific differences between life tables, G) Age at first maturity at 1 year old; constant age-specific survival; increasing fecundity with age and sex-specific differences between life tables, H) Age at first maturity at 1 year old; increasing age-specific survival rate; increasing fecundity with age and sex-specific differences between life tables, Full points represents non-brooding species, empty points, brooding species. Dashed blue line and solid green represent the beta regression between relative genetic diversity and relative estimated $\frac{N_e}{N}$. Dotted red line represents the identity curve, with slope equals 1 and intercept 0

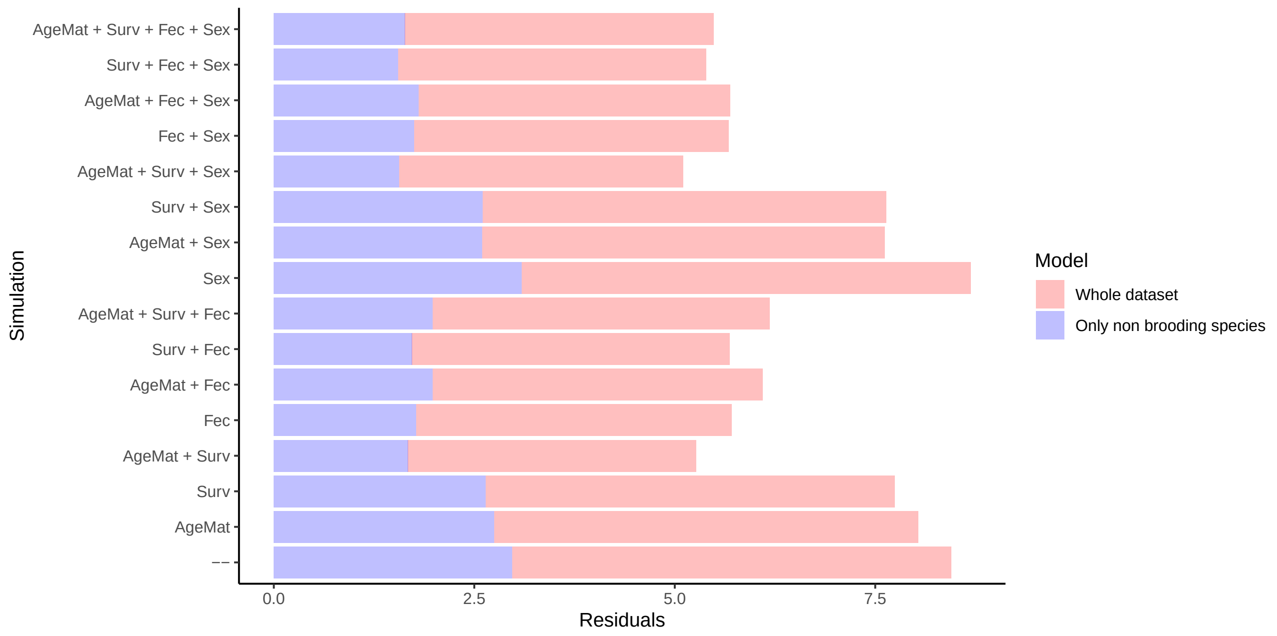

Figure S12: **Residuals of the linear model between genetic diversity and variance in reproductive success estimated from various combinations of life tables from a model with slope equals 1 and intercept 0**. For each set of life tables, we estimated $\frac{N_e}{N}$ with `AgeNe` and calculated the sum of squared deviation between these estimate and a model with slope equals 1 and intercept 0, as residuals. Higher residual number means variation in $\frac{N_e}{N}$ explain little the variation of observed genetic diversity.

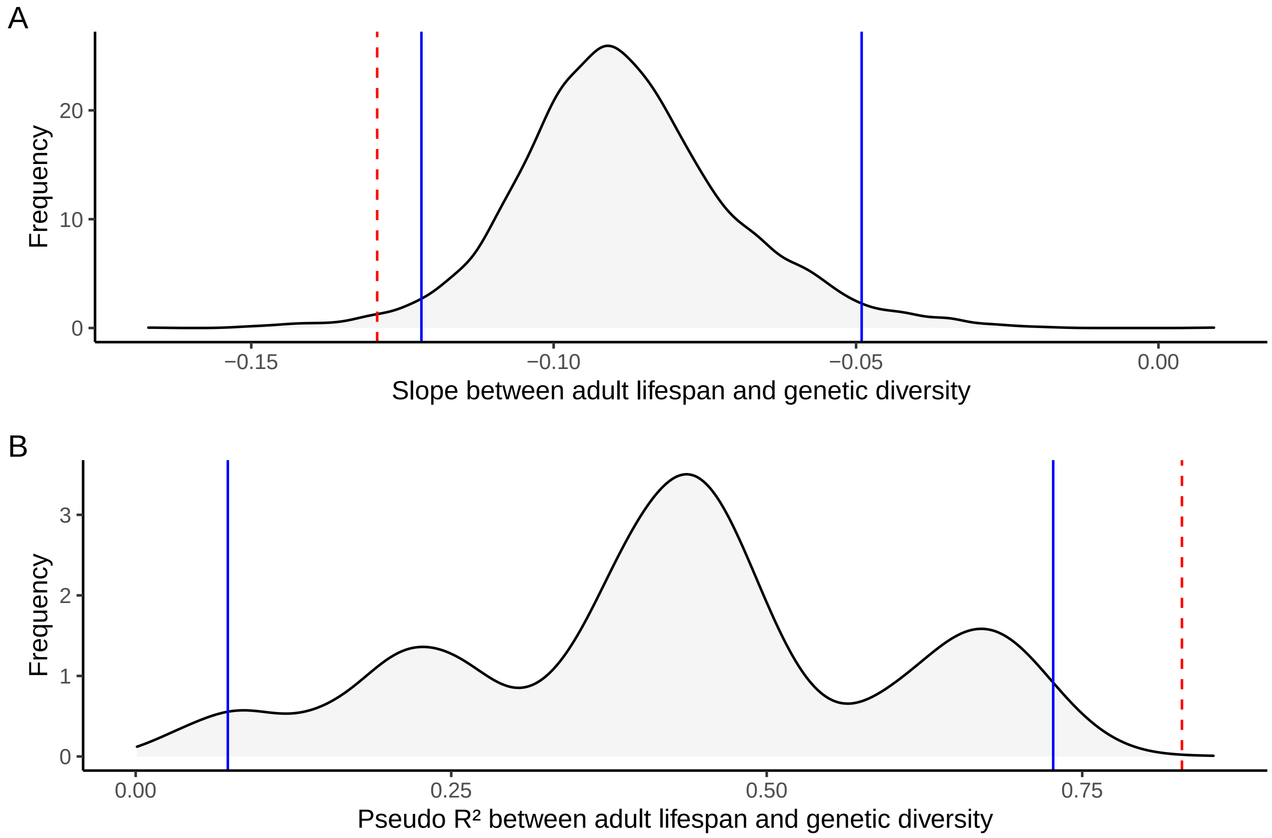

Figure S13: **Distribution of slope and pseudo $R^2$ of the beta regression between adult lifespan and genetic diversity for random subsets of 11 species** - We fit a beta regression between adult lifespan and genetic diversity for all sub-samples of 11 species and calculated the width of the 95% interval of the slope (A) and pseudo $R^2$ (B), represented with solid blue lines. Dashed red lines represent the estimated slope and pseudo $R^2$ for the sub-sample of the 11 species with no parental care behaviour. In (A) the 95% width interval ranged from -0.122 to -0.049, and the estimated slope for the non-brooders only was equal to -0.129. In (B), the 95% width interval ranged from 0.073 to 0.727, and the estimated pseudo $R^2$ for the non-brooders only was equal to 0.829.

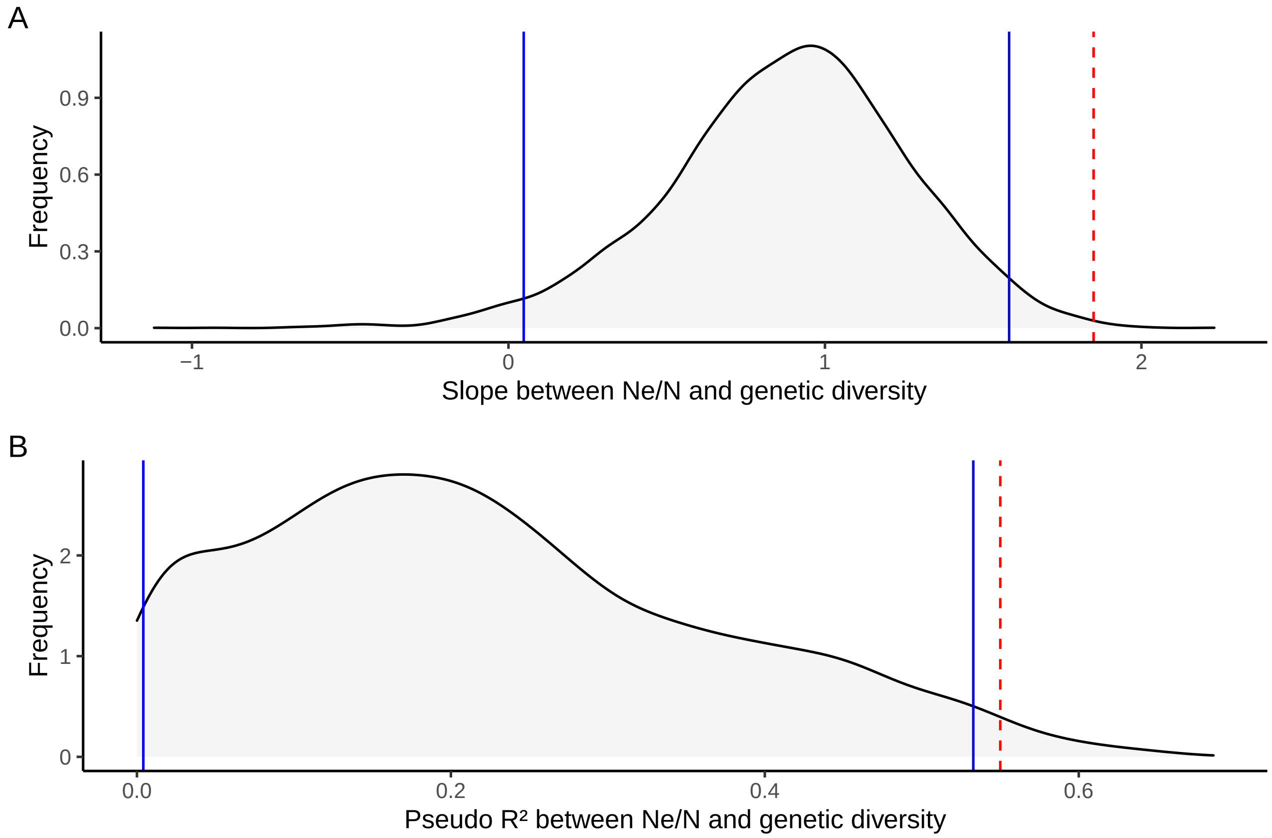

Figure S14: **Distribution of slope and pseudo $R^2$ of the beta regression between $\frac{N_e}{N}$ and genetic diversity for random subsets of 11 species** - We fit a beta regression between observed genetic diversity and $\frac{N_e}{N}$ for all sub-samples of 11 species and calculated the width of the 95% interval of the slope (A) and pseudo $R^2$ (B), represented with solid blue lines. Dashed red lines represent the estimated slope and pseudo $R^2$ for the sub-sample of the 11 species with no parental care behaviour. In (A) the 95% width interval ranged from 0.048 to 1.582, and the estimated slope for the non-brooders only was equal to 1.849. In (B), the 95% width interval ranged from 0.004 to 0.533, and the estimated pseudo $R^2$ from the non-brooders only was equal to 0.55.

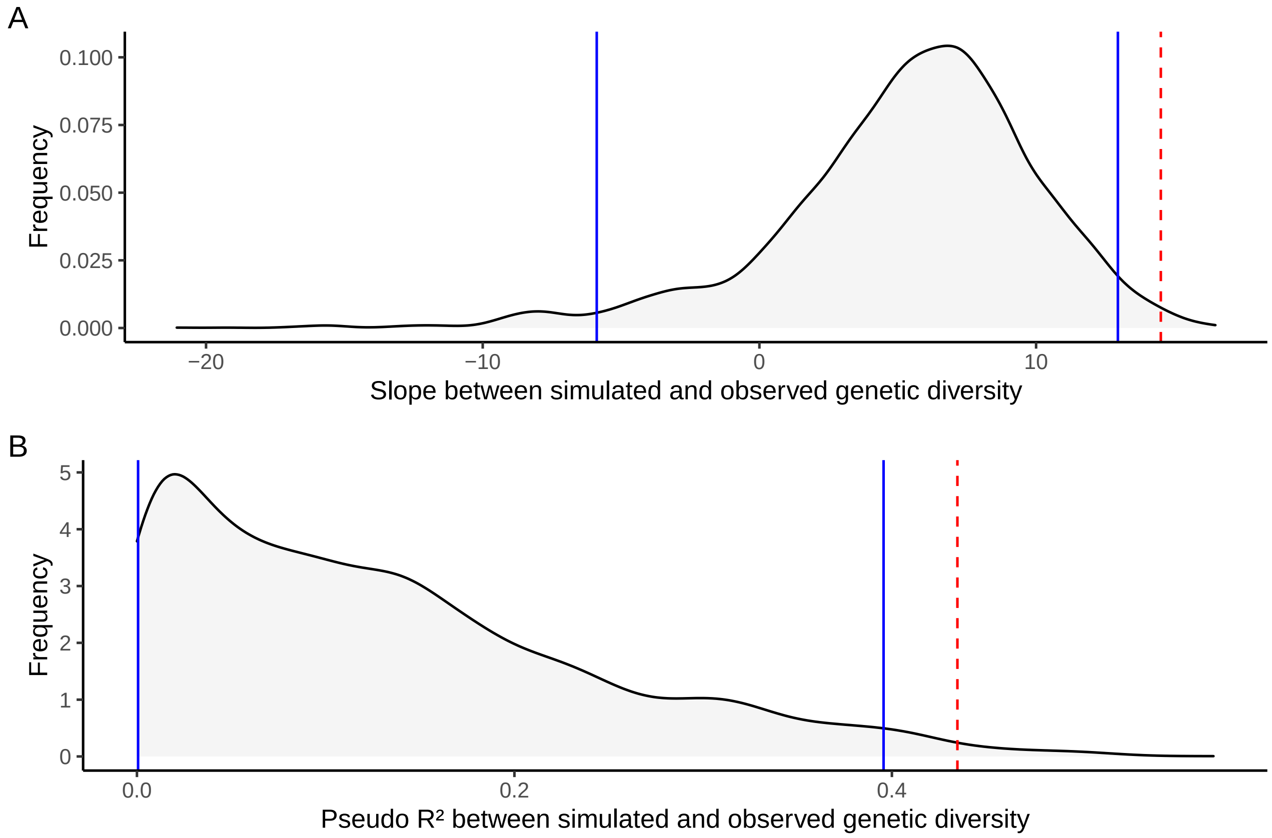

Figure S15: **Distribution of slope and pseudo $R^2$ of the beta regression between simulated, with `SLiM` v.3.3.1, and observed genetic diversity for random subset of 11 species** - We fit a beta regression between simulated genetic diversity and observed genetic diversity for all sub-samples of 11 species and calculated the width of the 95% interval of the slope (A) and pseudo $R^2$ (B), represented with solid blue lines. Dashed red lines represent the estimated slope and pseudo $R^2$ for the sub-sample of the 11 species with no parental care behaviour. In (A) the 95% width interval ranged from -5.879 to 12.957, and the estimated slope for the non-brooders only was equal to 14.509. In (B), the 95% width interval ranged from 0.001 to 0.396, and the estimated pseudo $R^2$ from the non-brooders only was equal to 0.435.

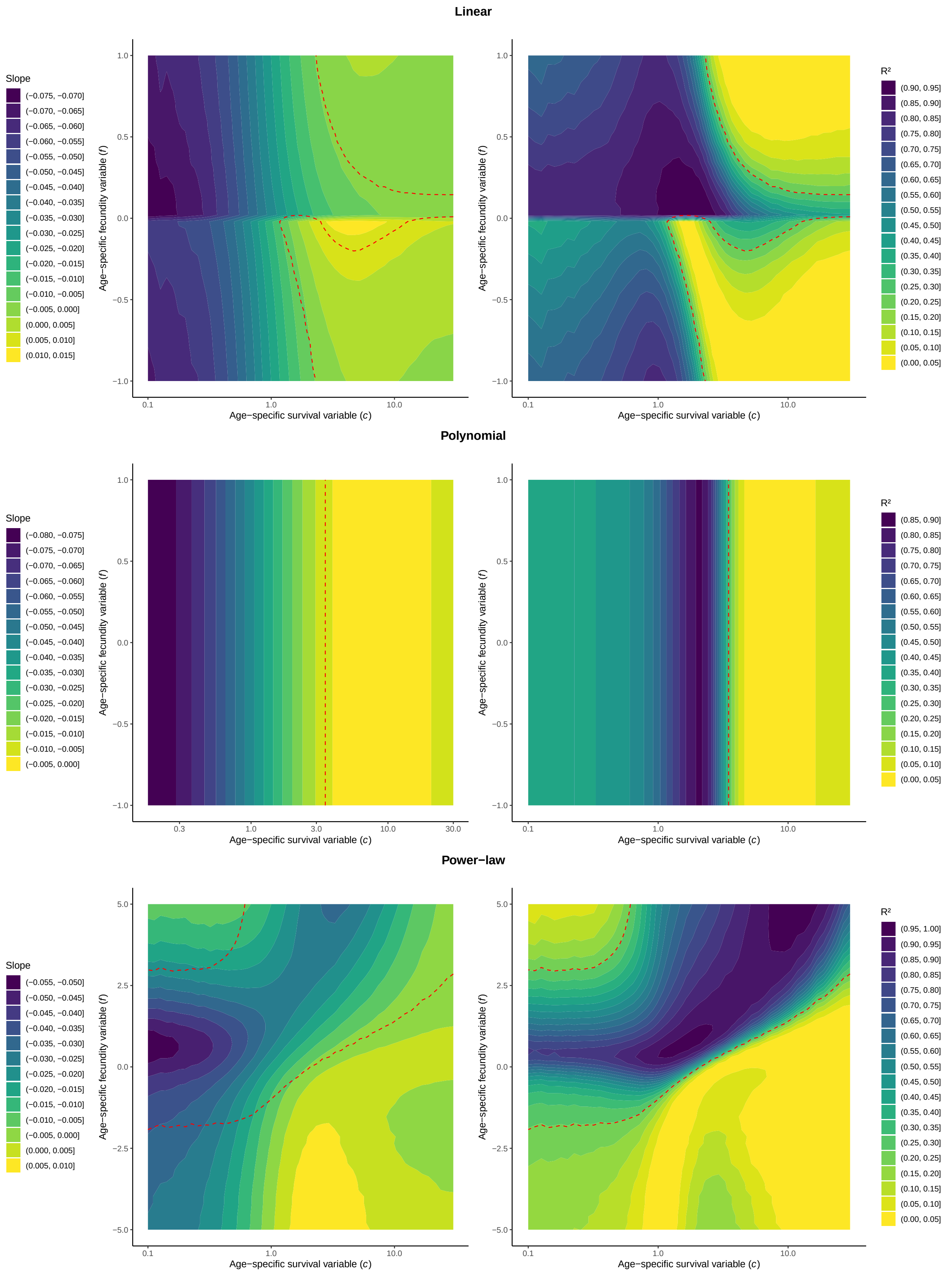

Figure S16: **Slope of and proportion of variance explained by linear models between adult lifespan and $\frac{N_e}{N}$ estimated with `AgeNe` for different combinations of age-specific survival and fecundity for three fecundity-age models: linear, polynomial and power-law**. Colder colors indicate steeper slope (left panels) or higher $R^2$ (right panels) of the regression between adult lifespan and $\frac{N_e}{N}$. On top, linear fecundity-age model ($F = \alpha \times L + \beta$), middle, polynomial fecundity-age model and on the bottom, power-law model, ($F = \alpha L^{\beta}$)

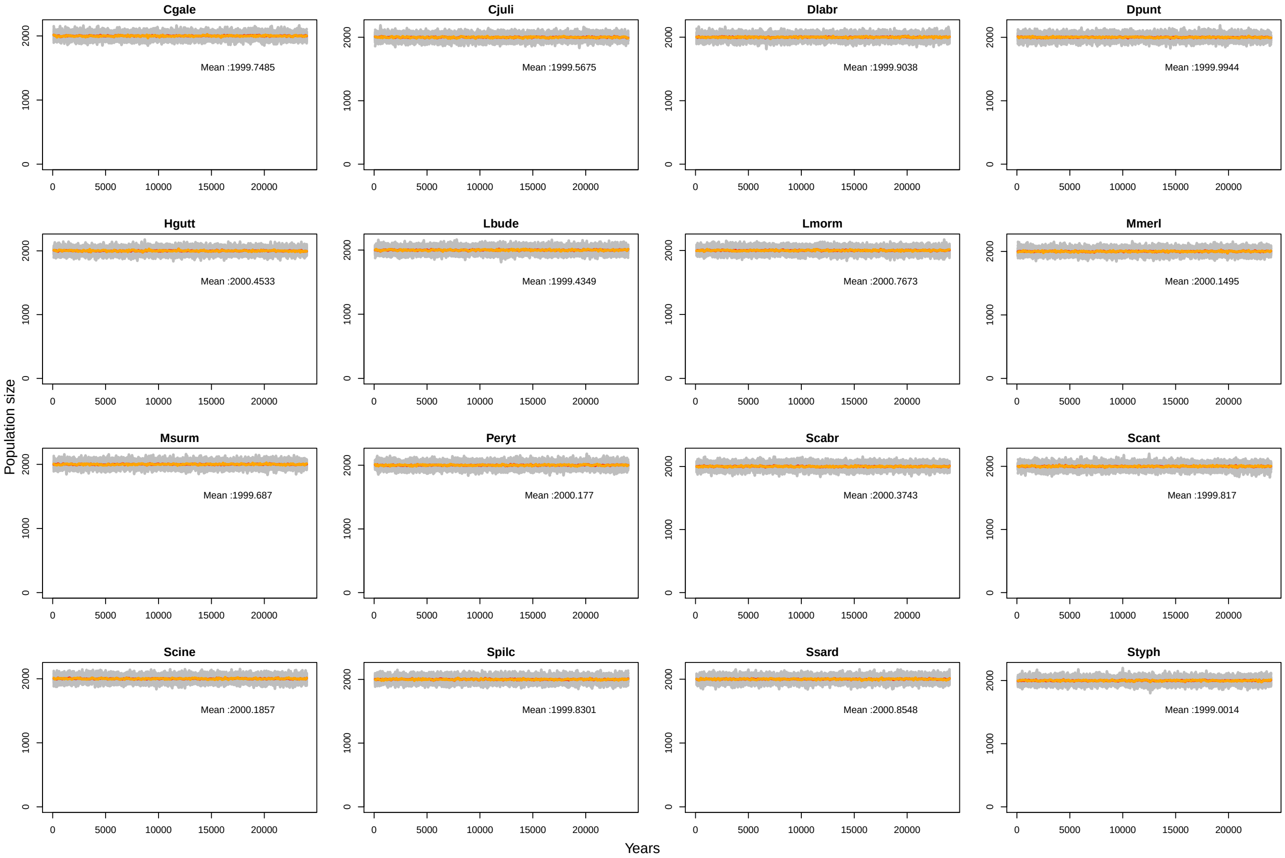

Figure S17: **Population size count for the 50 iterations of the 16 species for set 1 of life tables (age at first maturity at 1 year old, constant age-specific survival rate, constant age-specific fecundity and no differences between sex-specific life tables)**. Adult population size was count for each iteration every 100 years during the 25000 years of the simulation. Each grey line represents population size fluctuations for one iteration. For each species, orange and red line, respectively, represents the median and the mean for all the 50 iterations. Mean over all 50 iterations is written on the plot for each species. Cgale = *Coryphoblennius galerita*, Cjuli = *Coris julis*, Dlabr = *Dicentrarchus labrax*, Dpunt = *Diplodus puntazzo*, Hgutt = *Hippocampus guttulatus*, Lbude = *Lophius budegassa*, Lmorm = *Lithognathus mormyrus*, Mmerl = *Merluccius merluccius*, Msurm = *Mullus surmuletus*, Peryt = *Pagellus erythrinus*, Scabr = *Serranus cabrilla*, Scant = *Spondyliosoma cantharus*, Scine = *Symphodus cinereus*, Spilc = *Sardina pilchardus*, Ssard = *Sarda sarda*, Styph = *Syngnathus typhle*

.

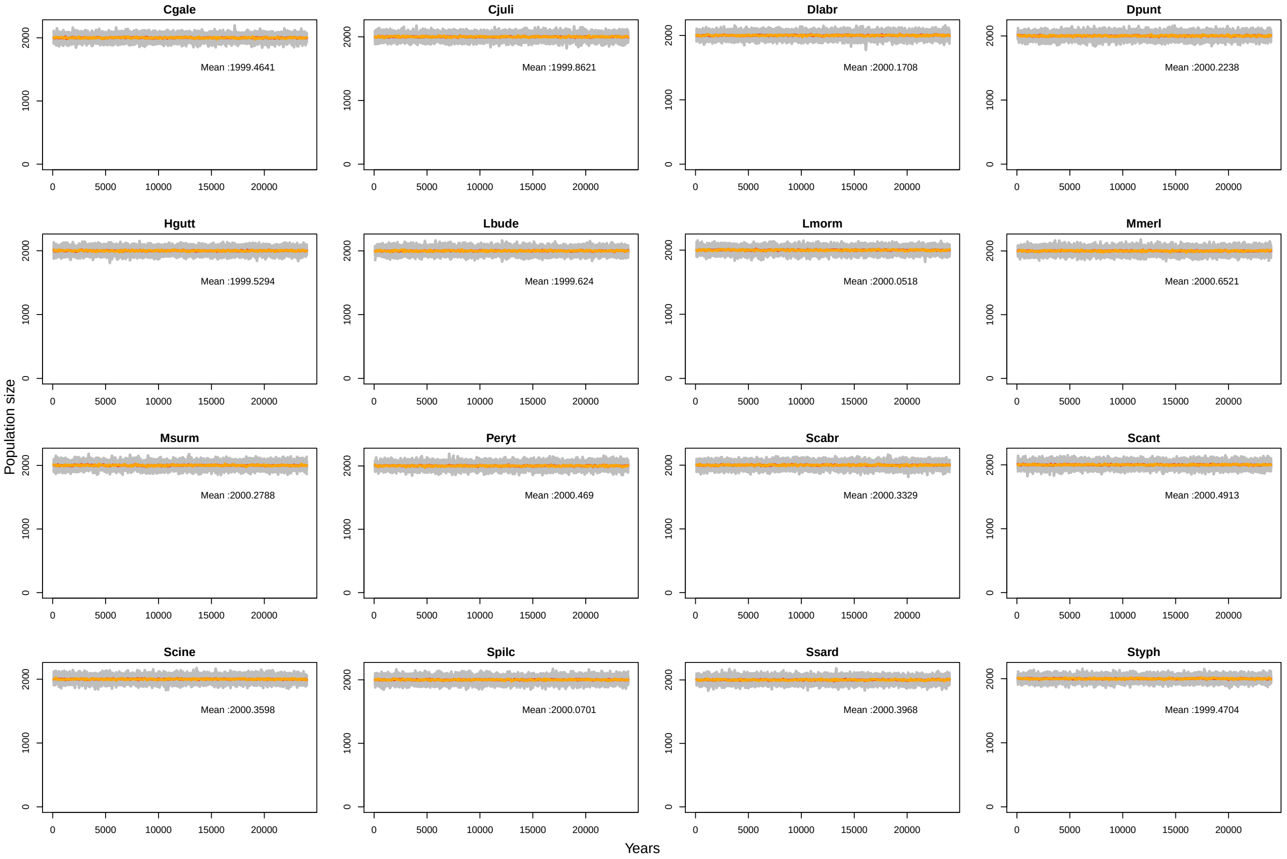

Figure S18: **Population size count for the 50 iterations of the 16 species for set 2 of life tables (age at first maturity at 1 year old, increasing age-specific survival rate, constant age-specific fecundity and no differences between sex-specific life tables)**. Adult population size was count for each iteration every 100 years during the 25000 years of the simulation. Each grey line represents population size fluctuations for one iteration. For each species, orange and red line, respectively, represents the median and the mean for all the 50 iterations. Mean over all 50 iterations is written on the plot for each species. Cgale = *Coryphoblennius galerita*, Cjuli = *Coris julis*, Dlabr = *Dicentrarchus labrax*, Dpunt = *Diplodus puntazzo*, Hgutt = *Hippocampus guttulatus*, Lbude = *Lophius budegassa*, Lmorm = *Lithognathus mormyrus*, Mmerl = *Merluccius merluccius*, Msurm = *Mullus surmuletus*, Peryt = *Pagellus erythrinus*, Scabr = *Serranus cabrilla*, Scant = *Spondyliosoma cantharus*, Scine = *Symphodus cinereus*, Spilc = *Sardina pilchardus*, Ssard = *Sarda sarda*, Styph = *Syngnathus typhle*

.

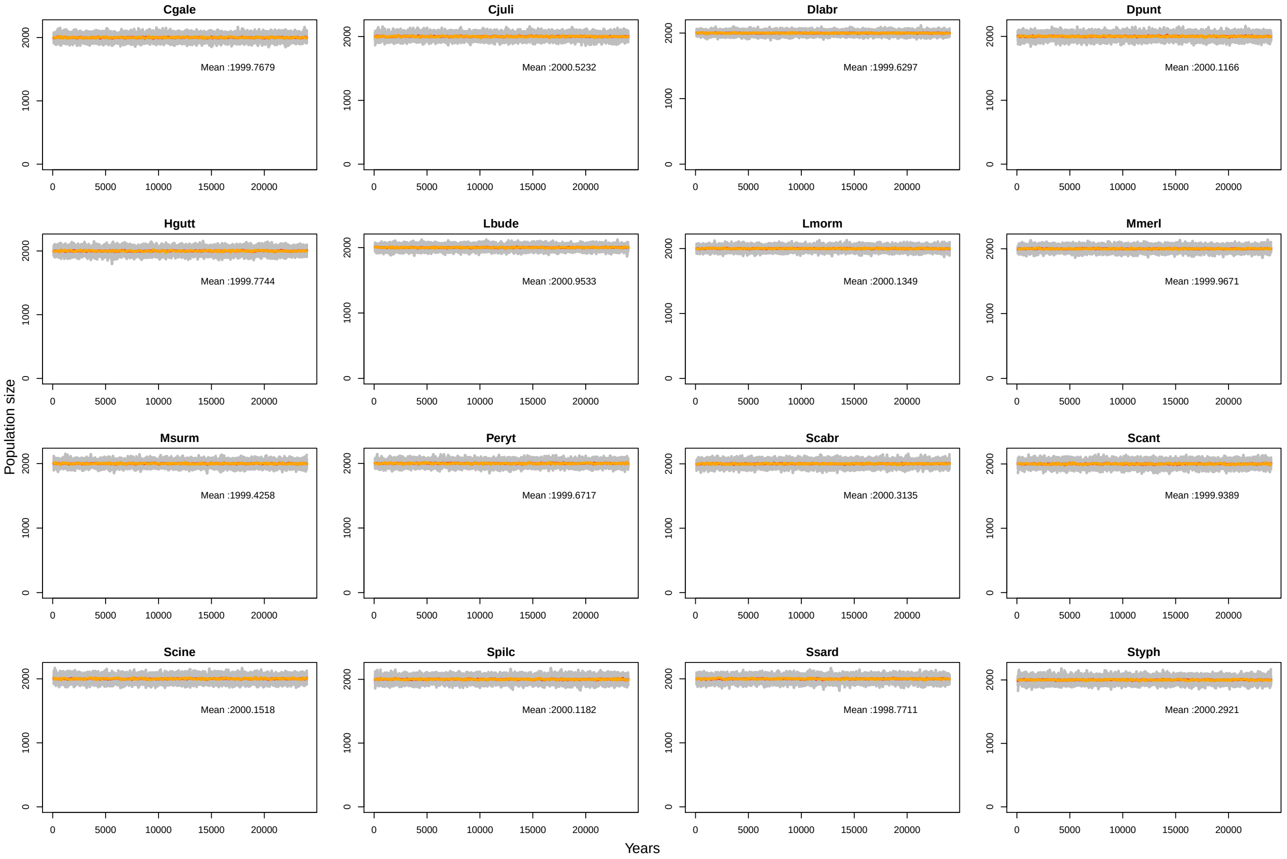

Figure S19: **Population size count for the 50 iterations of the 16 species for set 3 of life tables (age at first maturity at 1 year old, constant age-specific survival rate, increasing age-specific fecundity and no differences between sex-specific life tables)**. Adult population size was count for each iteration every 100 years during the 25000 years of the simulation. Each grey line represents population size fluctuations for one iteration. For each species, orange and red line, respectively, represents the median and the mean for all the 50 iterations. Mean over all 50 iterations is written on the plot for each species. Cgale = *Coryphoblennius galerita*, Cjuli = *Coris julis*, Dlabr = *Dicentrarchus labrax*, Dpunt = *Diplodus puntazzo*, Hgutt = *Hippocampus guttulatus*, Lbude = *Lophius budegassa*, Lmorm = *Lithognathus mormyrus*, Mmerl = *Merluccius merluccius*, Msurm = *Mullus surmuletus*, Peryt = *Pagellus erythrinus*, Scabr = *Serranus cabrilla*, Scant = *Spondyliosoma cantharus*, Scine = *Symphodus cinereus*, Spilc = *Sardina pilchardus*, Ssard = *Sarda sarda*, Styph = *Syngnathus typhle*

.

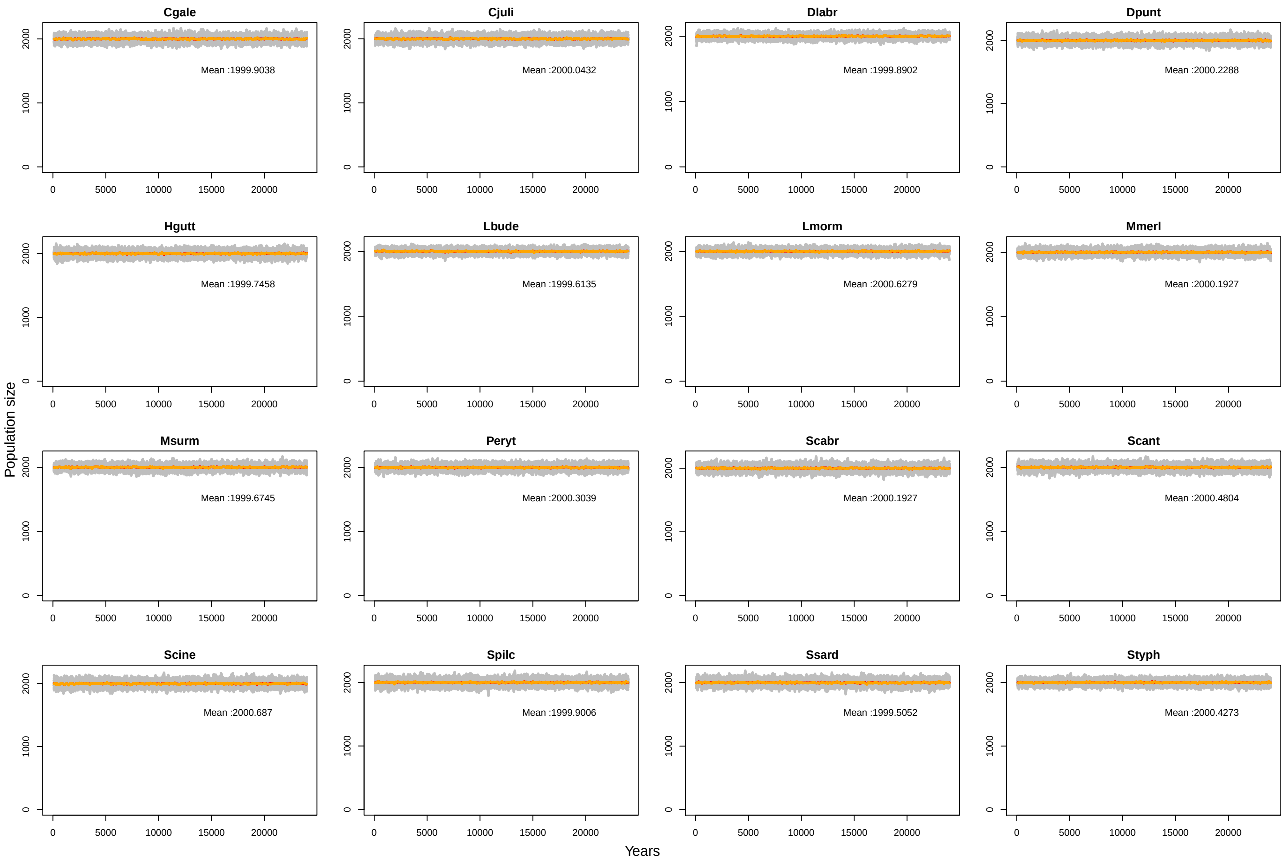

Figure S20: **Population size count for the 50 iterations of the 16 species for set 4 of life tables (age at first maturity at 1 year old, increasing age-specific survival rate, increasing age-specific fecundity and no differences between sex-specific life tables)**. Adult population size was count for each iteration every 100 years during the 25000 years of the simulation. Each grey line represents population size fluctuations for one iteration. For each species, orange and red line, respectively, represents the median and the mean for all the 50 iterations. Mean over all 50 iterations is written on the plot for each species. Cgale = *Coryphoblennius galerita*, Cjuli = *Coris julis*, Dlabr = *Dicentrarchus labrax*, Dpunt = *Diplodus puntazzo*, Hgutt = *Hippocampus guttulatus*, Lbude = *Lophius budegassa*, Lmorm = *Lithognathus mormyrus*, Mmerl = *Merluccius merluccius*, Msurm = *Mullus surmuletus*, Peryt = *Pagellus erythrinus*, Scabr = *Serranus cabrilla*, Scant = *Spondyliosoma cantharus*, Scine = *Symphodus cinereus*, Spilc = *Sardina pilchardus*, Ssard = *Sarda sarda*, Styph = *Syngnathus typhle*

.

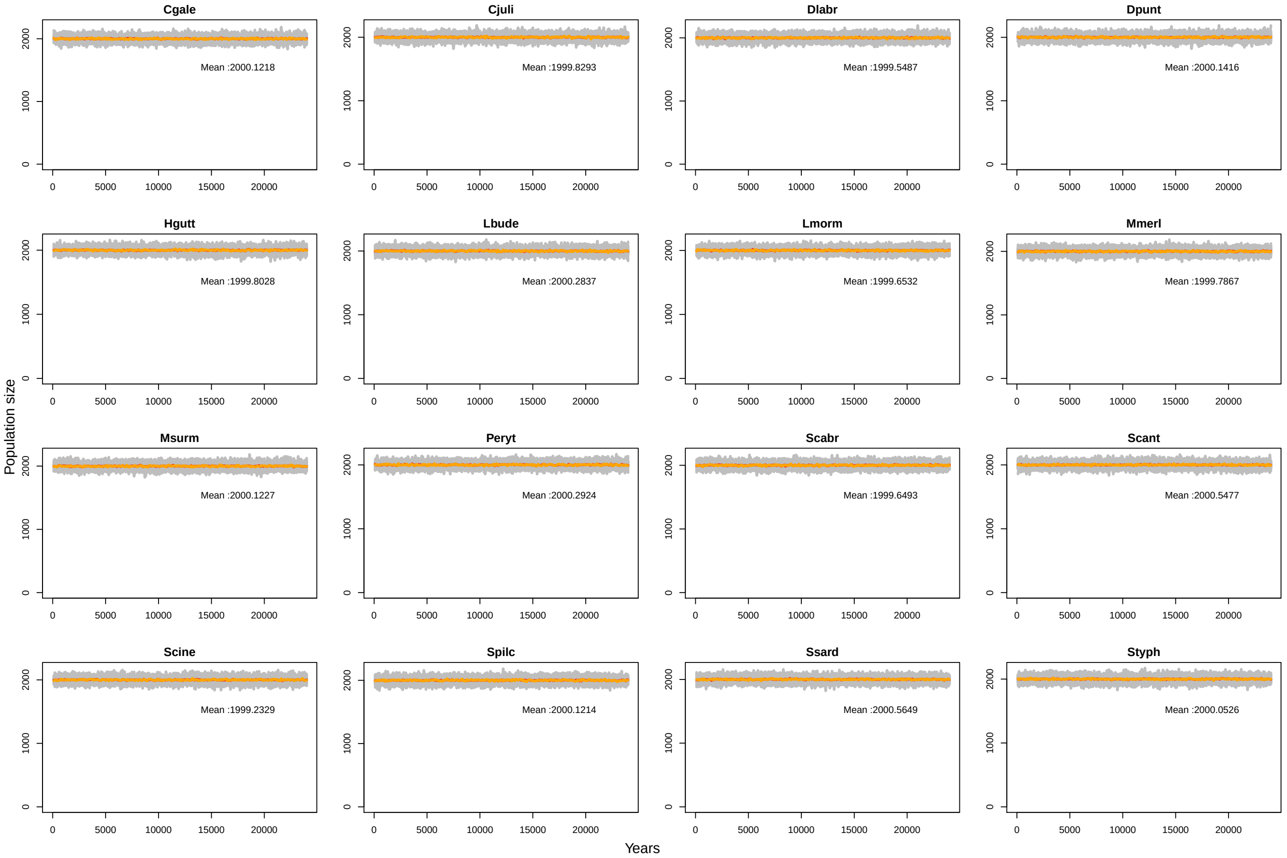

Figure S21: **Population size count for the 50 iterations of the 16 species for set 5 of life tables (age at first maturity at 1 year old, constant age-specific survival rate, constant age-specific fecundity and sex-specific differences in life tables)**. Adult population size was count for each iteration every 100 years during the 25000 years of the simulation. Each grey line represents population size fluctuations for one iteration. For each species, orange and red line, respectively, represents the median and the mean for all the 50 iterations. Mean over all 50 iterations is written on the plot for each species. Cgale = *Coryphoblennius galerita*, Cjuli = *Coris julis*, Dlabr = *Dicentrarchus labrax*, Dpunt = *Diplodus puntazzo*, Hgutt = *Hippocampus guttulatus*, Lbude = *Lophius budegassa*, Lmorm = *Lithognathus mormyrus*, Mmerl = *Merluccius merluccius*, Msurm = *Mullus surmuletus*, Peryt = *Pagellus erythrinus*, Scabr = *Serranus cabrilla*, Scant = *Spondyliosoma cantharus*, Scine = *Symphodus cinereus*, Spilc = *Sardina pilchardus*, Ssard = *Sarda sarda*, Styph = *Syngnathus typhle*

.

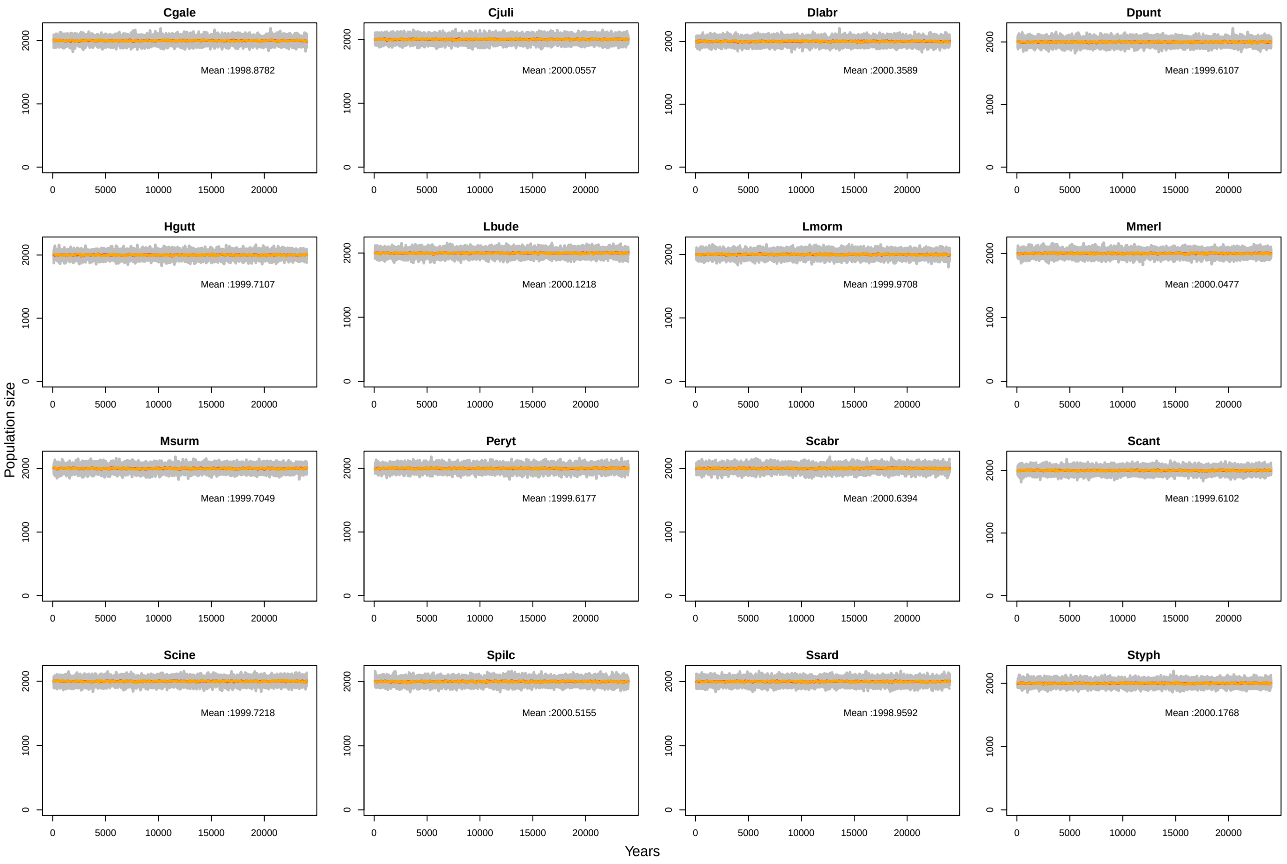

Figure S22: **Population size count for the 50 iterations of the 16 species for set 6 of life tables (age at first maturity at 1 year old, increasing age-specific survival rate, constant age-specific fecundity and sex-specific differences in life tables)**. Adult population size was count for each iteration every 100 years during the 25000 years of the simulation. Each grey line represents population size fluctuations for one iteration. For each species, orange and red line, respectively, represents the median and the mean for all the 50 iterations. Mean over all 50 iterations is written on the plot for each species. Cgale = *Coryphoblennius galerita*, Cjuli = *Coris julis*, Dlabr = *Dicentrarchus labrax*, Dpunt = *Diplodus puntazzo*, Hgutt = *Hippocampus guttulatus*, Lbude = *Lophius budegassa*, Lmorm = *Lithognathus mormyrus*, Mmerl = *Merluccius merluccius*, Msurm = *Mullus surmuletus*, Peryt = *Pagellus erythrinus*, Scabr = *Serranus cabrilla*, Scant = *Spondyliosoma cantharus*, Scine = *Symphodus cinereus*, Spilc = *Sardina pilchardus*, Ssard = *Sarda sarda*, Styph = *Syngnathus typhle*

.

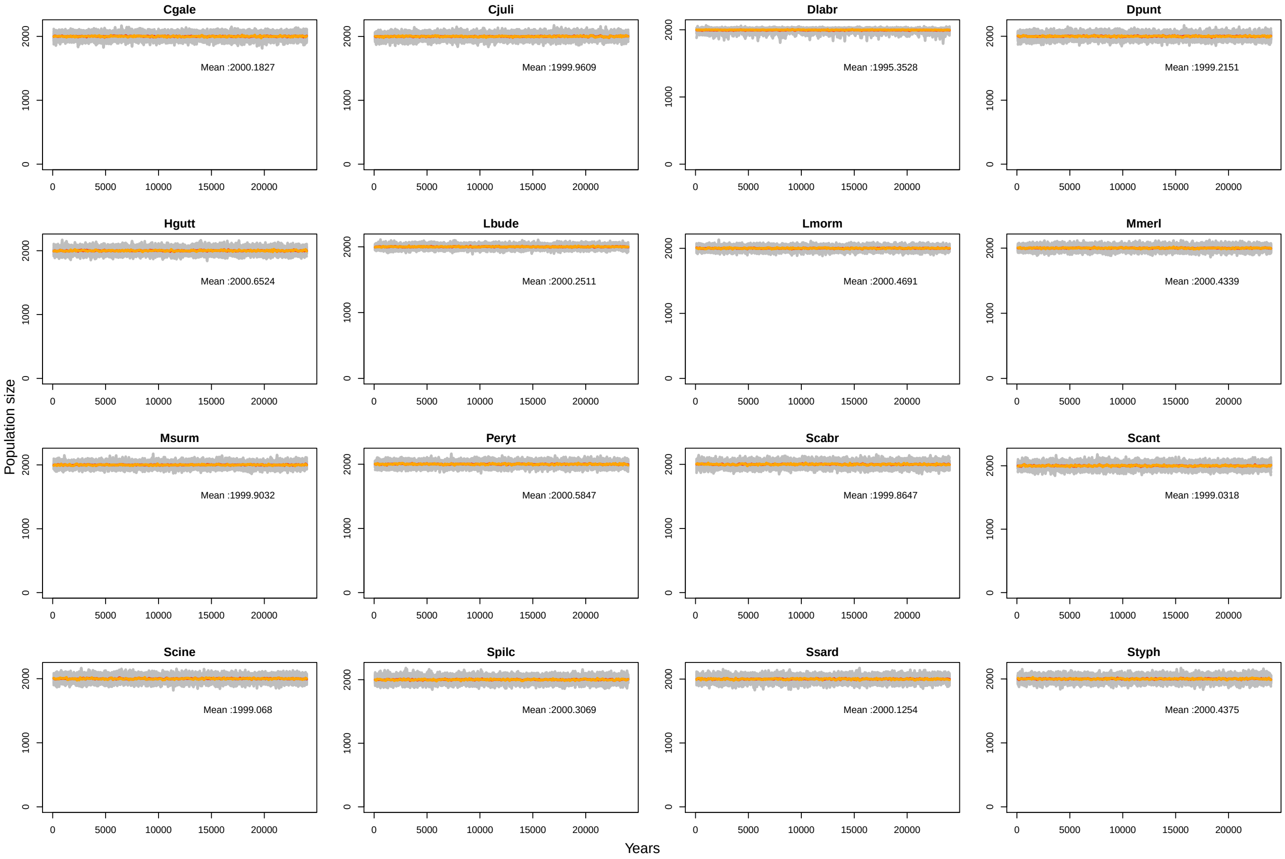

Figure S23: **Population size count for the 50 iterations of the 16 species for set 7 of life tables (age at first maturity at 1 year old, constant age-specific survival rate, increasing age-specific fecundity and sex-specific differences in life tables)**. Adult population size was count for each iteration every 100 years during the 25000 years of the simulation. Each grey line represents population size fluctuations for one iteration. For each species, orange and red line, respectively, represents the median and the mean for all the 50 iterations. Mean over all 50 iterations is written on the plot for each species. Cgale = *Coryphoblennius galerita*, Cjuli = *Coris julis*, Dlabr = *Dicentrarchus labrax*, Dpunt = *Diplodus puntazzo*, Hgutt = *Hippocampus guttulatus*, Lbude = *Lophius budegassa*, Lmorm = *Lithognathus mormyrus*, Mmerl = *Merluccius merluccius*, Msurm = *Mullus surmuletus*, Peryt = *Pagellus erythrinus*, Scabr = *Serranus cabrilla*, Scant = *Spondyliosoma cantharus*, Scine = *Symphodus cinereus*, Spilc = *Sardina pilchardus*, Ssard = *Sarda sarda*, Styph = *Syngnathus typhle*

.

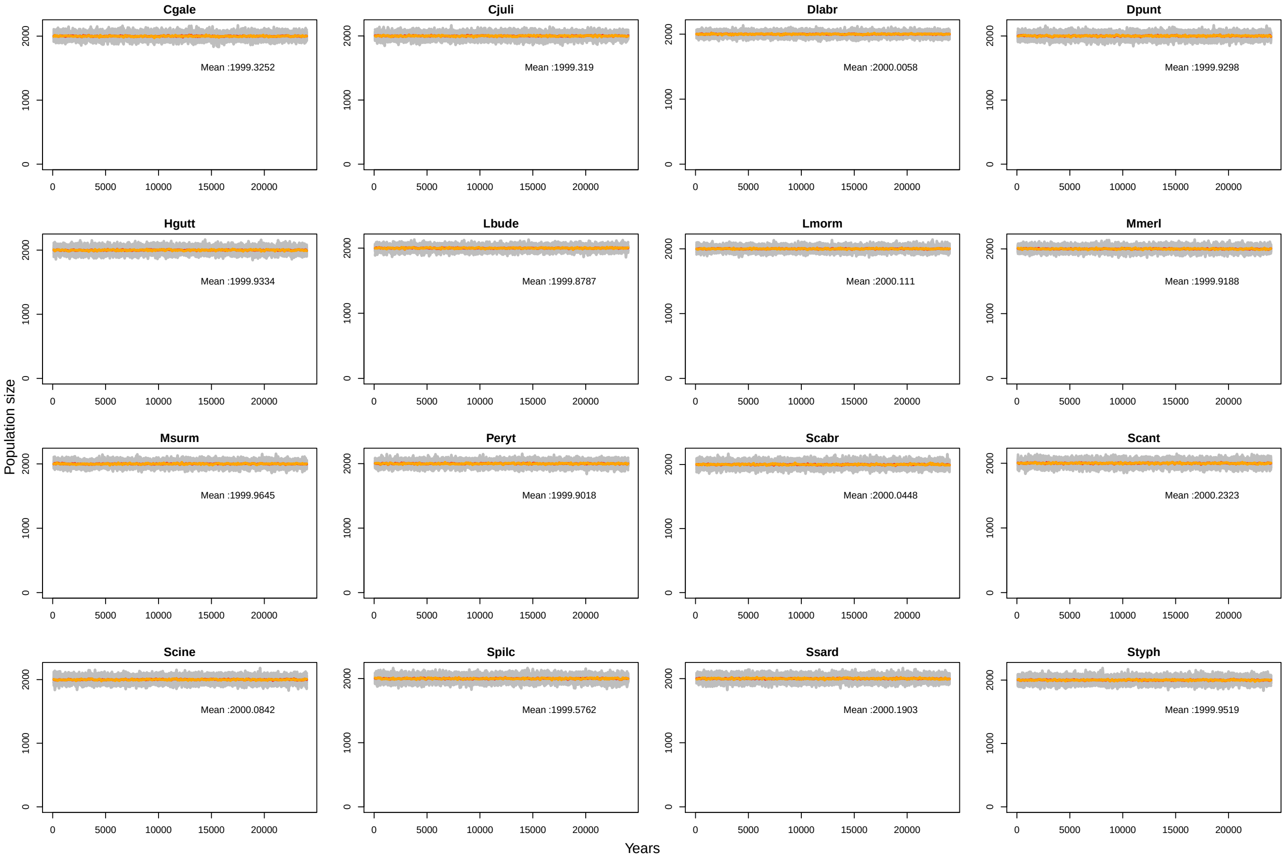

Figure S24: **Population size count for the 50 iterations of the 16 species for set 8 of life tables (age at first maturity at 1 year old, increasing age-specific survival rate, increasing age-specific fecundity and sex-specific differences in life tables)**. Adult population size was count for each iteration every 100 years during the 25000 years of the simulation. Each grey line represents population size fluctuations for one iteration. For each species, orange and red line, respectively, represents the median and the mean for all the 50 iterations. Mean over all 50 iterations is written on the plot for each species. Cgale = *Coryphoblennius galerita*, Cjuli = *Coris julis*, Dlabr = *Dicentrarchus labrax*, Dpunt = *Diplodus puntazzo*, Hgutt = *Hippocampus guttulatus*, Lbude = *Lophius budegassa*, Lmorm = *Lithognathus mormyrus*, Mmerl = *Merluccius merluccius*, Msurm = *Mullus surmuletus*, Peryt = *Pagellus erythrinus*, Scabr = *Serranus cabrilla*, Scant = *Spondyliosoma cantharus*, Scine = *Symphodus cinereus*, Spilc = *Sardina pilchardus*, Ssard = *Sarda sarda*, Styph = *Syngnathus typhle*

.

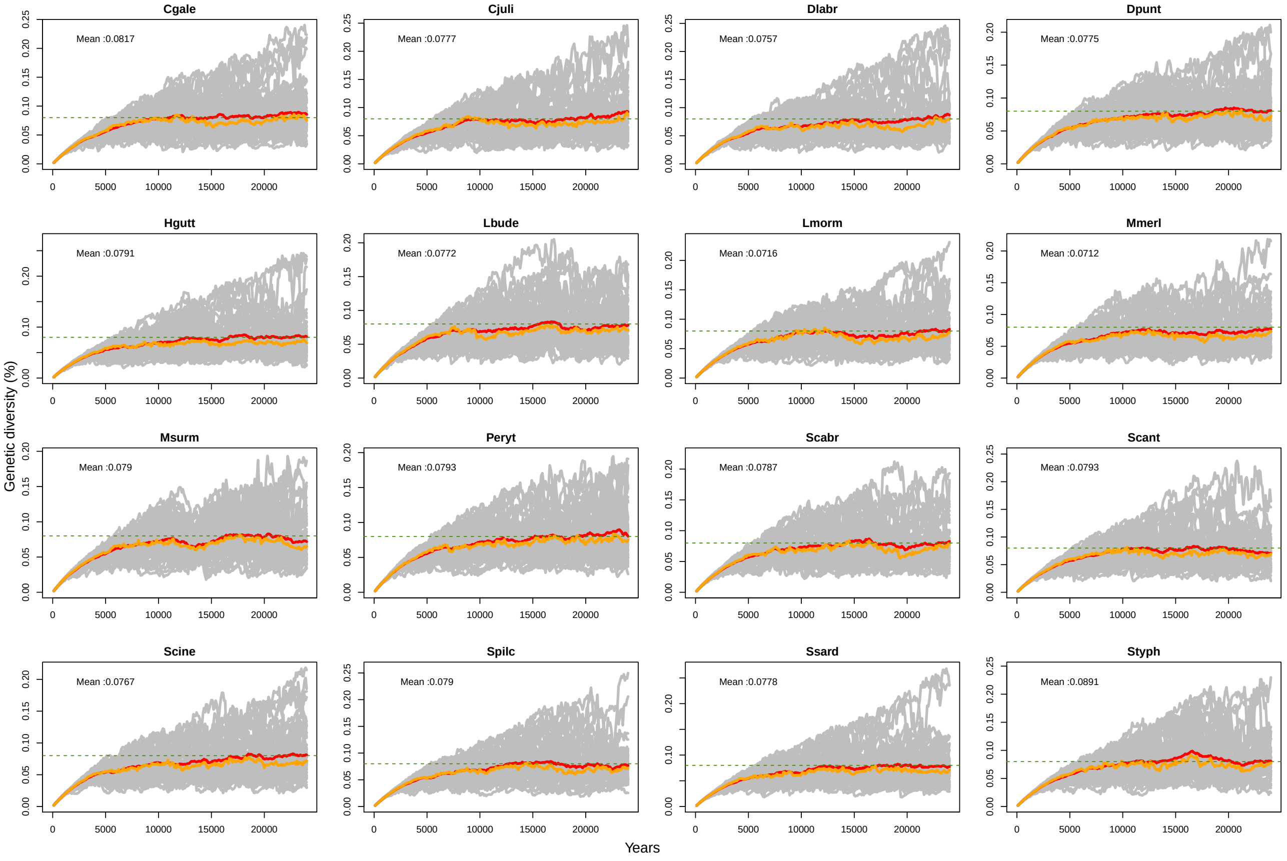

Figure S25: **Genetic diversity simulated for each of the 16 species for set 1 of life tables (age at first maturity at 1 year old, constant age-specific survival rate, constant age-specific fecundity and no differences between sex-specific life tables)**. Simulation were conducted with `SLiM` v.3.3.1 for 25000 years, with mutation rate, $\mu = 1e^{-7}$ on a 1 Mb non recombining loci, and carrying capacity equals $N = 2000$. Genetic diversity was estimated for each iteration every 100 years during the 25000 years of the simulation. Each grey line represents genetic diversity fluctuation for one iteration. For each species, orange and red line, respectively, represents the median and the mean for all the 50 iterations. Green dashed line represents the genetic diversity expected at mutation-drift equilibruim in a Wright-Fisher model ($4N\mu = 4 \times 2000 \times 1e^{-7} = 0.0008 = 0.08\%$). Mean over all 50 iterations, calculated with genetic diversity values estimated between 15 000 and 25 000 years after the beginning of the simulation, is written on the plot for each species. Cgale = *Coryphoblennius galerita*, Cjuli = *Coris julis*, Dlabr = *Dicentrarchus labrax*, Dpunt = *Diplodus puntazzo*, Hgutt = *Hippocampus guttulatus*, Lbude = *Lophius budegassa*, Lmorm = *Lithognathus mormyrus*, Mmerl = *Merluccius merluccius*, Msurm = *Mullus surmuletus*, Peryt = *Pagellus erythrinus*, Scabr = *Serranus cabrilla*, Scant = *Spondyliosoma cantharus*, Scine = *Symphodus cinereus*, Spilc = *Sardina pilchardus*, Ssard = *Sarda sarda*, Styph = *Syngnathus typhle*

.

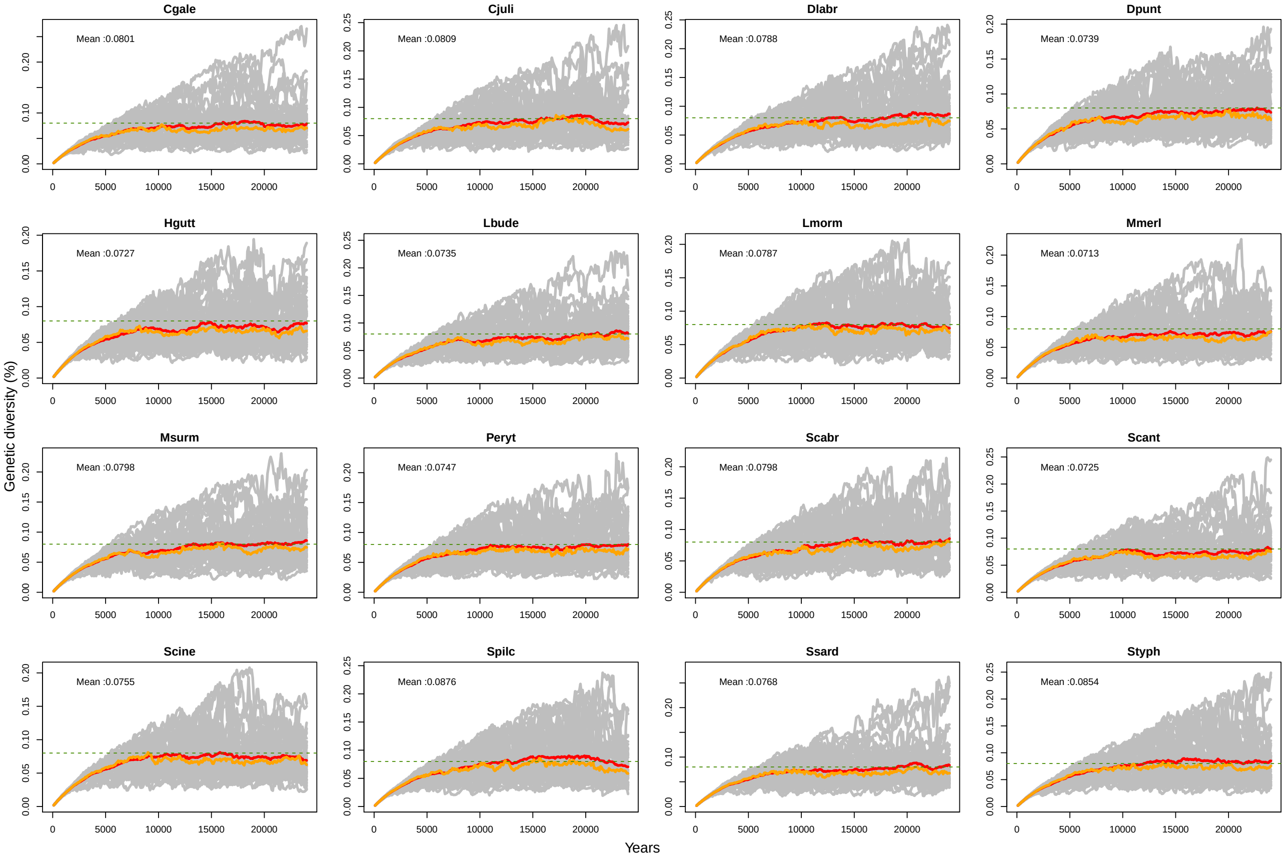

Figure S26: **Genetic diversity simulated for each of the 16 species for set 2 of life tables (age at first maturity at 1 year old, increasing age-specific survival rate, constant age-specific fecundity and no differences between sex-specific life tables)**. Simulation were conducted with `SLiM` v.3.3.1 for 25000 years, with mutation rate, $\mu = 1e^{-7}$ on a 1 Mb non recombining loci, and carrying capacity equals $N = 2000$. Genetic diversity was estimated for each iteration every 100 years during the 25000 years of the simulation. Each grey line represents genetic diversity fluctuation for one iteration. For each species, orange and red line, respectively, represents the median and the mean for all the 50 iterations. Green dashed line represents the genetic diversity expected at mutation-drift equilibruim in a Wright-Fisher model ($4N\mu = 4 \times 2000 \times 1e^{-7} = 0.0008 = 0.08\%$). Mean over all 50 iterations, calculated with genetic diversity values estimated between 15 000 and 25 000 years after the beginning of the simulation, is written on the plot for each species. Cgale = *Coryphoblennius galerita*, Cjuli = *Coris julis*, Dlabr = *Dicentrarchus labrax*, Dpunt = *Diplodus puntazzo*, Hgutt = *Hippocampus guttulatus*, Lbude = *Lophius budegassa*, Lmorm = *Lithognathus mormyrus*, Mmerl = *Merluccius merluccius*, Msurm = *Mullus surmuletus*, Peryt = *Pagellus erythrinus*, Scabr = *Serranus cabrilla*, Scant = *Spondyliosoma cantharus*, Scine = *Symphodus cinereus*, Spilc = *Sardina pilchardus*, Ssard = *Sarda sarda*, Styph = *Syngnathus typhle*

.

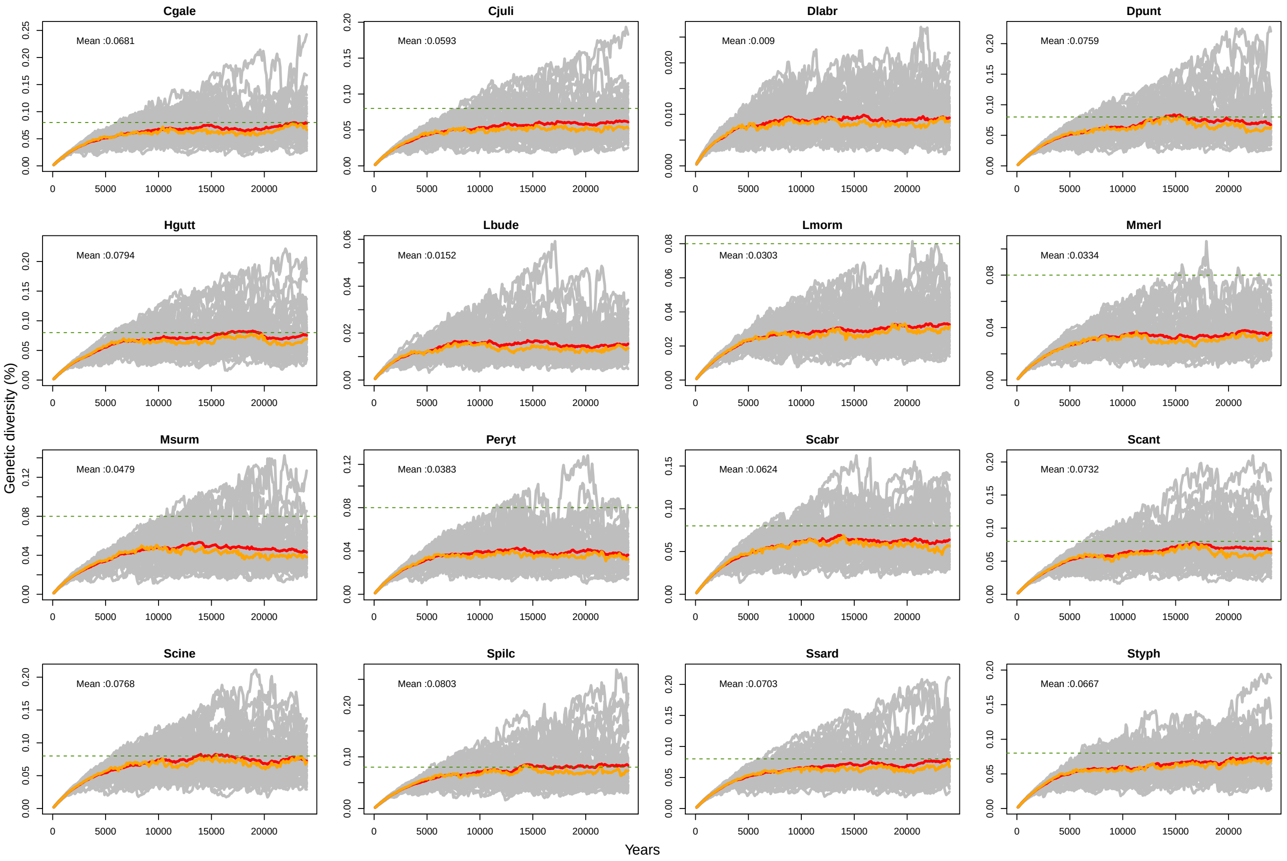

Figure S27: **Genetic diversity simulated for each of the 16 species for set 3 of life tables (age at first maturity at 1 year old, constant age-specific survival rate, increasing age-specific fecundity and no differences between sex-specific life tables)**. Simulation were conducted with `SLiM` v.3.3.1 for 25000 years, with mutation rate, $\mu = 1e^{-7}$ on a 1 Mb non recombining loci, and carrying capacity equals $N = 2000$. Genetic diversity was estimated for each iteration every 100 years during the 25000 years of the simulation. Each grey line represents genetic diversity fluctuation for one iteration. For each species, orange and red line, respectively, represents the median and the mean for all the 50 iterations. Green dashed line represents the genetic diversity expected at mutation-drift equilibruim in a Wright-Fisher model ($4N\mu = 4 \times 2000 \times 1e^{-7} = 0.0008 = 0.08\%$). Mean over all 50 iterations, calculated with genetic diversity values estimated between 15 000 and 25 000 years after the beginning of the simulation, is written on the plot for each species. Cgale = *Coryphoblennius galerita*, Cjuli = *Coris julis*, Dlabr = *Dicentrarchus labrax*, Dpunt = *Diplodus puntazzo*, Hgutt = *Hippocampus guttulatus*, Lbude = *Lophius budegassa*, Lmorm = *Lithognathus mormyrus*, Mmerl = *Merluccius merluccius*, Msurm = *Mullus surmuletus*, Peryt = *Pagellus erythrinus*, Scabr = *Serranus cabrilla*, Scant = *Spondyliosoma cantharus*, Scine = *Symphodus cinereus*, Spilc = *Sardina pilchardus*, Ssard = *Sarda sarda*, Styph = *Syngnathus typhle*

.

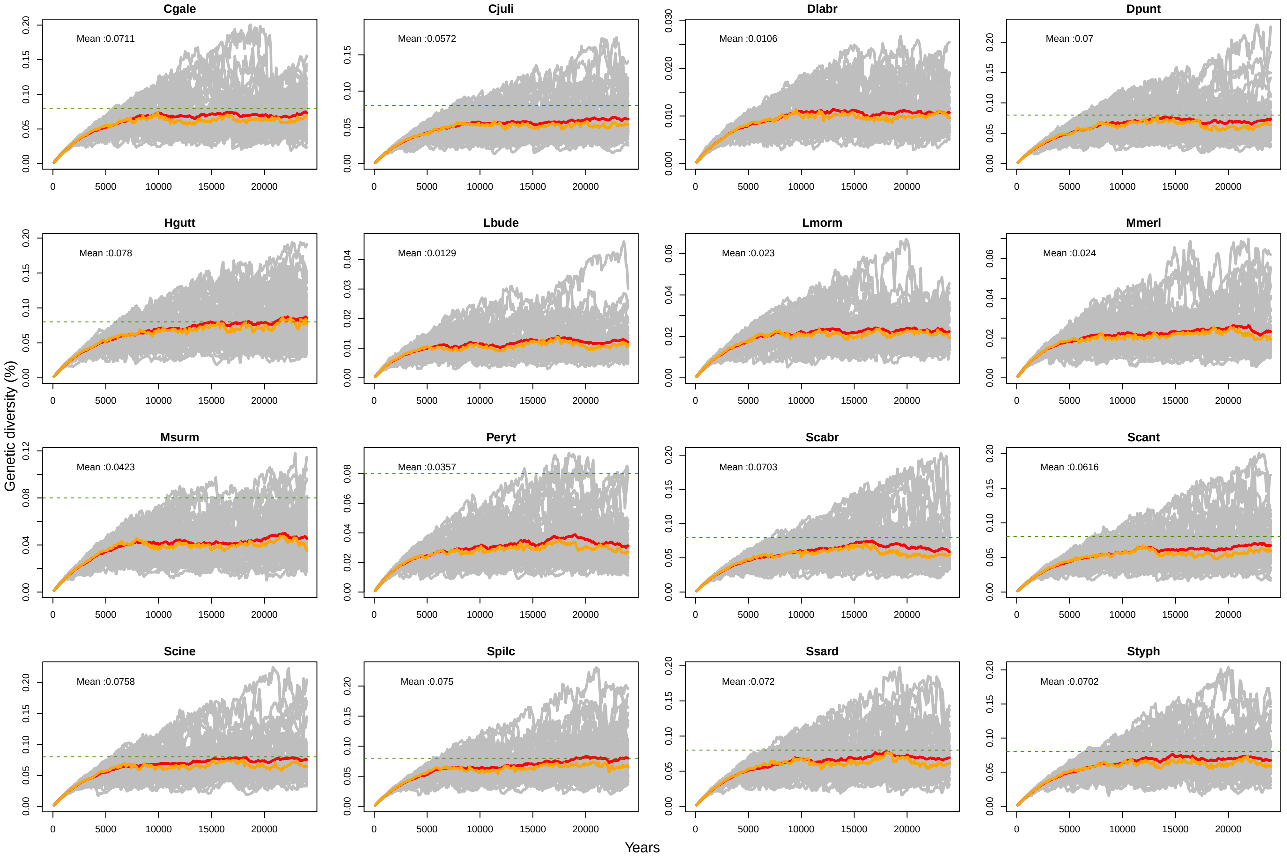

Figure S28: **Genetic diversity simulated for each of the 16 species for set 4 of life tables (age at first maturity at 1 year old, increasing age-specific survival rate, increasing age-specific fecundity and no differences between sex-specific life tables)**. Simulation were conducted with `SLiM` v.3.3.1 for 25000 years, with mutation rate, $\mu = 1e^{-7}$ on a 1 Mb non recombining loci, and carrying capacity equals $N = 2000$. Genetic diversity was estimated for each iteration every 100 years during the 25000 years of the simulation. Each grey line represents genetic diversity fluctuation for one iteration. For each species, orange and red line, respectively, represents the median and the mean for all the 50 iterations. Green dashed line represents the genetic diversity expected at mutation-drift equilibruim in a Wright-Fisher model ($4N\mu = 4 \times 2000 \times 1e^{-7} = 0.0008 = 0.08\%$). Mean over all 50 iterations, calculated with genetic diversity values estimated between 15 000 and 25 000 years after the beginning of the simulation, is written on the plot for each species. Cgale = *Coryphoblennius galerita*, Cjuli = *Coris julis*, Dlabr = *Dicentrarchus labrax*, Dpunt = *Diplodus puntazzo*, Hgutt = *Hippocampus guttulatus*, Lbude = *Lophius budegassa*, Lmorm = *Lithognathus mormyrus*, Mmerl = *Merluccius merluccius*, Msurm = *Mullus surmuletus*, Peryt = *Pagellus erythrinus*, Scabr = *Serranus cabrilla*, Scant = *Spondyliosoma cantharus*, Scine = *Symphodus cinereus*, Spilc = *Sardina pilchardus*, Ssard = *Sarda sarda*, Styph = *Syngnathus typhle*

.

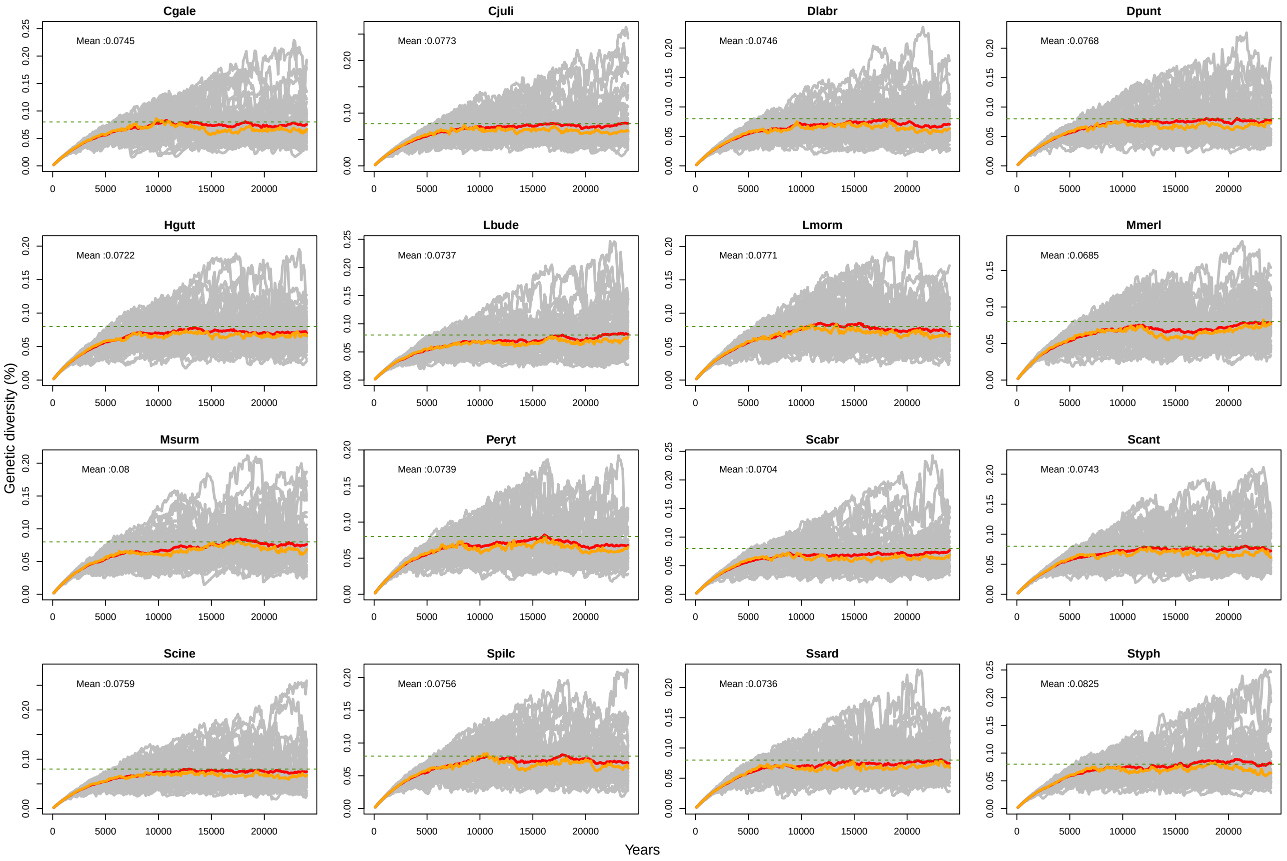

Figure S29: **Genetic diversity simulated for each of the 16 species for set 5 of life tables (age at first maturity at 1 year old, constant age-specific survival rate, constant age-specific fecundity and sex-specific differences in life tables)**. Simulation were conducted with `SLiM` v.3.3.1 for 25000 years, with mutation rate, $\mu = 1e^{-7}$ on a 1 Mb non recombining loci, and carrying capacity equals $N = 2000$. Genetic diversity was estimated for each iteration every 100 years during the 25000 years of the simulation. Each grey line represents genetic diversity fluctuation for one iteration. For each species, orange and red line, respectively, represents the median and the mean for all the 50 iterations. Green dashed line represents the genetic diversity expected at mutation-drift equilibruim in a Wright-Fisher model ($4N\mu = 4 \times 2000 \times 1e^{-7} = 0.0008 = 0.08\%$). Mean over all 50 iterations, calculated with genetic diversity values estimated between 15 000 and 25 000 years after the beginning of the simulation, is written on the plot for each species. Cgale = *Coryphoblennius galerita*, Cjuli = *Coris julis*, Dlabr = *Dicentrarchus labrax*, Dpunt = *Diplodus puntazzo*, Hgutt = *Hippocampus guttulatus*, Lbude = *Lophius budegassa*, Lmorm = *Lithognathus mormyrus*, Mmerl = *Merluccius merluccius*, Msurm = *Mullus surmuletus*, Peryt = *Pagellus erythrinus*, Scabr = *Serranus cabrilla*, Scant = *Spondyliosoma cantharus*, Scine = *Symphodus cinereus*, Spilc = *Sardina pilchardus*, Ssard = *Sarda sarda*, Styph = *Syngnathus typhle*

.

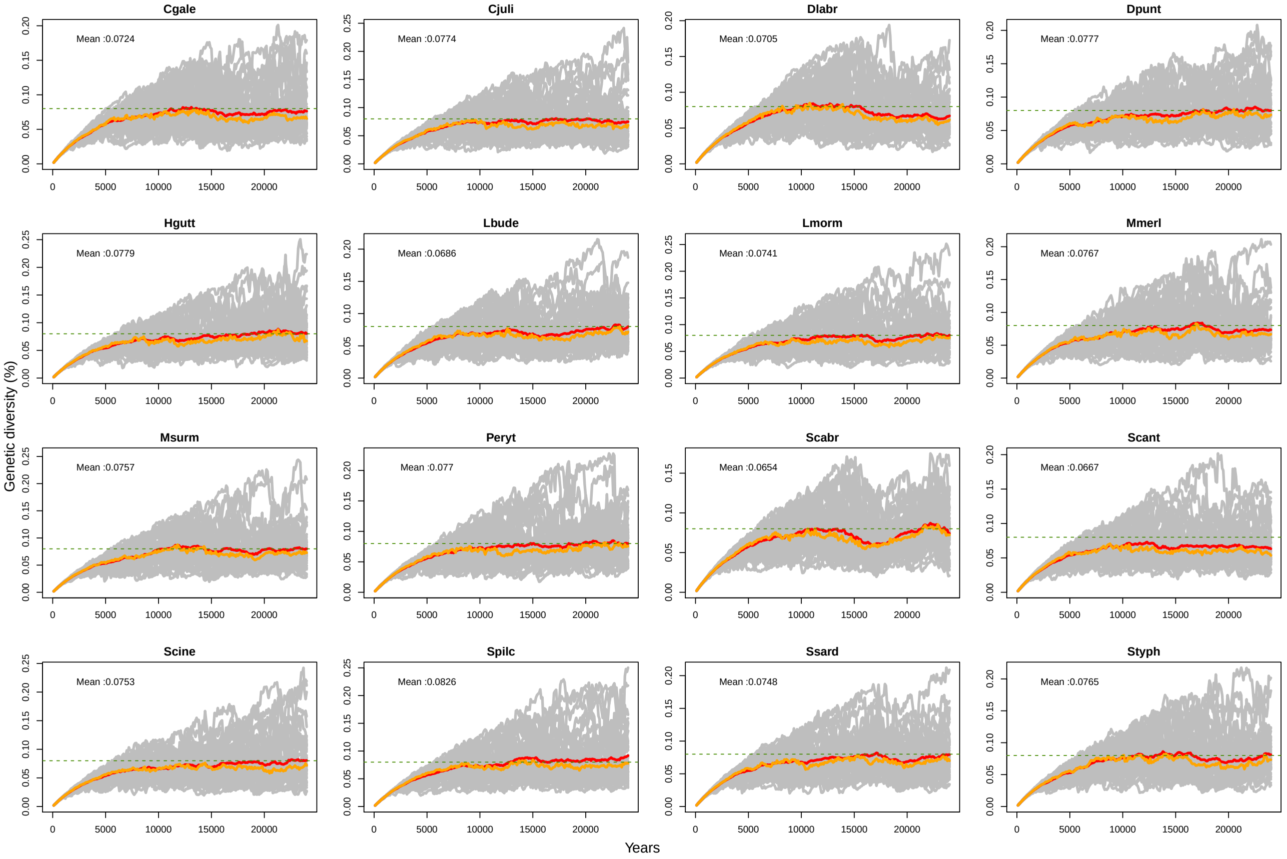

Figure S30: **Genetic diversity simulated for each of the 16 species for set 6 of life tables (age at first maturity at 1 year old, increasing age-specific survival rate, constant age-specific fecundity and sex-specific differences in life tables)**. Simulation were conducted with `SLiM` v.3.3.1 for 25000 years, with mutation rate, $\mu = 1e^{-7}$ on a 1 Mb non recombining loci, and carrying capacity equals $N = 2000$. Genetic diversity was estimated for each iteration every 100 years during the 25000 years of the simulation. Each grey line represents genetic diversity fluctuation for one iteration. For each species, orange and red line, respectively, represents the median and the mean for all the 50 iterations. Green dashed line represents the genetic diversity expected at mutation-drift equilibruim in a Wright-Fisher model ($4N\mu = 4 \times 2000 \times 1e^{-7} = 0.0008 = 0.08\%$). Mean over all 50 iterations, calculated with genetic diversity values estimated between 15 000 and 25 000 years after the beginning of the simulation, is written on the plot for each species. Cgale = *Coryphoblennius galerita*, Cjuli = *Coris julis*, Dlabr = *Dicentrarchus labrax*, Dpunt = *Diplodus puntazzo*, Hgutt = *Hippocampus guttulatus*, Lbude = *Lophius budegassa*, Lmorm = *Lithognathus mormyrus*, Mmerl = *Merluccius merluccius*, Msurm = *Mullus surmuletus*, Peryt = *Pagellus erythrinus*, Scabr = *Serranus cabrilla*, Scant = *Spondyliosoma cantharus*, Scine = *Symphodus cinereus*, Spilc = *Sardina pilchardus*, Ssard = *Sarda sarda*, Styph = *Syngnathus typhle*

.

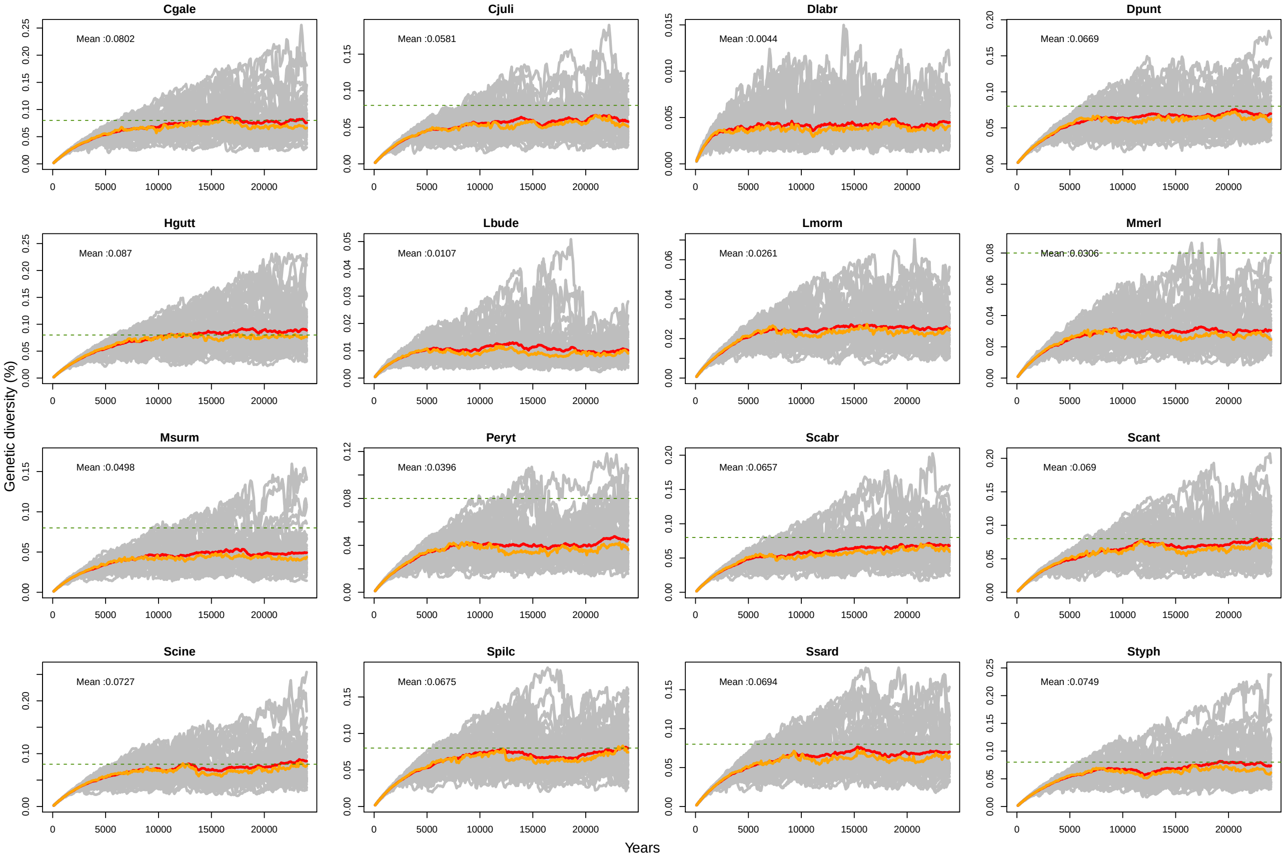

Figure S31: **Genetic diversity simulated for each of the 16 species for set 7 of life tables (age at first maturity at 1 year old, constant age-specific survival rate, increasing age-specific fecundity and sex-specific differences in life tables)**. Simulation were conducted with `SLiM` v.3.3.1 for 25000 years, with mutation rate, $\mu = 1e^{-7}$ on a 1 Mb non recombining loci, and carrying capacity equals $N = 2000$. Genetic diversity was estimated for each iteration every 100 years during the 25000 years of the simulation. Each grey line represents genetic diversity fluctuation for one iteration. For each species, orange and red line, respectively, represents the median and the mean for all the 50 iterations. Green dashed line represents the genetic diversity expected at mutation-drift equilibruim in a Wright-Fisher model ($4N\mu = 4 \times 2000 \times 1e^{-7} = 0.0008 = 0.08\%$). Mean over all 50 iterations, calculated with genetic diversity values estimated between 15 000 and 25 000 years after the beginning of the simulation, is written on the plot for each species. Cgale = *Coryphoblennius galerita*, Cjuli = *Coris julis*, Dlabr = *Dicentrarchus labrax*, Dpunt = *Diplodus puntazzo*, Hgutt = *Hippocampus guttulatus*, Lbude = *Lophius budegassa*, Lmorm = *Lithognathus mormyrus*, Mmerl = *Merluccius merluccius*, Msurm = *Mullus surmuletus*, Peryt = *Pagellus erythrinus*, Scabr = *Serranus cabrilla*, Scant = *Spondyliosoma cantharus*, Scine = *Symphodus cinereus*, Spilc = *Sardina pilchardus*, Ssard = *Sarda sarda*, Styph = *Syngnathus typhle*

.

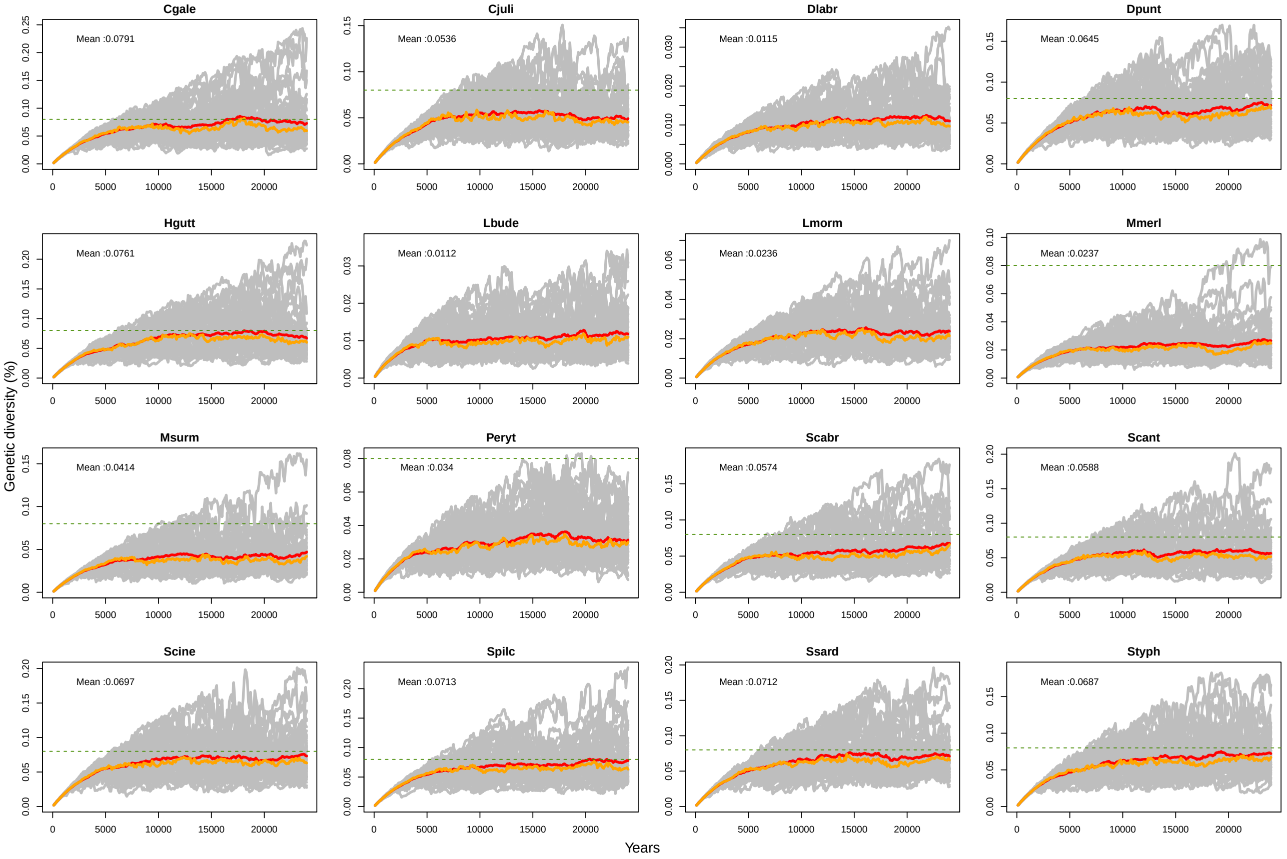

Figure S32: **Genetic diversity simulated for each of the 16 species for set 8 of life tables (age at first maturity at 1 year old, increasing age-specific survival rate, increasing age-specific fecundity and sex-specific differences in life tables)**. Simulation were conducted with `SLiM` v.3.3.1 for 25000 years, with mutation rate, $\mu = 1e^{-7}$ on a 1 Mb non recombining loci, and carrying capacity equals $N = 2000$. Genetic diversity was estimated for each iteration every 100 years during the 25000 years of the simulation. Each grey line represents genetic diversity fluctuation for one iteration. For each species, orange and red line, respectively, represents the median and the mean for all the 50 iterations. Green dashed line represents the genetic diversity expected at mutation-drift equilibruim in a Wright-Fisher model ($4N\mu = 4 \times 2000 \times 1e^{-7} = 0.0008 = 0.08\%$). Mean over all 50 iterations, calculated with genetic diversity values estimated between 15 000 and 25 000 years after the beginning of the simulation, is written on the plot for each species. Cgale = *Coryphoblennius galerita*, Cjuli = *Coris julis*, Dlabr = *Dicentrarchus labrax*, Dpunt = *Diplodus puntazzo*, Hgutt = *Hippocampus guttulatus*, Lbude = *Lophius budegassa*, Lmorm = *Lithognathus mormyrus*, Mmerl = *Merluccius merluccius*, Msurm = *Mullus surmuletus*, Peryt = *Pagellus erythrinus*, Scabr = *Serranus cabrilla*, Scant = *Spondyliosoma cantharus*, Scine = *Symphodus cinereus*, Spilc = *Sardina pilchardus*, Ssard = *Sarda sarda*, Styph = *Syngnathus typhle*

.
