## Appendix for "Age-specific survivorship and fecundity shape genetic diversity in marine fishes"

### Appendix 1 to Barry and al. "Life tables shape genetic diversity in marine fishes": life tables of the 16 marine teleostean species

Pierre Barry, Thomas Broquet, Pierre-Alexandre Gagnaire

Each page represents the informations on life tables and corresponding bibliographic references for each of the 16 species. On top, the image from Iglésias[13], reproduced with permissions, vernacular and Latin name of the species.

The first table, on top, shows length at infinity $L_{inf}$, growth parameter $K$, and $t_0$ from Von Bertalanffy equation modeling age-length relationship. Maturity show age at first maturity, and maximum age, lifespan. Values are shown for males, females and for combined sexes. On bottom of the first table, $F = y(L)$ shows the corresponding model between age and fecundity, either linear ($F = \alpha + \beta \times L$), exponential ($F = \alpha \times exp[\beta \times L]$) or power-law ($F = \alpha \times L^{\beta}$). $\alpha$ and $\beta$ show the corresponding parameter of the relationship between age and fecundity. Corresponding bibliographic references are indicated in the last column.

The second table shows life tables components calculated with the parameters of the first table: for each age, $L_x$ indicates length in centimeters for combined sexes, $L_{x,f}$, length for females only and $L_{x,m}$, length for males only. $l_x$, $l_{x,f}$ and $l_{x,m}$ indicates age-specific survival for combined sexes, females and males, respectively. $Cum_c$, $Cum_f$ and $Cum_m$ indicates cumulative age-specific survival for combined sexes, females and males respectively. $B_x$, $B_{x,f}$ and $B_{x,m}$ show relative fecundity for combined sexes, females and males, respectively (max fecundity equals 1 at lifespan).

On the bottom, solid lines represent the age-specific probability of survival until age $x$ curves ($y$ left-axis), dashed lines represent the age-specific fecundity curves ($y$ right-axis). Black, blue and red lines represent combined sexes, male and females curves, respectively.

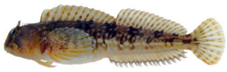

### Montagu's blenny

### *Coryphoblennius galerita*

| Parameter | Combined | Female | Male | Ref |
|---|---|---|---|---|
| $L_{inf}$ | 6.54 | 6.54 | 6.54 | Milton[16] |
| $K$ | 0.432 | 0.432 | 0.432 | |
| $t_0$ | -1.247 | -1.247 | -1.247 | |
| Maturity | 1 | 1 | 1 | |
| Max age | 6 | 6 | 6 | |
| $F = y(L)$ | Linear | | | Carrassón and Bau[5] |
| $\alpha$ | -2146.4 | | | |
| $\beta$ | 710.07 | | | |

| Age | $L_x$ | $L_{x,f}$ | $L_{x,m}$ | $l_x$ | $l_{x,f}$ | $l_{x,m}$ | $Cum_c$ | $Cum_f$ | $Cum_m$ | $B_x$ | $B_{x,f}$ | $B_{x,m}$ |
|---|---|---|---|---|---|---|---|---|---|---|---|---|
| 1 | 4.4 | 4.4 | 4.4 | 0.46 | 0.46 | 0.46 | 0.180 | 0.180 | 0.180 | 0.420 | 0.420 | 0.420 |
| 2 | 4.8 | 4.8 | 4.8 | 0.50 | 0.50 | 0.50 | 0.083 | 0.083 | 0.083 | 0.542 | 0.542 | 0.542 |
| 3 | 5.3 | 5.3 | 5.3 | 0.55 | 0.55 | 0.55 | 0.041 | 0.041 | 0.041 | 0.695 | 0.695 | 0.695 |
| 4 | 5.8 | 5.8 | 5.8 | 0.60 | 0.60 | 0.60 | 0.023 | 0.023 | 0.023 | 0.847 | 0.847 | 0.847 |
| 5 | 6.2 | 6.2 | 6.2 | 0.63 | 0.63 | 0.63 | 0.013 | 0.013 | 0.013 | 0.969 | 0.969 | 0.969 |
| 6 | 6.3 | 6.3 | 6.3 | 0.00 | 0.00 | 0.00 | 0.008 | 0.008 | 0.008 | 1.00 | 1.00 | 1.00 |

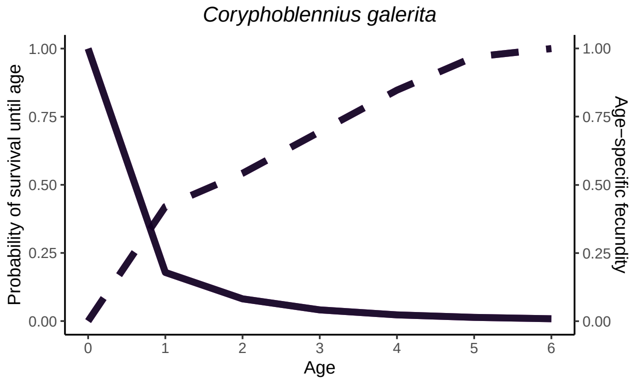

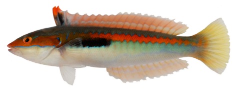

### Rainbow wrasse

### *Coris julis*

| Parameter | Combined | Female | Male | Ref |
|---|---|---|---|---|
| $L_{inf}$ | 25.4 | 21.27 | 29.66 | Škeljo [25] |
| $K$ | 0.16 | 0.21 | 0.12 | |
| $t_0$ | -1.19 | -1.08 | -1.52 | |
| Maturity | 1 | 1 | 1 | |
| Max age | 7 | 5 | 6 | |
| $F = y(L)$ | Power | | | Škeljo [25] |
| $\alpha$ | 0.902 | | | |
| $\beta$ | 3.643 | | | |

| Age | $L_x$ | $L_{x,f}$ | $L_{x,m}$ | $l_x$ | $l_{x,f}$ | $l_{x,m}$ | $Cum_c$ | $Cum_f$ | $Cum_m$ | $B_x$ | $B_{x,f}$ | $B_{x,m}$ |
|---|---|---|---|---|---|---|---|---|---|---|---|---|
| 1 | 8.5 | 8.7 | 8.5 | 0.44 | 0.45 | 0.46 | 0.282 | 0.305 | 0.327 | 0.037 | 0.119 | 0.037 |
| 2 | 11.4 | 11.3 | 11.5 | 0.59 | 0.58 | 0.61 | 0.123 | 0.137 | 0.150 | 0.108 | 0.309 | 0.112 |
| 3 | 12.9 | 12.6 | 13.1 | 0.64 | 0.63 | 0.66 | 0.073 | 0.079 | 0.091 | 0.169 | 0.459 | 0.179 |
| 4 | 14.8 | 14.9 | 14.8 | 0.70 | 0.70 | 0.71 | 0.047 | 0.050 | 0.060 | 0.280 | 0.846 | 0.280 |
| 5 | 16.5 | 15.6 | 16.6 | 0.74 | 0.72 | 0.75 | 0.033 | 0.035 | 0.043 | 0.415 | 1.000 | 0.425 |
| 6 | 18.1 | – | 18.1 | 0.77 | – | 0.78 | 0.024 | 0.025 | 0.032 | 0.582 | – | 0.582 |
| 7 | 21 | – | 21 | 0.81 | – | 0.82 | 0.018 | – | 0.025 | 1.000 | – | 1.000 |

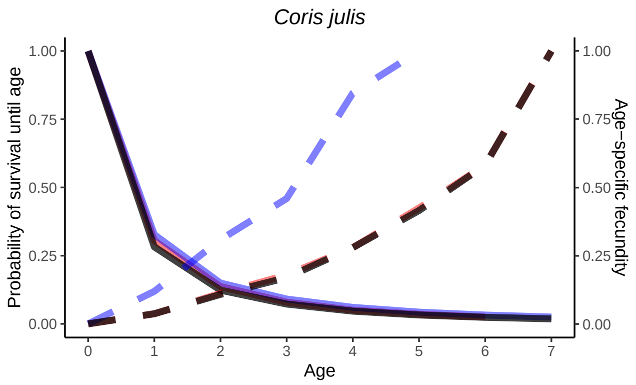

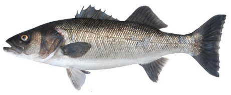

### European sea bass

#### *Dicentrarchus labrax*

| Parameter | Combined | Female | Male | Ref |
|---|---|---|---|---|
| $L_{inf}$ | 83.2 | 87.8 | 78.1 | Wassef and Emary[28] |
| $K$ | 0.066 | 0.061 | 0.075 | |
| $t_0$ | -1.745 | -1.797 | -1.765 | |
| Maturity | 3 | 2 | 4 | |
| Max age | 15 | 15 | 9 | |
| $F = y(L)$ | Power | | | Kara[14] |
| $\alpha$ | 0.00087 | | | |
| $\beta$ | 5 | | | |

| Age | $L_x$ | $L_{x,f}$ | $L_{x,m}$ | $l_x$ | $l_{x,f}$ | $l_{x,m}$ | $Cum_c$ | $Cum_f$ | $Cum_m$ | $B_x$ | $B_{x,f}$ | $B_{x,m}$ |
|---|---|---|---|---|---|---|---|---|---|---|---|---|
| 1 | 13.3 | 13.3 | 13.6 | 0.36 | 0.36 | 0.36 | 0.376 | 0.162 | 0.179 | 0.000 | 0.000 | 0.000 |
| 2 | 18.1 | 18.1 | 18.2 | 0.52 | 0.52 | 0.51 | 0.134 | 0.057 | 0.064 | 0.000 | 0.000 | 0.000 |
| 3 | 21.8 | 21.8 | 21.7 | 0.61 | 0.61 | 0.60 | 0.070 | 0.030 | 0.033 | 0.000 | 0.000 | 0.028 |
| 4 | 25.5 | 25.5 | 25.4 | 0.68 | 0.68 | 0.67 | 0.043 | 0.018 | 0.020 | 0.008 | 0.000 | 0.062 |
| 5 | 29.5 | 29.5 | 29.4 | 0.73 | 0.73 | 0.72 | 0.029 | 0.012 | 0.013 | 0.017 | 0.017 | 0.129 |
| 6 | 33.2 | 33.2 | 33 | 0.77 | 0.77 | 0.76 | 0.021 | 0.009 | 0.009 | 0.031 | 0.031 | 0.229 |
| 7 | 36.5 | 36.5 | 36.2 | 0.80 | 0.80 | 0.79 | 0.016 | 0.007 | 0.007 | 0.051 | 0.051 | 0.364 |
| 8 | 40.3 | 40.3 | 40.3 | 0.82 | 0.82 | 0.82 | 0.013 | 0.006 | 0.006 | 0.083 | 0.083 | 0.623 |
| 9 | 44.3 | 44.3 | 44.3 | 0.84 | 0.84 | 0.84 | 0.011 | 0.005 | 0.005 | 0.133 | 0.133 | 1.000 |
| 10 | 48.3 | 48.3 | – | 0.86 | 0.86 | – | 0.009 | 0.004 | 0.004 | 0.205 | 0.205 | – |
| 11 | 52.4 | 52.4 | – | 0.88 | 0.88 | – | 0.008 | 0.003 | – | 0.308 | 0.308 | – |
| 12 | 56.3 | 56.3 | – | 0.89 | 0.89 | – | 0.007 | 0.003 | – | 0.442 | 0.442 | – |
| 13 | 59.8 | 59.8 | – | 0.90 | 0.90 | – | 0.006 | 0.003 | – | 0.597 | 0.597 | – |
| 14 | 63.4 | 63.4 | – | 0.91 | 0.91 | – | 0.005 | 0.002 | – | 0.800 | 0.800 | – |
| 15 | 66.3 | 66.3 | – | 0.91 | 0.91 | – | 0.005 | 0.002 | – | 1.000 | 1.000 | – |

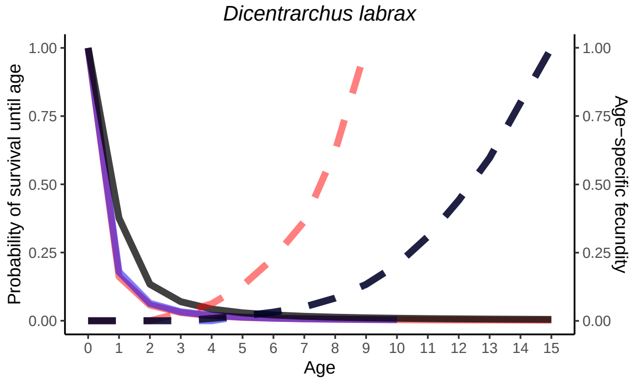

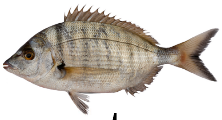

### Sharp-snout seabream

#### *Diplodus puntazzo*

| Parameter | Combined | Female | Male | Ref |
|---|---|---|---|---|
| $L_{inf}$ | 54.1 | 52.3 | 52.7 | Domínguez-Seoane et al.[8] |
| $K$ | 0.182 | 0.203 | 0.187 | |
| $t_0$ | -2.531 | -2.225 | -2.761 | |
| Maturity | 2 | 2 | 2 | |
| Max age | 10 | 10 | 10 | |
| $F = y(L)$ | Power | | | Taieb et al.[26] |
| $\alpha$ | 40.269 | | | |
| $\beta$ | 2.0483 | | | |

| Age | $L_x$ | $L_{x,f}$ | $L_{x,m}$ | $l_x$ | $l_{x,f}$ | $l_{x,m}$ | $Cum_c$ | $Cum_f$ | $Cum_m$ | $B_x$ | $B_{x,f}$ | $B_{x,m}$ |
|---|---|---|---|---|---|---|---|---|---|---|---|---|
| 1 | 25.3 | 25.3 | 25.3 | 0.57 | 0.55 | 0.57 | 0.359 | 0.338 | 0.364 | 0.000 | 0.000 | 0.000 |
| 2 | 31.0 | 31.0 | 31.0 | 0.66 | 0.64 | 0.66 | 0.203 | 0.185 | 0.207 | 0.000 | 0.000 | 0.000 |
| 3 | 34.6 | 34.6 | 34.6 | 0.70 | 0.69 | 0.70 | 0.134 | 0.118 | 0.137 | 0.454 | 0.454 | 0.454 |
| 4 | 37.5 | 37.5 | 37.5 | 0.73 | 0.72 | 0.73 | 0.094 | 0.081 | 0.096 | 0.535 | 0.535 | 0.535 |
| 5 | 39.1 | 39.1 | 39.1 | 0.74 | 0.73 | 0.75 | 0.068 | 0.058 | 0.071 | 0.583 | 0.583 | 0.583 |
| 6 | 43.5 | 43.5 | 43.5 | 0.78 | 0.77 | 0.78 | 0.051 | 0.042 | 0.053 | 0.725 | 0.725 | 0.725 |
| 7 | 45.6 | 45.6 | 45.6 | 0.79 | 0.78 | 0.79 | 0.039 | 0.032 | 0.041 | 0.798 | 0.798 | 0.798 |
| 8 | 48.4 | 48.4 | 48.4 | 0.81 | 0.80 | 0.81 | 0.031 | 0.025 | 0.033 | 0.902 | 0.902 | 0.902 |
| 9 | 49.8 | 49.8 | 49.8 | 0.81 | 0.80 | 0.82 | 0.025 | 0.020 | 0.026 | 0.956 | 0.956 | 0.956 |
| 10 | 50.9 | 50.9 | 50.9 | 0.82 | 0.81 | 0.82 | 0.020 | 0.016 | 0.021 | 1.000 | 1.000 | 1.000 |

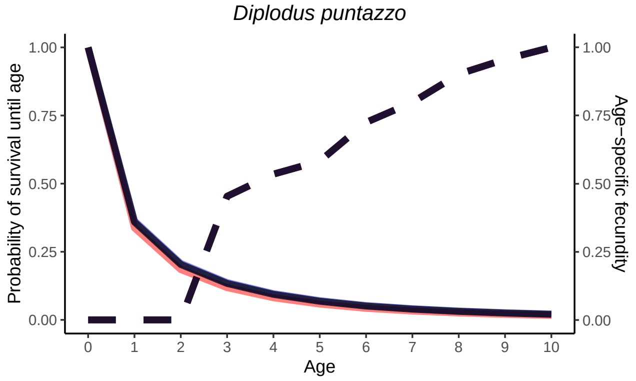

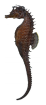

### Long-snouted seahorse

#### *Hippocampus guttulatus*

| Parameter | Combined | Female | Male | Ref |
|---|---|---|---|---|
| $L_{inf}$ | 19.76 | 19.76 | 19.76 | |
| $K$ | 0.571 | 0.571 | 0.571 | |
| $t_0$ | -0.05 | -0.083 | -0.044 | Curtis and Vincent[7] |
| Maturity | 1 | 1 | 1 | |
| Max age | 5 | 5 | 5 | |
| $F = y(L)$ | Exponential | | | |
| $\alpha$ | 78.54 | | | Curtis and Vincent[7] |
| $\beta$ | 0.16 | | | |

| Age | $L_x$ | $L_{x,f}$ | $L_{x,m}$ | $l_x$ | $l_{x,f}$ | $l_{x,m}$ | $Cum_c$ | $Cum_f$ | $Cum_m$ | $B_x$ | $B_{x,f}$ | $B_{x,m}$ |
|---|---|---|---|---|---|---|---|---|---|---|---|---|
| 1 | 13.5 | 13.5 | 13.5 | 0.36 | 0.36 | 0.36 | 0.253 | 0.253 | 0.253 | 0.415 | 0.415 | 0.415 |
| 2 | 16.5 | 16.5 | 16.5 | 0.47 | 0.47 | 0.47 | 0.092 | 0.092 | 0.092 | 0.670 | 0.670 | 0.670 |
| 3 | 18.0 | 18.0 | 18.0 | 0.52 | 0.52 | 0.52 | 0.044 | 0.044 | 0.044 | 0.852 | 0.852 | 0.852 |
| 4 | 18.5 | 18.5 | 18.5 | 0.53 | 0.53 | 0.53 | 0.023 | 0.023 | 0.023 | 0.923 | 0.923 | 0.923 |
| 5 | 19.0 | 19.0 | 19.0 | 0.55 | 0.55 | 0.55 | 0.012 | 0.012 | 0.012 | 1.000 | 1.000 | 1.000 |

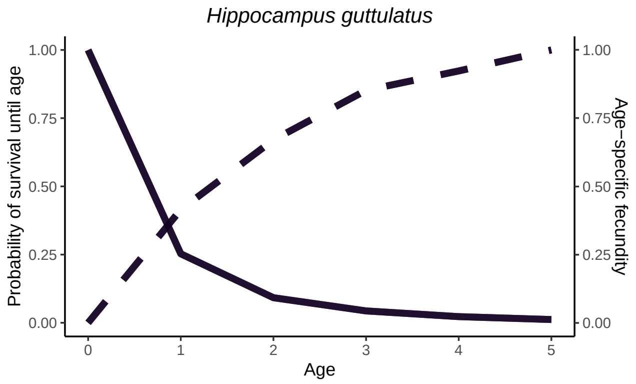

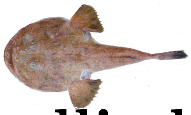

### Blackbellied angler

#### *Lophius budegassa*

| Parameter | Combined | Female | Male | Ref |
|---|---|---|---|---|
| $L_{inf}$ | 102 | 147.3 | 102.5 | García-Rodríguez et al.[11] |
| $K$ | 0.15 | 0.091 | 0.189 | |
| $t_0$ | -0.05 | -0.083 | -0.044 | |
| Maturity | 7 | 6 | 8 | |
| Max age | 21 | 21 | 13 | |
| $F = y(L)$ | Linear | | | Colmenero et al.[6] |
| $\alpha$ | -694487 | | | |
| $\beta$ | 16422 | | | |

| Age | $L_x$ | $L_{x,f}$ | $L_{x,m}$ | $l_x$ | $l_{x,f}$ | $l_{x,m}$ | $Cum_c$ | $Cum_f$ | $Cum_m$ | $B_x$ | $B_{x,f}$ | $B_{x,m}$ |
|---|---|---|---|---|---|---|---|---|---|---|---|---|
| 1 | 21.12 | 25.44 | 32.85 | 0.20 | 0.28 | 0.35 | 1.000 | 1.000 | 1.000 | 0.000 | 0.000 | 0.000 |
| 2 | 32.38 | 36.03 | 44.84 | 0.43 | 0.47 | 0.52 | 0.204 | 0.281 | 0.353 | 0.000 | 0.000 | 0.000 |
| 3 | 42.08 | 45.71 | 54.77 | 0.57 | 0.59 | 0.62 | 0.088 | 0.133 | 0.184 | 0.000 | 0.000 | 0.000 |
| 4 | 50.43 | 54.55 | 62.99 | 0.65 | 0.67 | 0.68 | 0.050 | 0.078 | 0.113 | 0.000 | 0.000 | 0.000 |
| 5 | 57.61 | 62.62 | 69.79 | 0.70 | 0.72 | 0.71 | 0.032 | 0.052 | 0.076 | 0.000 | 0.000 | 0.000 |
| 6 | 63.79 | 69.98 | 75.43 | 0.74 | 0.76 | 0.74 | 0.023 | 0.038 | 0.055 | 0.000 | 0.000 | 0.000 |
| 7 | 69.12 | 76.71 | 80.09 | 0.76 | 0.78 | 0.76 | 0.017 | 0.029 | 0.040 | 0.000 | 0.000 | 0.645 |
| 8 | 73.70 | 82.85 | 83.95 | 0.78 | 0.81 | 0.77 | 0.013 | 0.022 | 0.031 | 0.564 | 0.000 | 0.711 |
| 9 | 77.64 | 88.45 | 87.14 | 0.80 | 0.82 | 0.79 | 0.010 | 0.018 | 0.024 | 0.635 | 0.541 | 0.765 |
| 10 | 81.03 | 93.57 | 89.79 | 0.81 | 0.84 | 0.79 | 0.008 | 0.015 | 0.019 | 0.696 | 0.601 | 0.810 |
| 11 | 83.95 | 98.25 | 91.98 | 0.82 | 0.85 | 0.80 | 0.007 | 0.012 | 0.015 | 0.748 | 0.656 | 0.848 |
| 12 | 86.47 | 102.51 | 93.79 | 0.83 | 0.85 | 0.81 | 0.005 | 0.010 | 0.012 | 0.793 | 0.706 | 0.879 |
| 13 | 88.63 | 106.41 | 95.29 | 0.83 | 0.86 | 0.81 | 0.004 | 0.009 | 0.010 | 0.832 | 0.752 | 0.904 |
| 14 | 90.49 | 109.97 | 96.53 | 0.84 | 0.87 | 0.81 | 0.004 | 0.008 | 0.008 | 0.866 | 0.794 | 0.925 |
| 15 | 92.10 | 113.21 | 97.56 | 0.84 | 0.87 | 0.82 | 0.003 | 0.007 | 0.006 | 0.895 | 0.832 | 0.943 |
| 16 | 93.48 | 116.18 | 98.41 | 0.84 | 0.88 | 0.82 | 0.003 | 0.006 | 0.005 | 0.919 | 0.867 | 0.957 |
| 17 | 94.66 | 118.88 | 99.11 | 0.85 | 0.88 | 0.82 | 0.002 | 0.005 | 0.004 | 0.941 | 0.898 | 0.969 |
| 18 | 95.68 | 121.36 | 99.70 | 0.85 | 0.89 | 0.82 | 0.002 | 0.005 | 0.003 | 0.959 | 0.927 | 0.979 |
| 19 | 96.56 | 123.61 | 100.18 | 0.85 | 0.89 | 0.82 | 0.002 | 0.004 | 0.003 | 0.975 | 0.954 | 0.988 |
| 20 | 97.32 | 125.67 | 100.58 | 0.85 | 0.89 | 0.82 | 0.001 | 0.004 | 0.002 | 0.988 | 0.978 | 0.994 |
| 21 | 97.97 | 127.55 | 100.91 | 0.85 | 0.89 | 0.82 | 0.001 | 0.003 | 0.002 | 1.000 | 1.000 | 1.000 |

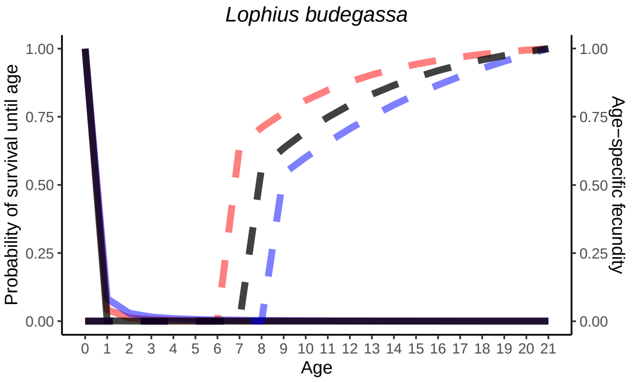

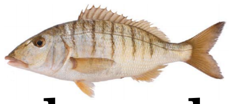

### Striped seabream

### *Lithognathus mormyrus*

| Parameter | Combined | Female | Male | Ref |
|---|---|---|---|---|
| $L_{inf}$ | 35.3 | 35.3 | 35.3 | Monteiro et al.[17] |
| $K$ | 0.264 | 0.264 | 0.264 | |
| $t_0$ | -0.809 | -0.809 | -0.809 | |
| Maturity | 2 | 2 | 2 | |
| Max age | 12 | 12 | 9 | |
| $F = y(L)$ | Power | | | Faraj et al.[9] |
| $\alpha$ | 0.0026 | | | |
| $\beta$ | 4.94 | | | |

| Age | $L_x$ | $L_{x,f}$ | $L_{x,m}$ | $l_x$ | $l_{x,f}$ | $l_{x,m}$ | $Cum_c$ | $Cum_f$ | $Cum_m$ | $B_x$ | $B_{x,f}$ | $B_{x,m}$ |
|---|---|---|---|---|---|---|---|---|---|---|---|---|
| 1 | 13.3 | 16.3 | 15.5 | 0.32 | 0.43 | 0.40 | 1.000 | 1.000 | 1.000 | 0.000 | 0.000 | 0.000 |
| 2 | 19.1 | 19.3 | 19.3 | 0.52 | 0.52 | 0.52 | 0.319 | 0.431 | 0.404 | 0.000 | 0.000 | 0.000 |
| 3 | 23.0 | 23.2 | 22.9 | 0.61 | 0.61 | 0.60 | 0.165 | 0.224 | 0.210 | 0.051 | 0.053 | 0.148 |
| 4 | 24.9 | 24.8 | 24.9 | 0.64 | 0.64 | 0.64 | 0.100 | 0.137 | 0.127 | 0.076 | 0.074 | 0.224 |
| 5 | 26.1 | 26.0 | 26.1 | 0.66 | 0.66 | 0.66 | 0.064 | 0.087 | 0.081 | 0.095 | 0.094 | 0.283 |
| 6 | 28.1 | 28.1 | 28.4 | 0.69 | 0.69 | 0.69 | 0.042 | 0.058 | 0.054 | 0.137 | 0.137 | 0.429 |
| 7 | 30.2 | 30.5 | 29.8 | 0.72 | 0.72 | 0.71 | 0.029 | 0.040 | 0.037 | 0.196 | 0.206 | 0.545 |
| 8 | 32.0 | 31.9 | 32.4 | 0.74 | 0.74 | 0.74 | 0.021 | 0.029 | 0.026 | 0.261 | 0.257 | 0.823 |
| 9 | 33.9 | 34.1 | 33.7 | 0.76 | 0.76 | 0.75 | 0.015 | 0.021 | 0.020 | 0.347 | 0.357 | 1.000 |
| 10 | 34.8 | 34.4 | – | 0.76 | 0.76 | – | 0.012 | 0.016 | 0.015 | 0.395 | 0.373 | – |
| 11 | 32.5 | 35.2 | – | 0.74 | 0.74 | – | 0.009 | 0.012 | – | 0.282 | 0.282 | – |
| 12 | 42.0 | 42.0 | – | 0.82 | 0.82 | – | 0.007 | 0.009 | – | 1.000 | 1.000 | – |

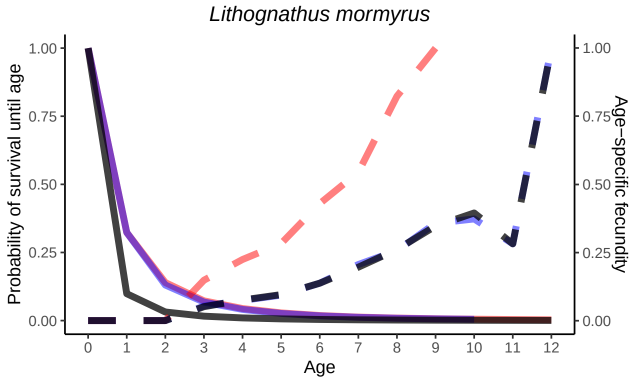

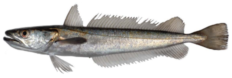

### European hake

#### *Merluccius merluccius*

| Parameter | Combined | Female | Male | Ref |
|---|---|---|---|---|
| $L_{inf}$ | 88.7 | 88.0 | 70 | Piñeiro and Saínza[22] |
| $K$ | 0.128 | 0.127 | 0.184 | |
| $t_0$ | -1.174 | -1.157 | -0.973 | |
| Maturity | 3 | 3 | 4 | |
| Max age | 11 | 11 | 9 | |
| $F = y(L)$ | Power | | | Biagi et al.[3] |
| $\alpha$ | 2.54 | | | |
| $\beta$ | 3.07 | | | |

| Age | $L_x$ | $L_{x,f}$ | $L_{x,m}$ | $l_x$ | $l_{x,f}$ | $l_{x,m}$ | $Cum_c$ | $Cum_f$ | $Cum_m$ | $B_x$ | $B_{x,f}$ | $B_{x,m}$ |
|---|---|---|---|---|---|---|---|---|---|---|---|---|
| 1 | 20.6 | 20.9 | 21.0 | 0.32 | 0.33 | 0.33 | 1.000 | 1.000 | 1.000 | 0.000 | 0.000 | 0.000 |
| 2 | 29.0 | 28.4 | 29.5 | 0.50 | 0.50 | 0.51 | 0.319 | 0.334 | 0.326 | 0.000 | 0.000 | 0.000 |
| 3 | 36.7 | 37.0 | 36.4 | 0.62 | 0.63 | 0.61 | 0.161 | 0.167 | 0.167 | 0.000 | 0.000 | 0.000 |
| 4 | 43.8 | 44.5 | 42.7 | 0.69 | 0.70 | 0.68 | 0.099 | 0.105 | 0.102 | 0.192 | 0.000 | 0.352 |
| 5 | 50.0 | 48.7 | 45.7 | 0.74 | 0.73 | 0.71 | 0.069 | 0.074 | 0.069 | 0.288 | 0.235 | 0.434 |
| 6 | 55.4 | 53.7 | 49.7 | 0.77 | 0.77 | 0.74 | 0.051 | 0.054 | 0.049 | 0.395 | 0.318 | 0.561 |
| 7 | 58.3 | 56.4 | 54.2 | 0.79 | 0.78 | 0.76 | 0.039 | 0.041 | 0.036 | 0.461 | 0.370 | 0.732 |
| 8 | 63.1 | 62.3 | 56.6 | 0.81 | 0.81 | 0.78 | 0.031 | 0.032 | 0.027 | 0.588 | 0.502 | 0.836 |
| 9 | 67.1 | 68.7 | 60.0 | 0.82 | 0.83 | 0.79 | 0.025 | 0.026 | 0.021 | 0.711 | 0.677 | 1.000 |
| 10 | 75.0 | 75.0 | – | 0.85 | 0.85 | – | 0.020 | 0.022 | 0.017 | 1.000 | 0.887 | – |
| 11 | 74.0 | 78.0 | – | 0.85 | 0.86 | – | 0.017 | 0.018 | – | 0.960 | 1.000 | – |

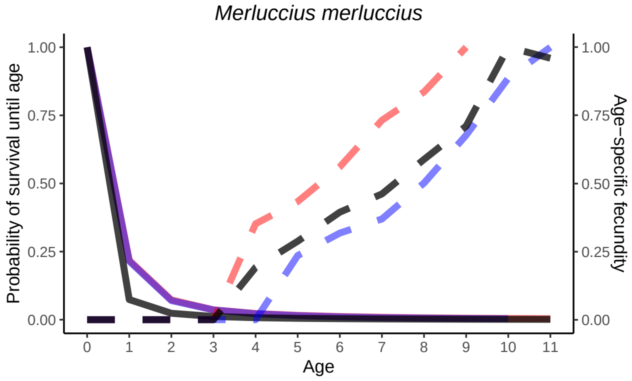

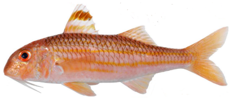

### Striped red mullet
### *Mullus surmuletus*

| Parameter | Combined | Female | Male | Ref |
|---|---|---|---|---|
| $L_{inf}$ | 31.28 | 31.9 | 25.54 | Reñones et al.[23] |
| $K$ | 0.211 | 0.205 | 0.273 | |
| $t_0$ | -2.348 | -2.605 | -2.45 | |
| Maturity | 2 | 1 | 2 | |
| Max age | 6 | 6 | 6 | |
| $F = y(L)$ | Power | | | Amin et al.[2] |
| $\alpha$ | 0.0255 | | | |
| $\beta$ | 5.031 | | | |

| Age | $L_x$ | $L_{x,f}$ | $L_{x,m}$ | $l_x$ | $l_{x,f}$ | $l_{x,m}$ | $Cum_c$ | $Cum_f$ | $Cum_m$ | $B_x$ | $B_{x,f}$ | $B_{x,m}$ |
|---|---|---|---|---|---|---|---|---|---|---|---|---|
| 1 | 15.9 | 16.7 | 15.7 | 0.56 | 0.58 | 0.57 | 1.000 | 1.000 | 1.000 | 0.000 | 0.000 | 0.146 |
| 2 | 18.6 | 19.5 | 17.8 | 0.63 | 0.65 | 0.62 | 0.559 | 0.582 | 0.567 | 0.000 | 0.000 | 0.272 |
| 3 | 21.3 | 21.9 | 19.8 | 0.69 | 0.70 | 0.67 | 0.352 | 0.379 | 0.354 | 0.203 | 0.157 | 0.465 |
| 4 | 23.1 | 23.6 | 21.5 | 0.72 | 0.72 | 0.70 | 0.242 | 0.264 | 0.237 | 0.304 | 0.227 | 0.717 |
| 5 | 24.6 | 25.3 | 22.0 | 0.74 | 0.75 | 0.71 | 0.174 | 0.191 | 0.167 | 0.419 | 0.320 | 0.800 |
| 6 | 29.3 | 31.7 | 23.0 | 0.79 | 0.81 | 0.73 | 0.128 | 0.143 | 0.118 | 1.000 | 1.000 | 1.000 |

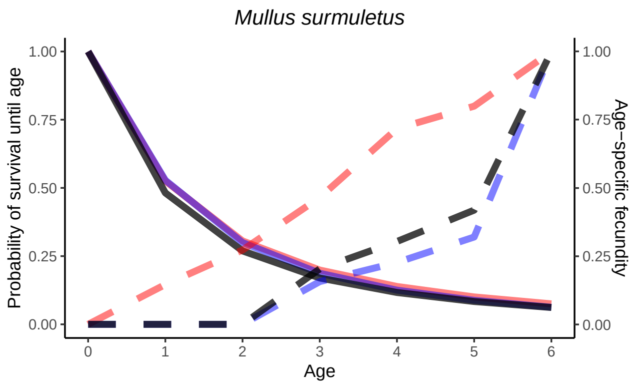

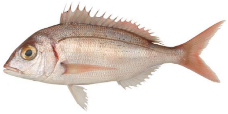

### Common pandora

#### *Pagellus erythrinus*

| Parameter | Combined | Female | Male | Ref |
|---|---|---|---|---|
| $L_{inf}$ | 38.29 | 35.41 | 40.01 | Yapici and Filiz[29] |
| $K$ | 0.148 | 0.17 | 0.135 | |
| $t_0$ | -1.42 | -1.32 | -1.54 | |
| Maturity | 2 | 1 | 2 | |
| Max age | 8 | 8 | 8 | |
| $F = y(L)$ | Power | | | Papaconstantinou et al.[21] |
| $\alpha$ | 1 | | | |
| $\beta$ | 3.74 | | | |

| Age | $L_x$ | $L_{x,f}$ | $L_{x,m}$ | $l_x$ | $l_{x,f}$ | $l_{x,m}$ | $Cum_c$ | $Cum_f$ | $Cum_m$ | $B_x$ | $B_{x,f}$ | $B_{x,m}$ |
|---|---|---|---|---|---|---|---|---|---|---|---|---|
| 1 | 11.66 | 11.66 | 11.66 | 0.41 | 0.41 | 0.42 | 1.000 | 1.000 | 1.000 | 0.000 | 0.000 | 0.034 |
| 2 | 15.27 | 15.27 | 15.27 | 0.56 | 0.55 | 0.56 | 0.414 | 0.407 | 0.424 | 0.000 | 0.000 | 0.093 |
| 3 | 18.19 | 18.19 | 18.19 | 0.64 | 0.63 | 0.64 | 0.230 | 0.223 | 0.239 | 0.179 | 0.179 | 0.179 |
| 4 | 20.96 | 20.96 | 20.96 | 0.69 | 0.69 | 0.70 | 0.147 | 0.141 | 0.154 | 0.304 | 0.304 | 0.304 |
| 5 | 24.07 | 24.07 | 24.07 | 0.74 | 0.74 | 0.75 | 0.102 | 0.097 | 0.108 | 0.511 | 0.511 | 0.511 |
| 6 | 25.89 | 25.89 | 25.89 | 0.77 | 0.76 | 0.77 | 0.076 | 0.071 | 0.081 | 0.671 | 0.671 | 0.671 |
| 7 | 27.19 | 27.19 | 27.19 | 0.78 | 0.78 | 0.79 | 0.058 | 0.054 | 0.062 | 0.805 | 0.805 | 0.805 |
| 8 | 28.81 | 28.81 | 28.81 | 0.80 | 0.79 | 0.80 | 0.045 | 0.042 | 0.049 | 1.000 | 1.000 | 1.000 |

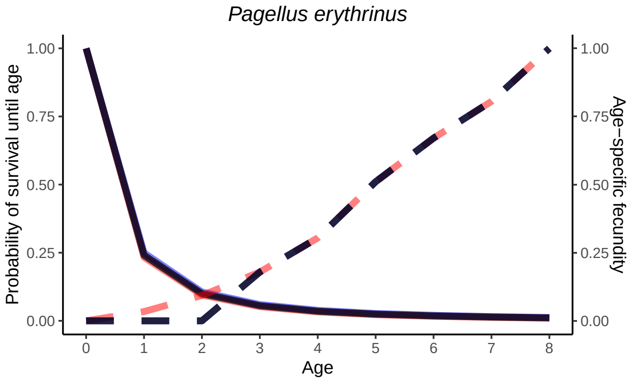

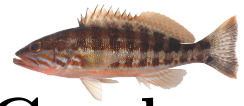

### Comber

#### *Serranus cabrilla*

| Parameter | Combined | Female | Male | Ref |
|---|---|---|---|---|
| $L_{inf}$ | 23.88 | 23.88 | 23.88 | Uçkun İlhan et al.[27] |
| $K$ | 0.298 | 0.298 | 0.298 | |
| $t_0$ | -1.577 | -1.577 | -1.577 | |
| Maturity | 2 | 2 | 2 | |
| Max age | 6 | 6 | 6 | |
| $F = y(L)$ | Exponential | | | Palacios Sartagal[20] |
| $\alpha$ | 72.46 | | | |
| $\beta$ | 0.22 | | | |

| Age | $L_x$ | $L_{x,f}$ | $L_{x,m}$ | $l_x$ | $l_{x,f}$ | $l_{x,m}$ | $Cum_c$ | $Cum_f$ | $Cum_m$ | $B_x$ | $B_{x,f}$ | $B_{x,m}$ |
|---|---|---|---|---|---|---|---|---|---|---|---|---|
| 1 | 12.79 | 12.79 | 12.79 | 0.47 | 0.47 | 0.47 | 1.000 | 1.000 | 1.000 | 0.000 | 0.000 | 0.000 |
| 2 | 15.84 | 15.84 | 15.84 | 0.58 | 0.58 | 0.58 | 0.468 | 0.468 | 0.468 | 0.000 | 0.000 | 0.000 |
| 3 | 17.61 | 17.61 | 17.61 | 0.62 | 0.62 | 0.62 | 0.269 | 0.269 | 0.269 | 0.432 | 0.432 | 0.432 |
| 4 | 19.20 | 19.20 | 19.20 | 0.66 | 0.66 | 0.66 | 0.168 | 0.168 | 0.168 | 0.614 | 0.614 | 0.614 |
| 5 | 20.61 | 20.61 | 20.61 | 0.69 | 0.69 | 0.69 | 0.111 | 0.111 | 0.111 | 0.837 | 0.837 | 0.837 |
| 6 | 21.42 | 21.42 | 21.42 | 0.70 | 0.70 | 0.70 | 0.077 | 0.077 | 0.077 | 1.000 | 1.000 | 1.000 |

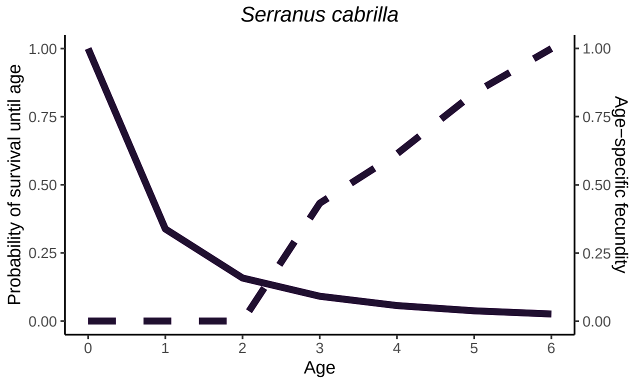

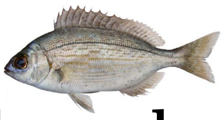

### Black seabream

#### *Spondyliosoma cantharus*

| Parameter | Combined | Female | Male | Ref |
|---|---|---|---|---|
| $L_{inf}$ | 43.35 | 41.92 | 45.89 | Pajuelo and Lorenzo[19] |
| $K$ | 0.24 | 0.25 | 0.2 | |
| $t_0$ | -0.11 | -0.29 | -078 | |
| Maturity | 2 | 3 | 2 | |
| Max age | 10 | 10 | 10 | |
| $F = y(L)$ | Power | | | Gonçalves and Erzini[12] |
| $\alpha$ | 436.27 | | | |
| $\beta$ | 1.5747 | | | |

| Age | $L_x$ | $L_{x,f}$ | $L_{x,m}$ | $l_x$ | $l_{x,f}$ | $l_{x,m}$ | $Cum_c$ | $Cum_f$ | $Cum_m$ | $B_x$ | $B_{x,f}$ | $B_{x,m}$ |
|---|---|---|---|---|---|---|---|---|---|---|---|---|
| 1 | 10.6 | 13.75 | 10.14 | 0.14 | 0.26 | 0.15 | 1.000 | 1.000 | 1.000 | 0.000 | 0.000 | 0.000 |
| 2 | 17.3 | 19.57 | 17.22 | 0.39 | 0.46 | 0.42 | 0.137 | 0.264 | 0.146 | 0.000 | 0.000 | 0.000 |
| 3 | 23.5 | 24.34 | 22.80 | 0.55 | 0.57 | 0.56 | 0.053 | 0.121 | 0.061 | 0.443 | 0.447 | 0.000 |
| 4 | 27.9 | 28.25 | 27.18 | 0.63 | 0.64 | 0.64 | 0.029 | 0.069 | 0.035 | 0.581 | 0.565 | 0.555 |
| 5 | 31.4 | 31.45 | 30.63 | 0.68 | 0.68 | 0.69 | 0.018 | 0.044 | 0.022 | 0.699 | 0.669 | 0.669 |
| 6 | 34.5 | 34.06 | 33.35 | 0.71 | 0.71 | 0.72 | 0.012 | 0.030 | 0.015 | 0.811 | 0.759 | 0.765 |
| 7 | 36.5 | 36.21 | 35.48 | 0.73 | 0.73 | 0.75 | 0.009 | 0.021 | 0.011 | 0.887 | 0.836 | 0.844 |
| 8 | 38.2 | 37.96 | 37.61 | 0.75 | 0.75 | 0.76 | 0.006 | 0.015 | 0.008 | 0.952 | 0.900 | 0.908 |
| 9 | 38.9 | 39.40 | 38.48 | 0.75 | 0.76 | 0.77 | 0.005 | 0.012 | 0.006 | 0.980 | 0.955 | 0.959 |
| 10 | 39.4 | 40.58 | 39.52 | 0.76 | 0.77 | 0.78 | 0.004 | 0.009 | 0.005 | 1.000 | 1.000 | 1.000 |

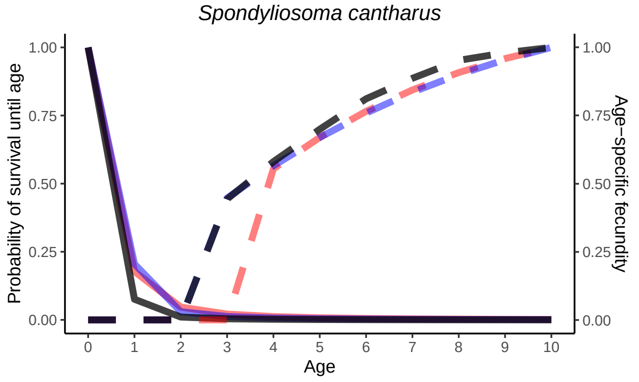

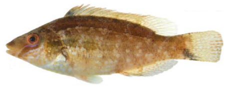

### Grey wrasse

#### *Symphodus cinereus*

| Parameter | Combined | Female | Male | Ref |
|---|---|---|---|---|
| $L_{inf}$ | 10.61 | 9.6 | 11.62 | Kara and Quignard[15] |
| $K$ | 0.483 | 0.483 | 0.485 | |
| $t_0$ | -0.74 | -0.74 | 0.44 | |
| Maturity | 1 | 1 | 1 | |
| Max age | 6 | 6 | 6 | |
| $F = y(L)$ | Exponential | | | Kara and Quignard[15] |
| $\alpha$ | 2659 | | | |
| $\beta$ | 0.0619 | | | |

| Age | $L_x$ | $L_{x,f}$ | $L_{x,m}$ | $l_x$ | $l_{x,f}$ | $l_{x,m}$ | $Cum_c$ | $Cum_f$ | $Cum_m$ | $B_x$ | $B_{x,f}$ | $B_{x,m}$ |
|---|---|---|---|---|---|---|---|---|---|---|---|---|
| 1 | 7.79 | 7.04 | 6.17 | 0.46 | 0.46 | 0.29 | 1.000 | 1.000 | 1.000 | 0.853 | 0.866 | 0.735 |
| 2 | 8.87 | 8.02 | 8.26 | 0.53 | 0.53 | 0.45 | 0.464 | 0.463 | 0.286 | 0.912 | 0.920 | 0.837 |
| 3 | 9.53 | 8.63 | 9.55 | 0.57 | 0.57 | 0.52 | 0.247 | 0.246 | 0.127 | 0.950 | 0.955 | 0.906 |
| 4 | 9.95 | 9.00 | 10.35 | 0.59 | 0.59 | 0.56 | 0.140 | 0.140 | 0.066 | 0.975 | 0.977 | 0.952 |
| 5 | 10.20 | 9.23 | 10.84 | 0.60 | 0.60 | 0.58 | 0.082 | 0.082 | 0.037 | 0.990 | 0.991 | 0.982 |
| 6 | 10.36 | 9.37 | 11.14 | 0.61 | 0.61 | 0.60 | 0.049 | 0.049 | 0.022 | 1.000 | 1.000 | 1.000 |

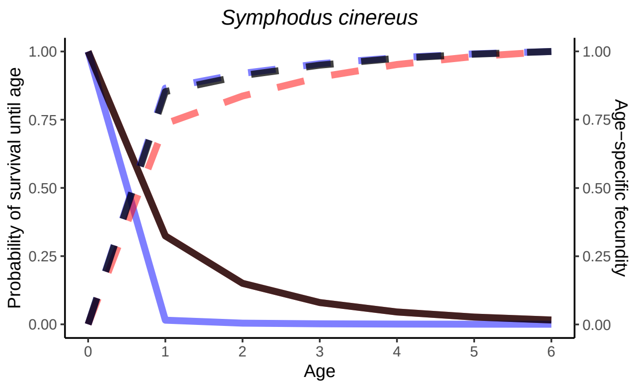

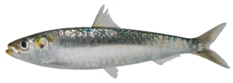

### European pilchard

#### *Sardina pilchardus*

| Parameter | Combined | Female | Male | Ref |
|---|---|---|---|---|
| $L_{inf}$ | 18.02 | 18.02 | 18.02 | Alemany and Alvarez[1] |
| $K$ | 0.65 | 0.65 | 0.65 | |
| $t_0$ | -0.67 | -0.67 | -0.67 | |
| Maturity | 1 | 1 | 1 | |
| Max age | 5 | 5 | 5 | |
| $F = y(L)$ | Power | | | Bouhali et al.[4] |
| $\alpha$ | 6.858 | | | |
| $\beta$ | 2.497 | | | |

| Age | $L_x$ | $L_{x,f}$ | $L_{x,m}$ | $l_x$ | $l_{x,f}$ | $l_{x,m}$ | $Cum_c$ | $Cum_f$ | $Cum_m$ | $B_x$ | $B_{x,f}$ | $B_{x,m}$ |
|---|---|---|---|---|---|---|---|---|---|---|---|---|
| 1 | 14.2 | 14.2 | 14.2 | 0.39 | 0.39 | 0.39 | 1.000 | 1.000 | 1.000 | 0.602 | 0.602 | 0.602 |
| 2 | 16.0 | 16.0 | 16.0 | 0.46 | 0.46 | 0.46 | 0.395 | 0.395 | 0.395 | 0.811 | 0.811 | 0.811 |
| 3 | 17.2 | 17.2 | 17.2 | 0.50 | 0.50 | 0.50 | 0.182 | 0.182 | 0.182 | 0.972 | 0.972 | 0.972 |
| 4 | 17.2 | 17.2 | 17.2 | 0.50 | 0.50 | 0.50 | 0.090 | 0.090 | 0.090 | 0.972 | 0.972 | 0.972 |
| 5 | 17.4 | 17.4 | 17.4 | 0.50 | 0.50 | 0.50 | 0.045 | 0.045 | 0.045 | 1.000 | 1.000 | 1.000 |

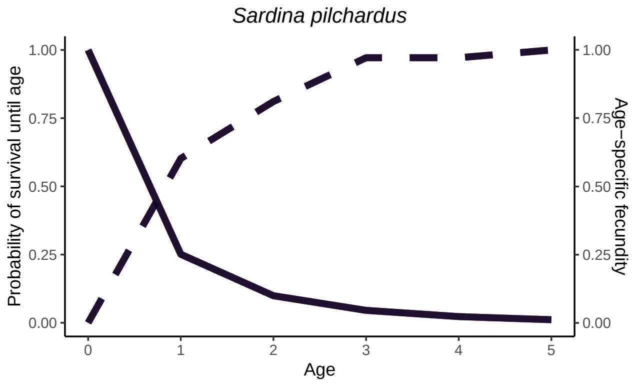

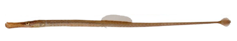

### Broadnosed pipefish

#### *Syngnathus typhle*

| Parameter | Combined | Female | Male | Ref |
|---|---|---|---|---|
| $L_{inf}$ | 26.2 | 26.2 | 26.2 | Froese et al.[10] |
| $K$ | 0.558 | 0.558 | 0.558 | |
| $t_0$ | -0.5 | -0.5 | -0.5 | |
| Maturity | 1 | 1 | 1 | |
| Max age | 3 | 3 | 3 | |
| $F = y(L)$ | Exponential | | | Rispoli and Wilson[24] |
| $\alpha$ | 6.7977 | | | |
| $\beta$ | 0.1522 | | | |

| Age | $L_x$ | $L_{x,f}$ | $L_{x,m}$ | $l_x$ | $l_{x,f}$ | $l_{x,m}$ | $Cum_c$ | $Cum_f$ | $Cum_m$ | $B_x$ | $B_{x,f}$ | $B_{x,m}$ |
|---|---|---|---|---|---|---|---|---|---|---|---|---|
| 1 | 14.86 | 14.86 | 14.86 | 0.27 | 0.27 | 0.27 | 1.000 | 1.000 | 1.000 | 0.314 | 0.314 | 0.314 |
| 2 | 19.71 | 19.71 | 19.71 | 0.43 | 0.43 | 0.43 | 0.271 | 0.271 | 0.271 | 0.656 | 0.656 | 0.656 |
| 3 | 22.48 | 22.48 | 22.48 | 0.50 | 0.50 | 0.50 | 0.115 | 0.115 | 0.115 | 1.000 | 1.000 | 1.000 |

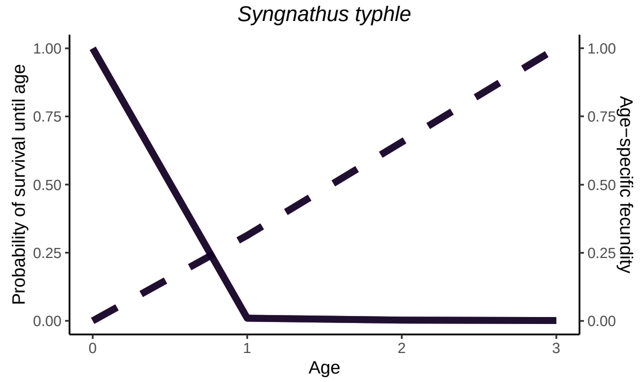

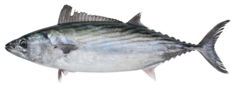

### Atlantic bonito

### *Sarda sarda*

| Parameter | Combined | Female | Male | Ref |
|---|---|---|---|---|
| $L_{inf}$ | 80.87 | 80.87 | 80.87 | Orsi Relini et al.[18] |
| $K$ | 0.352 | 0.352 | 0.352 | |
| $t_0$ | -1.7 | -1.7 | -1.7 | |
| Maturity | 1 | 1 | 1 | |
| Max age | 4 | 4 | 4 | |
| $F = y(L)$ | Power | | | Orsi Relini et al.[18] |
| $\alpha$ | 0.01 | | | |
| $\beta$ | 4.59 | | | |

| Age | $L_x$ | $L_{x,f}$ | $L_{x,m}$ | $l_x$ | $l_{x,f}$ | $l_{x,m}$ | $Cum_c$ | $Cum_f$ | $Cum_m$ | $B_x$ | $B_{x,f}$ | $B_{x,m}$ |
|---|---|---|---|---|---|---|---|---|---|---|---|---|
| 1 | 51.71 | 51.71 | 51.71 | 0.50 | 0.50 | 0.50 | 1.000 | 1.000 | 1.000 | 0.233 | 0.233 | 0.233 |
| 2 | 57.04 | 57.04 | 57.04 | 0.55 | 0.55 | 0.55 | 0.502 | 0.502 | 0.502 | 0.366 | 0.366 | 0.366 |
| 3 | 63.15 | 63.15 | 63.15 | 0.60 | 0.60 | 0.60 | 0.277 | 0.277 | 0.277 | 0.584 | 0.584 | 0.584 |
| 4 | 71.00 | 71.00 | 71.00 | 0.65 | 0.65 | 0.65 | 0.166 | 0.166 | 0.166 | 1.000 | 1.000 | 1.000 |

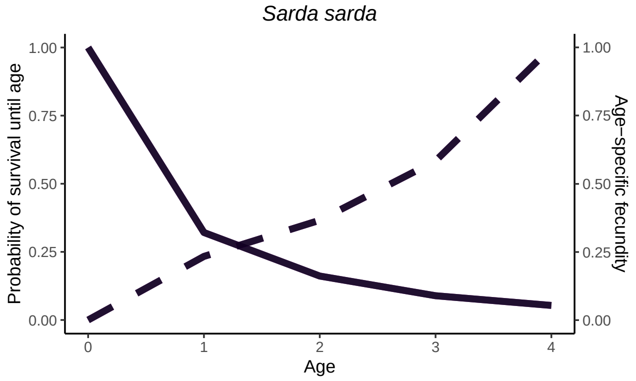

### References


[1] Alemany, F. and Alvarez, F. (1993). Growth differences among sardine (Sardina pilchardus Walb.) populations in Western Mediterranean. *Sci. Mar.*, 57:229–234.

[2] Amin, A., Madkour, F., Abu El-Regal, M., and Moustafa, A. (2016). Reproductive biology of Mullus surmuletus (Linnaeus, 1758) from the Egyptian Mediterranean Sea (Port Said. *iNTERNATIONAL jOURNAL OF eNVIRONMENTAL sCIENCE and engineering*, 7:1–10.

[3] Biagi, F., Sbrana, M., Cesarini, A., and Viva, C. (1995). Reproductive biology and fecundity of Merlucciusmerluccius (Linnaeus, 1758) in the Northern Tyrrhenian sea.Dynamique des populationsmarines. *Cahiers Options Méditerranéennes*, (10):47–48.

[4] Bouhali, F., Lechekhab, S., Ladaimia, S., Assia, B., Amara, R., and Borhane, D. (2015). Reproduction et maturation des gonades de Sardina pilchardus dans le golfe d'Annaba (Nord-Est algérien). *Cybium: international journal of ichthyology.*

[5] Carrassón, M. and Bau, M. (2003). Reproduction, gonad histology and fecundity of *Aidablennius sphynx* (Pisces: Blenniidae) of the Catalan Sea (North-Western Mediterranean). *Scientia Marina*, 67(4):461–469.

[6] Colmenero, A. I., Tuset, V. M., Recasens, L., and Sanchez, P. (2013). Reproductive biology of Black Anglerfish (Lophius budegassa) in the northwestern Mediterranean Sea. *Fishery Bulletin*, 111(4):390–401.

[7] Curtis, J. M. R. and Vincent, A. C. J. (2006). Life history of an unusual marine fish: Survival, growth and movement patterns of Hippocampus guttulatus Cuvier 1829. *Journal of Fish Biology*, 68(3):707–733.

[8] Domínguez-Seoane, R., Pajuelo, J. G., Lorenzo, J. M., and Ramos, A. G. (2006). Age and growth of the sharpsnout seabream Diplodus puntazzo (Cetti, 1777) inhabiting the Canarian archipelago, estimated by reading otoliths and by backcalculation. *Fisheries Research*, 81(2):142–148.

[9] Faraj, E., Alssalam, A., Ali, S., Sayed, M., Sayed, E., Mor, E., Ali, R., Ali, S., Salem, E., and Alfergani, E. (2016). Reproductive Biology of the Striped Seabream Lithognathus mormyrus (Linnaeus, 1758) from Al Haneah Fishing Site, Mediterranean Sea, Eastern Libya. *Journal of Life Sciences*, 10.

[10] Froese, R., Pauly, D., and Editors (2000). FishBase 2000: Concepts, design and data sources. page 344.

[11] García-Rodríguez, M., Pereda, P., Landa, J., and Esteban, A. (2005). On the biology and growth of the anglerfish Lophius budegassa Spinola, 1807 in the Spanish Mediterranean: A preliminary approach. *Fisheries Research*, 71:197–208.

[12] Gonçalves, J. and Erzini, K. (2000). The reproductive biology of Spondyliosoma cantharus (L.) from the SW Coast of Portugal. *Scientia Marina*, 64:403–411.

[13] Iglésias, S. (2013). *Actinopterygians from the North-Eastern Atlantic and the Mediterranean (A Natural Classification Based on Collection Specimens, with DNA Barcodes and Standardized Photographs), Volume I (Plates), Provisional Version 09.*

[14] Kara, M. H. (1997). Cycle sexuel et fécondité du Loup Dicentrarchus labrax (Poisson Moronidé) du golfe d'Annaba. *Cahiers de Biologie Marine*, (3).

[15] Kara, M. H. and Quignard, J.-P. (2018). *Les poissons des lagunes et des estuaires de Méditerranée. 2, 2,.*

[16] Milton, P. (1983). Biology of littoral blenniid fishes on the coast of south-west England. *Journal of the Marine Biological Association of the United Kingdom*, 63(1):223–237.

[17] Monteiro, P., Bentes, L., Coelho, R., Correia, C., Erzini, K., Lino, P. G., Ribeiro, J., and Gonçalves, J. M. S. (2010). Age and growth, mortality and reproduction of the striped sea bream, Lithognathus mormyrus Linnaeus 1758, from the south coast of Portugal (Algarve). *Marine Biology Research*, 6(1):53–65.

[18] Orsi Relini, L., Garibaldi, F., Cima, C., Palandri, G., Lanteri, L., and Relini, M. (2005). Biology of atlantic bonito, Sarda sarda (Bloch, 1793), in the western and central mediterranean. A summary concerning a possible stock unit.

[19] Pajuelo, J. G. and Lorenzo, J. M. (1999). Life History of Black Seabream, Spondyliosoma cantharus, off the Canary Islands, Central-east Atlantic. *Environmental Biology of Fishes*, 54(3):325–336.

[20] Palacios Sartagal, N. (2017). *Estudi de La Fecunditat i Estratègia Reproductiva de Serranus Cabrilla (Pisces, Serranidae)*. PhD thesis, Universitat de Girona.

[21] Papaconstantinou, C., G.Petrakis, and Vassilopoulou, V. (1986). The fecundity of hake (Merluccius merluccius L.) and red pandora (Pagellus erythrinus L.) in the Greek seas. *Acta Adriatica*, 27:85–95.

[22] Piñeiro, C. and Saínza, M. (2003). Age estimation, growth and maturity of the European hake (Merluccius merluccius (Linnaeus, 1758)) from Iberian Atlantic waters. *ICES Journal of Marine Science*, 60(5):1086–1102.

[23] Reñones, O., Massutí, E., and Morales-Nin, B. (1995). Life history of the red mullet Mullus surmuletus from the bottom-trawl fishery off the Island of Majorca (north-west Mediterranean). *Marine Biology*, 123(3):411–419.

[24] Rispoli, V. F. and Wilson, A. B. (2008). Sexual size dimorphism predicts the frequency of multiple mating in the sex-role reversed pipefish Syngnathus typhle. *Journal of Evolutionary Biology*, 21(1):30–38.

[25] Škeljo, F. (2012). *Dinamika Populacije Kneza, Coris Julis (Linnaeus, 1758) u Istočnom Jadranu*. PhD thesis, University of Split.

[26] Taieb, A. H., Ghorbel, M., and Jarboui, O. (2013). Study of fecundity for Diplodus vulgaris (Teleost, Sparidae) in Gulf of Gabes. page 5.

[27] Uçkun İlhan, D., Akalın, S., tosunoğlu, Z., and Ozaydin, O. (2010). Growth Characteristics and Reproduction of Comber, Serranus Cabrilla (Actinopterygii, Perciformes, Serranidae), in the Aegean Sea. *Acta Ichthyologica Et Piscatoria*, 40:55–60.

[28] Wassef, E. A. and Emary, H. (1989). Contribution to the biology of bass, Dicentrarchus labrax L. in the Egyptian Mediterranean waters off Alexandria. /paper/Contribution-to-the-biology-of-bass%2C-Dicentrarchus-Wassef-Emary/335360d91c9f3d088051211b1da6b92b41cfb5fa.

[29] Yapici, S. and Filiz, H. (2019). Biological aspects of two coexisting native and non-native fish species in the Aegean Sea: Pagellus erythrinus vs. Nemipterus randalli. *Mediterranean Marine Science*, 20(3):594–602.
